# Expandable DNA-origami rings as a nanomechanical platform for studying disordered nucleoporins

**DOI:** 10.64898/2026.09.22.753628

**Authors:** Eason Cao, Christopher Maffeo, Lukas Beckert, Larisa E. Kapinos, Ryan J. Malonis, Daniel Son, Kun Zhou, Zhuxi Wang, Qingzhou Feng, Sunandini Chandra, Qi Shen, Yong Xiong, C. Patrick Lusk, Roderick Y. H. Lim, Aleksei Aksimentiev, Chenxiang Lin

## Abstract

Phe-Gly-rich nucleoporins (FG-Nups) within the nuclear pore complex (NPC) form a diffusion barrier that controls the exchange of macromolecules between the nucleus and cytoplasm. The behaviors of these intrinsically disordered FG-Nups are fundamental to the NPC function but remain difficult to probe within the dynamic, native nuclear pore. To bridge this knowledge gap, here we build expandable DNA-origami rings with tunable elasticity and study FG-Nups site-specifically tethered in such mechanically responsive NPC mimics. The elastic deformation of DNA rings, measured by single-particle TEM, reveals the type (attraction or repulsion) and strength of FG-Nup interactions that shape the collective morphology of these unstructured proteins. Analyzing homotypic interactions of five FG-Nup species finds diverse FG-Nup behaviors that can be classified as repulsive/extended (Nsp1), moderately cohesive/extended (Nup62, Nup153, and Nup214), and strongly cohesive/compact (Nup98). Furthermore, we examine how FG-Nups’ primary sequence, *O*-glycosylation, grafting density, and binding to nuclear transport receptors (NTRs) impact their biophysical state. For example, *O*-GlcNAcylation reduced FG-Nup cohesiveness, switching Nup98 from a collapsed state to more extended conformations. Our work thus elucidates spatially confined Nup–Nup and Nup–NTR interactions within an NPC-like nanopore, suggests a previously underappreciated role of FG-Nups in modulating NPC dilation, and establishes a generalizable method for studying multivalent interactions among disordered proteins.

## Introduction

The nuclear pore complex (NPC) is the sole gateway for macromolecular exchange between the nucleus and cytoplasm^1,2^. Each NPC is a ∼50 (yeast) to ∼120 MDa (human) assembly composed of ∼30 distinct nucleoporins (Nups), arranged with eight-fold symmetry to form a conduit spanning the nuclear envelope (NE)^3,4^. About a dozen types of Nups containing intrinsically disordered Phe-Gly-rich domains (FG-Nups) line the entire pathway through the NPC, mainly inside its ∼45–70 nm wide central channel, creating a permeability barrier to curb the diffusion of inert macromolecules while supporting the rapid, nuclear transport receptor (NTR)-mediated cargo translocation^5,6^.

How NPCs form a selective barrier has been a focal point of research for decades, spawning several models that all recognize the central roles of highly concentrated FG-Nups in forming a diffusion barrier and NTRs in remodeling the FG network and licensing the passage of otherwise impermissible cargos. These models differ in their depiction of the FG-Nup assemblies as a sieve-like hydrogel^7–10^, liquid-like condensate^11–13^ or brush-like filaments^14–17^, thus suggesting different (but not mutually exclusive) physicochemical underpinnings of the NPC’s function: the cohesive interactions or the entropic dynamics of the unstructured FG domains. Reconstituted systems are instrumental in conceptualizing these models: phase-separated FG repeats, including macroscopic hydrogels and micrometer-sized condensates, and surface-tethered FG domains evidently recapitulate the NPC’s basic barrier-transporter function^7,8,11,13,17^. Such models are further enabled by biochemical and biophysical characterizations of purified FG-Nups, including their material properties as biopolymers and their cohesive/repulsive interactions^6,7,15,18^. However, recent work using advanced fluorescence cell imaging, *in situ* atomic force microscopy (AFM) and molecular dynamics (MD) simulation revealed unique protein dynamics within the NPC that are not represented by bulk condensates or gels formed from unconstrained FG-Nups *in vitro*^13,19–21^. These findings highlight the joint importance of FG-Nups’ intrinsic properties, including charge and hydrophobicity as dictated by their amino acid sequence and post-translational modifications (PTMs), as well as extrinsic factors, such as spatial confinement, surface grafting, and NTR occupancy, in determining the functional behaviors of FG-Nups in the NPC. Yet, even with cutting-edge genetic and biochemical tools, it remains difficult to precisely manipulate (e.g., insert, delete, modify, reposition) FG-Nups in cells, hampering our ability to pinpoint the regulatory role of each determinant. Consequently, dissecting the nucleocytoplasmic transport mechanism calls for the analysis of the conformations, dynamics, and interactions of FG-Nups in an advanced *in vitro* model system where the composition and positioning of these intrinsically disordered proteins (IDPs) can be exquisitely controlled in an NPC-like architecture.

In a bid to build programmable NPC mimics, nanopores functionalized with FG-Nups represent a particularly promising route. With lumens coated by a monolayer of FG-Nups, solid-state nanopores enabled NTR-facilitated cargo translocation with tunable permeability in ways resembling selective transport through the NPC^22–26^. Tethering FG-Nups in DNA-origami nanochannels achieved similar results and afforded additional engineering power and experimental flexibility: these water-soluble NPC mimics (termed NuPODs, for Nups organized on DNA) offer exquisite control over the positioning and stoichiometry of individual Nups and compatibility with a broad spectrum of biochemical and high-resolution microscopy analyses such as AFM and transmission electron microscopy (TEM)^27–32^. Importantly, in both biomimetic systems, the permeability of reconstituted FG-Nup barriers can be modulated by the FG-domain composition and density as well as pore geometry.

Despite the remarkable progress, no existing *in vitro* system captures the NPC’s mechanical adaptability, namely the dynamic expansion and constriction at different cellular states and locations as recently illustrated by *in situ* cryo-EM and super-resolution microscopy^33–35^. Though its functional consequences remain unclear, existing data suggest NPC dilation may regulate the NPC’s permeability to macromolecules and the bulk flow of water into and out of the nucleus^34,36^. Compared to FG-Nup composition, the geometry and mechanics of the NPC are arguably even harder to deterministically control due to heterogeneous NPC diameters and the lack of knowledge about the forces driving nuclear pore dilation. Additionally, although FG-Nup interactions have been recognized for their roles in mediating NPC assembly^37,38^, the dilatability of matured NPCs has so far been attributed to the core scaffold and linker Nups that define the NPC architecture^39–41^. The possible contribution of FG-Nup interactions in the transport channel to the NPC’s dilatory state has not been studied. Thus, experimentally probing the behaviors of FG-Nup collectives in the elastic NPC channel necessitates NPC mimics with expandable structural scaffolds.

Here we present expandable NuPODs to address these needs. DNA origami enables the bottom-up construction of nanostructures with well-defined geometry, but also with programmable motion and mechanics^42,43^. Combining the design principles of tensegrity^44^, curved DNA helix bundles^45^, and self-limiting hierarchical oligomerization^46^, we engineered a set of expandable DNA origami rings with the inner diameter (*d*_in_) spanning an identical dynamic range (46–76 nm) but with different elastic energy landscapes. Within these dynamic scaffolds, homotypic interactions of tethered FG-domains from different Nup species generated outward expansion (Nsp1) or inward contractile forces (Nup62, Nup153, Nup98 and Nup214) that change the dilatory state of the ring. Single-particle TEM analysis of the expandable NuPODs revealed a range of DNA ring diameters and distinct FG-Nup morphologies. Leveraging computational modeling and high-speed AFM (HS-AFM) experiments, we provide quantitative measurements of the collective FG cohesion strength and insights into the underlying IDP conformational dynamics. Taking advantage of the programmability of the platform, we show that protein stoichiometry, NTRs, and *O*-GlcNAcylation differentially modulate the FG-Nup interactions. The expandable NuPODs thus offer an enabling platform for studying the biophysical states and functional interactions of FG-Nups, and ultimately the regulation of NPC permeability.

## Results

### Expandable DNA origami rings with programmable mechanics

We have previously built DNA origami channels with fixed widths (40, 45, 60, and 79 nm) to host FG-Nups at designated locations with programmable stoichiometry (up to 96 copies), creating bottom-up NPC mimics^27,30,31^. Motivated by recently discovered NPC structural plasticity, we set out to design mechanically expandable DNA origami channels with the following considerations. First, the channel width needs to cover the observed range of central channel diameters of native NPCs, which, in human cells, span from ∼45 nm in the constricted state to ∼70 nm in the fully dilated state^33,34,47^. Second, to mimic NPC geometry, the DNA channels need to maintain their overall cylindrical shape and depth as they dilate and constrict, thus precluding the use of existing dynamic DNA nanopores/tubes whose size reconfiguration is accompanied by major changes in aspect ratio^48–53^. Third, since the elasticity of the NPC scaffold has not been quantitatively measured and may vary within and across cells considering the NPC’s compositional heterogeneity^54^, the expandable NuPODs under development should have tunable mechanical dilatability. With these criteria in mind, we designed a mechanically responsive DNA channel as a multi-segment, string-suspended cylinder (46–76 nm wide, 15 nm deep) within a stationary octagonal frame (**Fig. 1a, Fig. S1**). In practice, to achieve the target dimensions and sufficient stability, we built the entire structure (hereafter referred to as the expandable DNA ring) as a homo-tetramer, with each quarter ring comprising two 45° inner arcs connected to each other by four 72-base single-stranded DNA ‘strings’ and to a bracket-shaped outer frame by eight 51-base strings (**Fig. 1b**). The termini of both outer and inner domains display complementary DNA sequences to enable self-limiting head-to-tail oligomerization (**Fig. 1b**). By design, the opposing elastic forces exerted by DNA strings reach an equilibrium to define the resting conformation of the expandable DNA ring: the arc-to-frame strings pull the arcs apart, while the strings between inner arcs draw them towards each other. Following one-pot thermal annealing and rate-zonal centrifugation (see Methods), we obtained self-assembled ‘Free’ rings with the expected concentric channel geometry, great size homogeneity (*d*_in_ = 61.2 ± 1.3 nm, measured by negative-stain TEM, **Fig. 1d**), and satisfactory yield (∼35%, estimated by agarose gel electrophoresis, **Fig. S2**). Similarly, changing the inter-arc strings to stiff struts (uninterrupted DNA duplexes), semi-flexible joints (two duplexes flanking a 4-base single-stranded domain), or closed zippers (anti-parallel two-helix bundles) created ‘Open’, ‘Hinged’, and ‘Closed’ rings with *d*_in_ of 73.3 ± 0.9, 69.6 ± 1.4, and 55.3 ± 1.8 nm (**Fig. 1c,e**, See **Table S1** for *d*_in_ of all DNA rings and NuPODs), respectively.

**Fig. 1.**
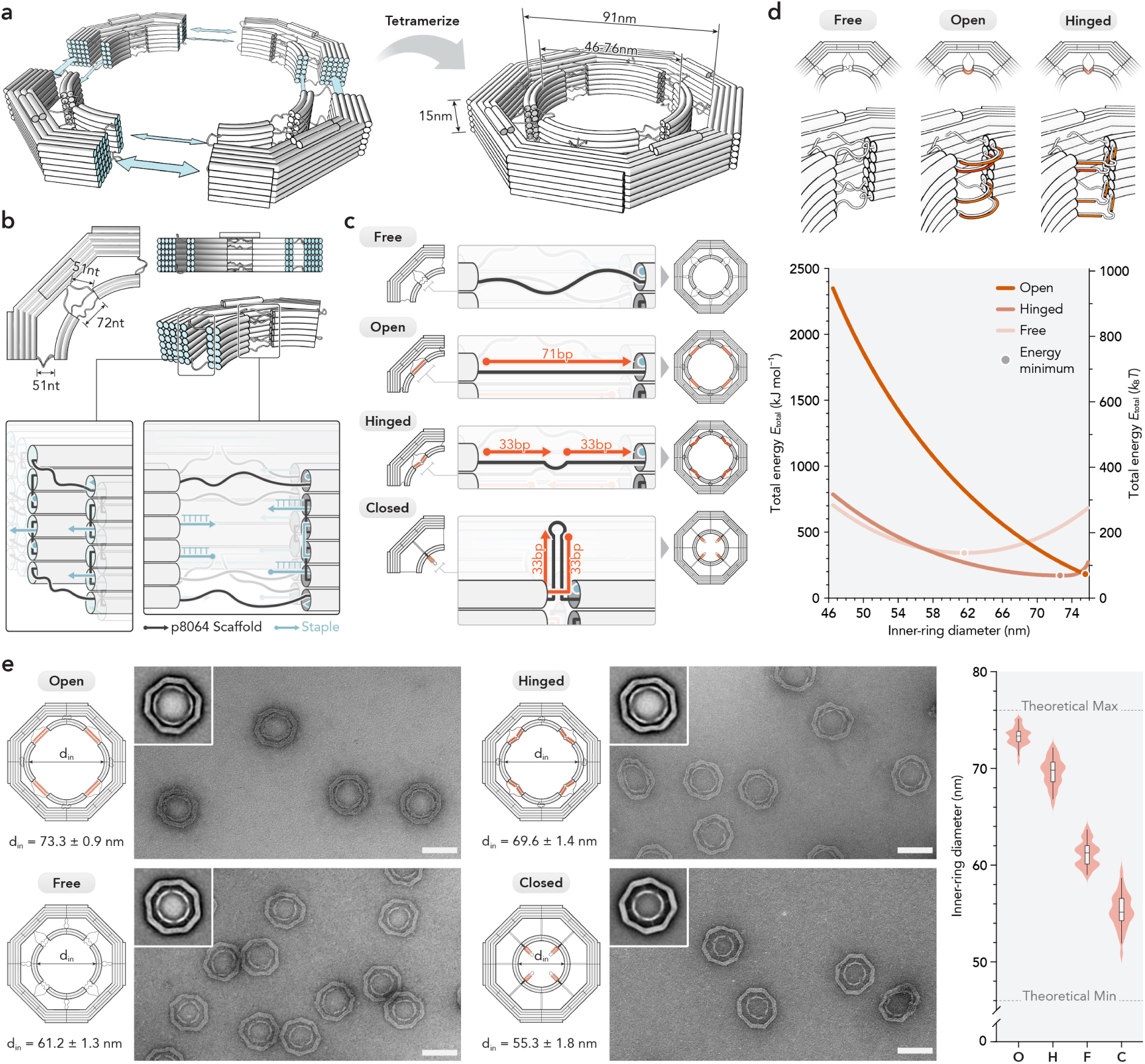
Expandable DNA origami ring with programmable configurations. a, Schematic of tetramerization of four p8064-based monomers into a complete expandable ring. Key dimensions, including the dynamic inner-ring diameters (*d*_in_) are indicated. **b,** Closeup views of the design of a monomeric quarter ring. Four 72-base single-stranded DNA strings connect adjacent inner arcs, and eight 51-base strings connect the inner arcs to the outer frame. Insets show the sticky ends at a terminus of the monomer to enable oligomerization (left) and poly-thymine extensions in between string-connected arcs to deter unwanted base stacking (right). **c,** Expandable rings in four configurations. Orange strands denote string-binding DNA strands that define the ring configuration. **d**, Calculated elastic energy as functions of *d*_in_. The energy minimum predicts the equilibrium *d*_in_ for each configuration. Bending of the DNA connectors in the Open ring model is exaggerated for clarity. **e,** Representative negative-stain TEM images of the four DNA ring configurations. Insets show class averages. *d*_in_ values are listed as mean and s.d.; *n* = 43, 53, 50, and 34 for Open (**O**), Hinged (**H**), Free (**F**), and Closed (**C**) rings, respectively. Lines, boxes, and whiskers in violin plots show the median, 25th–75th, and 5th–95th percentiles, respectively. Scale bars, 100 nm.

In addition to the different diameters at equilibrium, the Free, Open, and Hinged DNA rings should have distinct energetics and hence respond differently to mechanical forces, such as the repulsion and cohesion between ring-tethered IDPs. It is thus useful to establish the expandable DNA rings’ elasticity on theoretical grounds. For simplicity, here we analyzed the mechanical compliance of the inter-arc and arc-to-frame connectors (i.e., DNA strings, struts, and joints) using the well-established worm-like chain models^55^ while considering DNA multi-helix bundles (persistence length ≫ 1 µm) as rigid bodies. We ignored the electrostatic interactions between the frames and arcs, as they are typically more than 5 Debye lengths (∼3.5 nm) apart under the experimental conditions (ionic strength = 0.18 M). Gratifyingly, the theoretical energy minima of the Free, Open, and Hinged DNA rings derived from this analysis corresponded to *d*_in_ of 61.6, 75.4, and 72.8 nm, respectively, closely matching experimentally measured values (**Fig. 1d, Fig. S3**). Furthermore, their energy landscapes provided guidance to subsequent studies of FG-Nup interactions. For example, although the Open and Hinged rings have similar resting *d*_in_, the energy cost to constrict the Hinged ring is substantially less (∼5 *k*_B_*T* nm^-1^ on average) than the Open ring (∼19 *k*_B_*T* nm^-1^), making the former more sensitive for measuring weak to modest protein cohesion. For context, typical NTR–FG-Nup binding energy is ∼13–16 *k*_B_*T* (*K*_D_ = 0.1–2 µM)^56,57^. On the other hand, a Free DNA ring at equilibrium has roughly equal energy barriers to dilate and constrict (∼6 *k*_B_*T* nm^-1^), making it potentially suitable for sensing both repulsion and attraction.

### DNA-guided Nup-assembly forms expandable NuPODs

Having established programmable control over the size and mechanics of the expandable DNA ring, we next functionalized its interior with FG-Nups to reconstitute an FG-lined NPC-like nanopore. A DNA ring displays 48 identical inward-facing DNA handles (12 on a quarter ring, spaced 5–6 nm apart) (**Fig. S4**), which recruits complementary DNA oligonucleotides (anti-handles) conjugated to FG-Nups (**Fig. 2a**). To sample disordered Nup domains with different FG motifs and charge density, we chose full-length human Nup62 (aa 1–522), the entire FG domain of budding yeast Nsp1 (aa 2–603), human Nup153 (aa 896–1475) and human Nup98 (aa 1–498), and the C-terminal FG fragment of human Nup214 (aa 1912–2090) for this study (**Fig. 2b, Table S2**). Following established protocols (see Methods for details), we expressed recombinant Nups as fusion proteins with maltose-binding protein (MBP)-SUMO and SNAP-tag, and chemically conjugated them with benzylguanine-modified anti-handles (**Fig. S5–S7**). The C-terminal Nup214 fragment (179 aa) is much shorter than the other FG-Nups (500–600 aa) tested. To compensate for the difference, we used 64-bp DNA duplexes for tethering Nup214 and 21-bp for all other Nups (**Fig. S8**). Combining 5 Nup species (Nup62, Nsp1^FG^, Nup153^FG^, Nup153^FG^, Nup214^C-term^ ^FG^) and 3 expandable DNA ring configurations (Hinged, Free, and Open), we generated 15 types of expandable NuPODs. After purification by rate-zonal centrifugation, all NuPODs electrophoresed as a sharp band in agarose gel with lower mobility compared to the corresponding Nup-free (apo) DNA rings, suggesting homogeneous populations of well-behaving NPC mimics (**Fig. S10–S30**). Protein quantification by Western blot showed an average of ∼46 copies of Nup62 per purified NuPOD or ∼95% Nup loading efficiency (**Fig. S9**). Imaging expandable NuPODs by negative-stain TEM not only resolves the collective FG-Nup morphology but also measures the DNA ring size change caused by the tethered FG-Nups, both indicative of the nature (attraction or repulsion) and strength of Nup interactions.

**Fig. 2.**
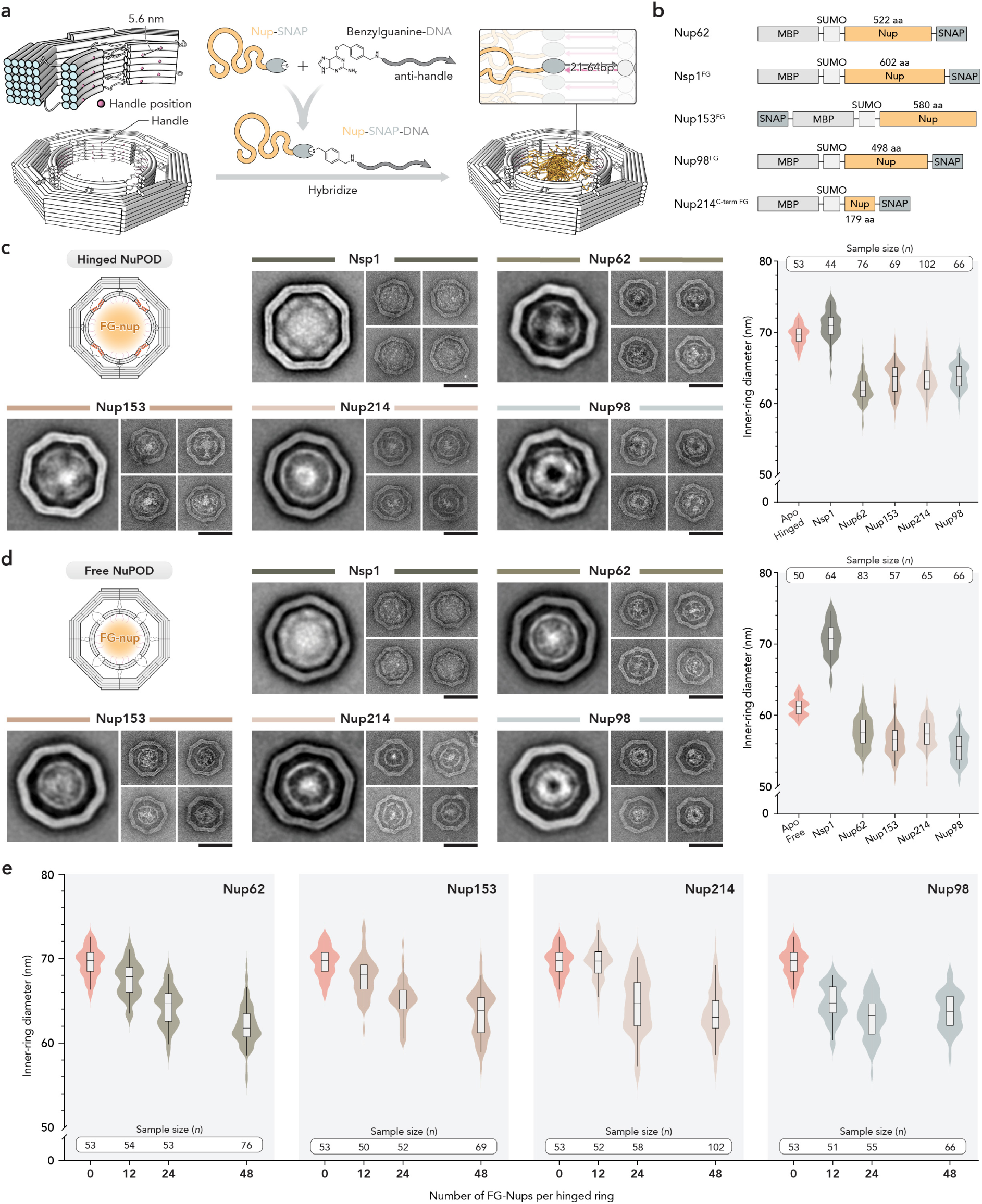
Expandable NuPODs with distinct ring sizes and FG-Nup morphologies. a, Schematics of Nup-DNA conjugation and DNA-hybridization–guided tethering of FG-Nups to the 48 handles in an expandable DNA ring. DNA tethers are 64-bp long for Nup214 and 21-bp for other Nups unless specified. **b,** Domains of the recombinant FG-Nup constructs. **c**, Schematic of Hinged NuPOD (top left), negative-stain TEM analyses including a class average and four representative images for each NuPOD type (middle), and measured *d*_in_ (right). Scale bars, 100 nm. **d**, same as **c**, but for Free-NuPODs. **e**, *d*_in_ of Hinged rings containing 0, 12, 24, or 48 copies of FG-Nups by design. Lines, boxes, and whiskers in violin plots show the median, 25th–75th, and 5th–95th percentiles, respectively.

### Nup62 and Nsp1 have opposite effects on NuPOD width

We first studied human Nup62 and its yeast homolog Nsp1, both featuring an N-terminal disordered domain rich in FxFG motifs^58^ and chiefly (32 out of 48 copies) residing in the NPC’s central channel^3,4^. Despite the similarities, computational analyses predict the FG domain of Nup62 to be cohesive (i.e., strong self-interacting propensity) and Nsp1 non-cohesive or even partially repulsive, mainly due to vastly more charged residues in the latter^59^. Experimental characterizations of their cohesiveness have produced inconsistent results, and a head-to-head comparison between the two is lacking. For example, although bead-immobilized Nsp1^FG^ does not bind other FG-Nups or itself in solution^6^, soluble Nsp1^FG^ has been shown to form hydrogels^7,8,60^. Likewise, overexpressed FG-domain of Nup62 was found highly mobile throughout live cells with no detectable self-association in the nucleus or cytoplasm^61^, while concentrated Nup62 jellifies rapidly *in vitro*^10^. Since the mechanically responsive DNA rings can host FG-Nups with physiologically relevant stoichiometry and spatial organization, we loaded up to 48 copies of Nup62 or Nsp1^FG^ to form expandable NuPODs and assessed the FG-Nup interactions by negative-stain TEM. Loading Nup62 into Hinged DNA rings (*d*_in_ = 69.6 ± 1.4 nm) produced ∼8-nm ring contraction (*d*_in_ = 62.0 ± 2.0 nm) and dense puncta near the center of NuPODs, suggesting cross-channel cohesive interactions among ring-grafted proteins (**Fig. 2c, Fig. S10**). In contrast, tethering Nsp1^FG^ via the same DNA handles led to a small expansion of Hinged rings (*d*_in_ = 71.0 ± 2.1 nm), implying that Nsp1^FG^ may repel one another within the confined space instead of cohering. Consistent with this, Nsp1^FG^ had a diffuse appearance in NuPODs with no obvious dense spot (**Fig. 2c, Fig. S11**), resembling extended non-cohesive brushes. Cleaving the MBP-SUMO tag by TEV protease had negligible effect on the Nup62 and Nsp1^FG^ NuPOD diameters (*d*_in_ = 61.9 ± 1.6 nm and 69.4 ± 1.7 nm, respectively, **Fig. S12**) or protein morphologies, demonstrating that the FG-Nups, instead of the fusion tags, determined the protein behaviors. Mutating 18 Phe in the FG domain of Nup62 to Ser (Nup62^F→S^) disrupted the dense protein cluster and diminished NuPOD contraction (*d*_in_ = 68.3 ± 1.9 nm, **Fig. S13**), underscoring the contribution of FG repeats to the Nup62 cohesivity. Further supporting this, surface plasmon resonance (SPR) experiments measured strong binding avidity (*K*_D_ = 0.6 ± 0.15 µM) of soluble wildtype Nup62 to a surface-tethered layer of Nup62 but detected no binding between soluble and immobilized Nup62^F→S^ at similar grafting density (**Fig. S33**). The contrasting behaviors of Nup62 and Nsp1^FG^ persisted in the Free and Open NuPODs: in the smaller Free NuPODs (*d*_in_ = 61.2 ± 1.3 nm before loading Nups), Nup62 shrank the ring modestly (*d*_in_ = 57.7 ± 2.3 nm), whereas Nsp1^FG^ generated pronounced expansion by ∼10 nm (*d*_in_ = 70.7 ± 2.4 nm, **Fig. 2d**); in the largest and stiffest Open NuPODs, Nup62 constricted DNA rings by ∼9 nm while Nsp1^FG^ had minimal effects on the ring diameter (**Fig. S14**). In all NuPOD configurations tested, Nup62 and Nsp1^FG^ could both extend far into the DNA ring to occlude the channel, despite their different morphology and effects on ring size. Together, these results provided strong evidence for the distinct self-interacting tendencies of Nup62 and Nsp1 in an NPC-like geometrical confinement and validated the utility of expandable NuPODs for studying the collective behaviors of nanopore-tethered IDPs.

### Several human FG-Nups contract NuPODs but with different protein morphologies and stoichiometry dependency

In addition to Nup62, we tested three human FG-Nups: Nup153, Nup214 and Nup98, which are located at the nuclear basket, cytoplasmic periphery, and central channel of the NPC, respectively. In Hinged, Free and Open DNA rings with 48 handles, tethering Nup153^FG^, Nup214^C-term^ ^FG^, or Nup98^FG^ markedly contracted DNA rings to a similar extent as the Nup62-NuPODs (**Fig. 2c,d and Fig. S15–S17**), reflecting considerable cohesiveness of all four human FG-Nups. The rank orders of NuPOD diameter vary by DNA ring configurations, with *d*_in_ of the widest and narrowest NuPODs in each configuration group differing by no more than 2 nm. Nup62, Nup153 and Nup214 NuPODs displayed a cross-channel meshwork with a clear protein-dense region at the center. Among the three, Nup214^C-term^ ^FG^ formed the smallest dense spot, as expected from its small mass; shorter DNA tethers resulted in separate, local protein clusters (**Fig. S18,S19**). Strikingly, the Nup98-NuPODs showed a unique donut-shaped protein mass close to the DNA ring (**Fig. 2c,d and Fig. S17**) reminiscent of collapsed polymer chains^13^, meaning the strong cohesion of Nup98^FG^, either intramolecularly or among neighboring Nups, likely prevented them from bridging across the DNA channel. These behaviors mirrored our prior SPR measurements showing that at similar surface grafting densities, Nup98’s GLFG-rich domain is more compact than the disordered domains of Nup62, Nup214 and Nup153, in which other FG-motifs (e.g., FxFG, SxFG, PxFG, FG) dominate^62^. The fact that Nup62 and Nup98^FG^, with similar length (498 vs 522 aa) and stoichiometry, exhibited drastically different IDP compactness regardless of DNA channel widths (∼55–73 nm without Nups) strongly suggests their inherently different cohesiveness. To ascertain the difference, we reduced the number of Nup-tethering handles in the Hinged DNA ring from 48 to 24 and 12 (**Fig. 2e, Fig. S20–S24**). Indeed, while all four human NuPODs generally became less constricted at decreased FG-Nup stoichiometry, *d*_in_ of the Nup98-NuPOD was consistently the smallest. As an illustrative comparison, it took ∼48 Nup62 to shrink NuPODs to roughly the same size as those containing ∼24 Nup98^FG^. Furthermore, Nup98^FG^ was the only FG-Nup showing a collapsed conformation at all grafting densities (**Fig. S24**). We therefore conclude that Nup98 contains the most cohesive FG-domain among all Nups tested, consistent with the observation that Nup98 forms the most restrictive hydrogels among major vertebrate FG-Nups^10^.

### Coarse-grained MD simulations corroborate NuPOD size and morphology measured by TEM

We simulated individual NuPOD particles using coarse-grained MD models to assess their structure and energetics. An mrDNA polymer model^63^ (4 bp per bead resolution) was constructed for each of the Open, Free, Hinged, and Closed DNA rings, starting with an idealized initial configuration mapped from the DNA origami design file. As the initial configuration included overbent and overstretched sections of DNA, the rings quickly contracted as the system relaxed (**Fig. 3a**), resulting in steady-state conformations with *d*_in_ in overall good agreement with the experimental results (**Fig. 3b, 1e**). The *d*_in_ distributions, taken from the final 10 µs of the simulation, are noticeably narrower compared to experimental values, especially for the Closed ring, which could be attributed to local defects in the DNA origami assembly and the idealized base pairing (e.g., lack of fraying) in the mrDNA model.

**Fig. 3.**
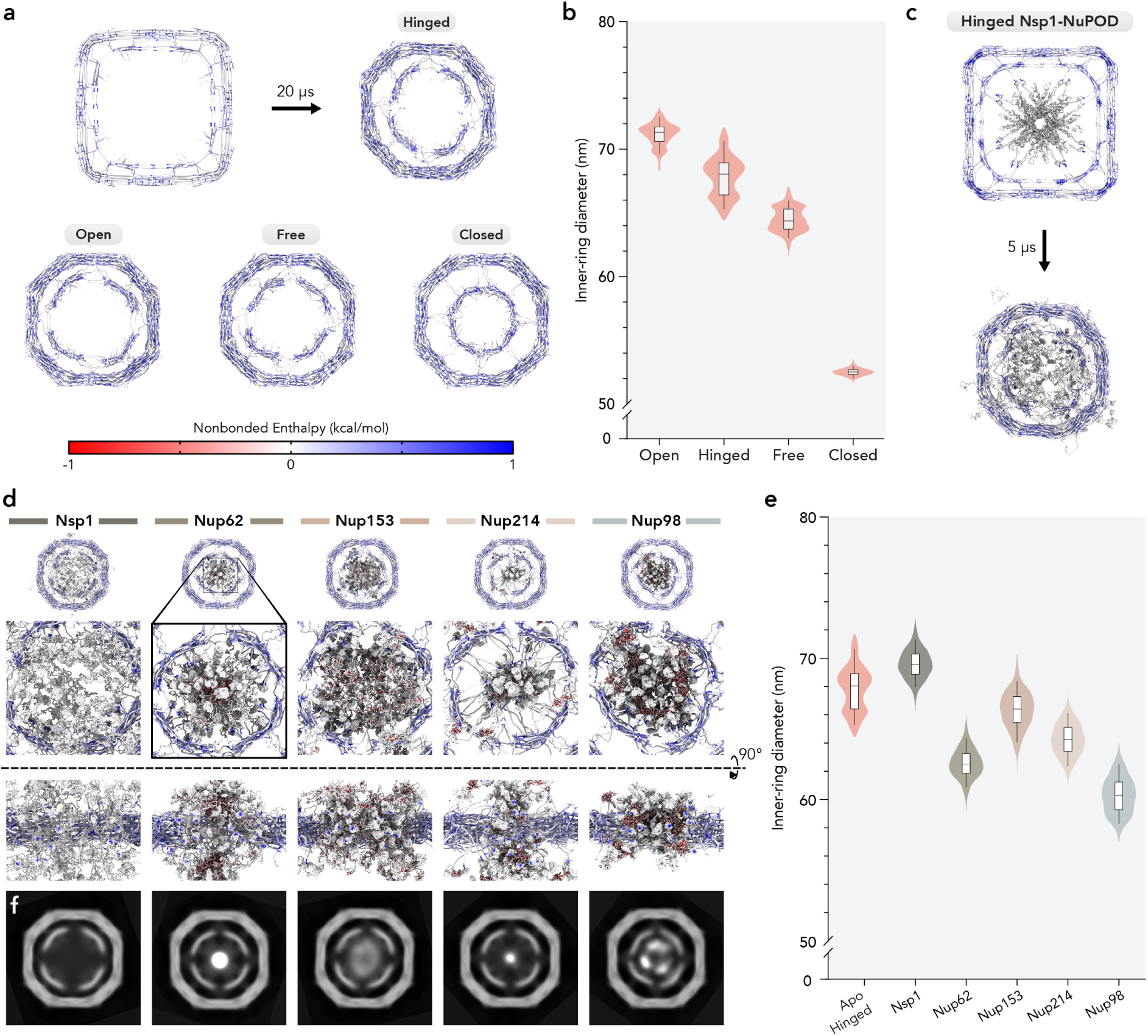
Structure of FG-Nups in molecular dynamics simulations. **a**, Relaxation of Hinged (top), Open (bottom left), Free (bottom center) and Closed (bottom right) Nup-free DNA rings. **b**, Inner ring diameters of NuPODs shown in **a**, sampled after first 10 μs of simulation. **c**, Relaxation of a Hinged NuPOD containing 48 copies of Nsp1^FG^. **d**, Snapshots showing the distribution of five different FG-Nups in Hinged NuPODs. **e**, Inner ring diameters of NuPODs during the second half of simulation. **f**, The simulated mass density in each NuPOD including DNA and disordered peptides averaged over the simulation trajectory. Lines, boxes, and whiskers in violin plots show the median, 25th–75th, and 5th–95th percentiles, respectively.

The mrDNA model was then combined with a model developed by the Onck lab (single amino acid resolution) specifically for disordered FG-Nup polypeptides^12,22^ to simulate Hinged NuPODs containing 48 copies of Nsp1^FG^, Nup62, Nup153^FG^, Nup214^C-term^ ^FG^ and Nup98^FG^. Notably, the NuPOD models consider both physical link and non-bonded interactions (e.g., screened Coulomb electrostatics and Lennard-Jones potential) between DNA and proteins (See Methods for details). Upon relaxation, the Nups quickly formed mesh-like networks predominantly within an axially extended cylindrical space defined by the inner ring, with a few FG-Nups straying outside this space occasionally (**Fig. 3c**). Close inspection of the meshwork shows diverse structures of varying compactness, with Nsp1 having the most diffuse conformation (i.e., most extended and porous network), and Nup98 the densest (**Fig. 3d**, **movies S1–S5**), consistent with TEM observations. Overall, *d*_in_ of simulated NuPODs aligned well with the experimental data (**Fig. 3e**, **2c**). As seen in the TEM analyses, Nsp1 caused slight expansion of the DNA ring, and all other Nup species caused various degrees of contraction. Finally, to visualize the FG-domain morphologies and better compare with experimental observations, we performed ‘particle averaging’ on NuPODs from the top view over the entire simulation trajectories, excluding the folded protein domains. These computationally generated images qualitatively resembled the negative-stain TEM 2D averages (**Fig. 3f**, **2c**) and revealed an annular density for Nup98^FG^ that was not apparent in snapshots taken from the trajectory. Therefore, in addition to validating the metrics observed in experiments, the simulations provide a high-resolution view of the mesh structures, reinforcing that unstructured FG-domains are the main drivers of protein behaviors in NuPODs.

### The mechanical work to resize a NuPOD correlates with the free energy of Nup–Nup interactions

The elastic deformation of the NuPOD by an FG-Nup network can provide quantitative information about the Nup–Nup interactions. Using experimentally measured *d*_in_ of NuPODs containing human FG-Nups (**Fig. 2c,d**) and theoretical prediction of the expandable rings’ energy landscape (**Fig. 1d**), we estimated ∼10–20 and 20–35 *k*_B_*T* elastic potential energy in a Free and a Hinged NuPOD caused by FG-Nup association, respectively. For Open NuPODs, the estimated elastic energy was much higher (∼100–160 *k*_B_*T*). However, these probably represent only a fraction of the total cohesive energy of FG-Nups in a NuPOD, as intramolecular or local associations, such as FG-Nups tethered to the same quarter ring, are unable to contract the DNA ring. In other words, cohesive FG-Nups may only bear mechanical load when they bind other Nups across the channel. Moreover, steric hindrance between Nups helps NuPODs resist contraction, which may explain why smaller rings generally constricted less upon loading FG-Nups (Free, Hinged, and Open rings contracted by 3.5–5.5, 5.8–7.6 and 6.1–8.9 nm, respectively). Thus, substantial amount of free energy generated by FG-Nup cohesion, especially in smaller DNA rings, was most likely dissipated by the flexible IDP chains instead of converted to mechanical work to resize DNA rings.

To gain further quantitative insights into the DNA ring deformation and FG-Nup cohesion, we employed MD simulations, which provide direct access to energetics and can estimate forces and associated free energy changes in a dynamic system. First, we evaluated the free energy required to resize NuPODs using thermodynamic integration. Specifically, for a given Nup species, we selected a relaxed state of the NuPOD from our prior equilibrium simulation, removed the DNA, and placed a harmonic restraint on the terminal amino acid tethering each Nup to the DNA scaffold (**Fig. 4a**). The harmonic restraint preserves the global geometry imposed by the NuPOD but also reports on the force acting on each Nup. Then, in a series of simulations, we moved the restraint points outward in 2 Å steps until they would reach a size equivalent to the Hinged Nup-free ring. In essence, our simulation recapitulates the reversed pathway of NuPOD contraction, obtaining an average radial force of ∼0.5–1 pN required to hold a Nup at each step (**Fig. 4b**). This force increases slightly as the Nups are pulled apart for all Nup species, suggesting that the system underwent elastic deformation. By integrating the average force along the entire expansion path, we estimated the energy responsible for resizing the NuPOD (**Fig. 4c**) to be 10–60 *k*_B_*T*, which is comparable to the elastic potential energy derived from TEM analyses. The discrepancy can likely be attributed to the difference between simulated and experimentally measured ring sizes.

**Fig. 4.**
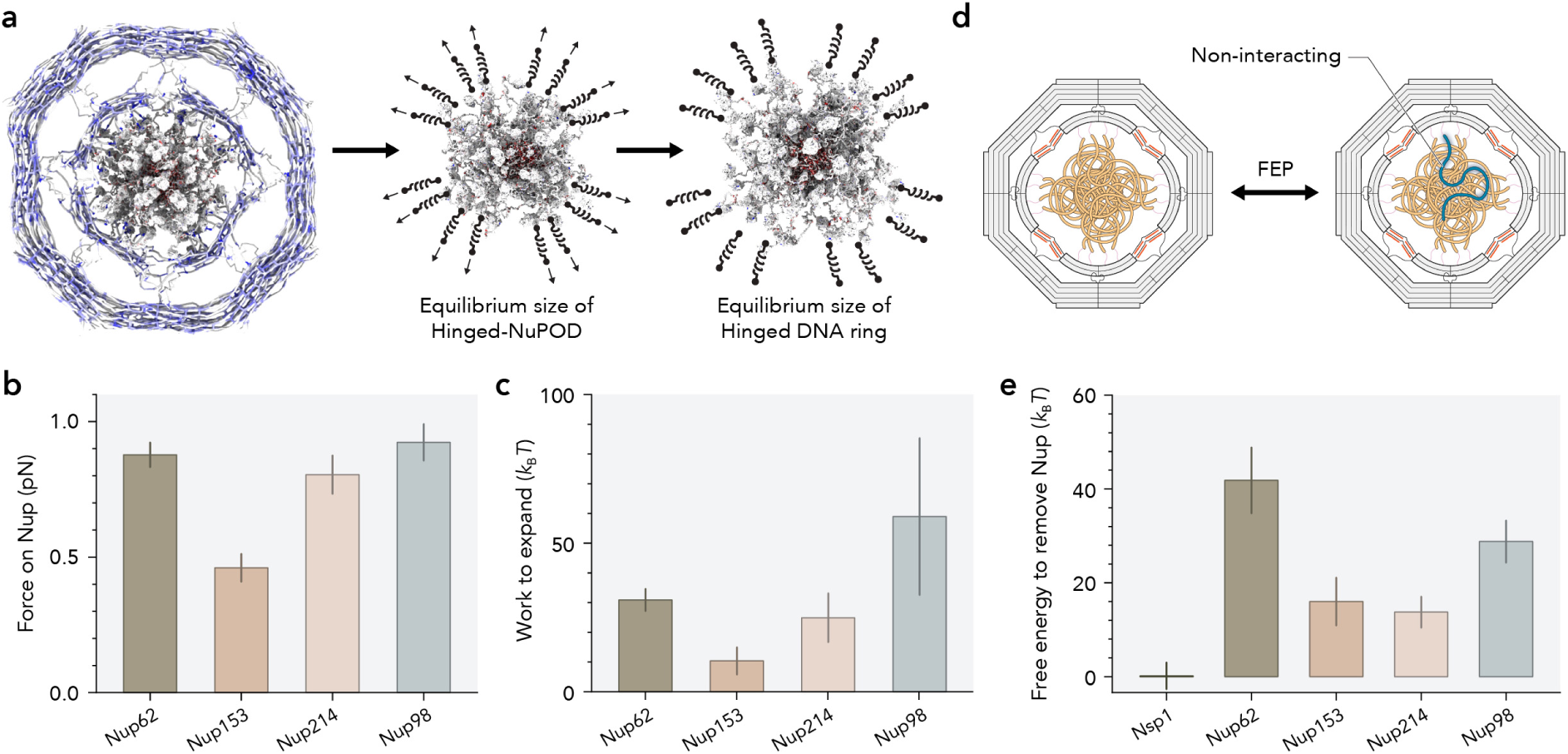
Energetics of NuPOD probed by molecular dynamics simulations. **a**, Schematic illustrating the procedure to estimate the force acting on each Nup. **b**, Average force required to hold FG-Nup mesh observed during a series of simulations in which the restraint points are spread from the steady state radius to a radius corresponding to a Hinged Nup-free DNA ring. **c,** Mechanical work required to spread FG-Nups during simulations described in **a**. In **b** and **c**, error bars report the standard error of the mean (s.e.m.) among the 48 Nup chains. **d**, Schematic depicting a free energy transformation (FEP) whereby a single FG-Nup chain is made to be non-interacting with its neighbors. **e,** Estimated free energy required to fully remove one FG-Nup molecule from forward and reverse FEP simulations. Error bars depict the statistical error reported by the MBAR algorithm.

MD simulations can also be used to more directly inspect the free energy of interactions, provided it is feasible to thoroughly sample configurations to properly assess entropic contributions. To assess the Nup–Nup interaction energy, we employed free energy perturbation (FEP) where, informally, one gradually changes the energy function governing the system and evaluates the free energy cost of doing so. To probe the free energy of a single Nup chain interacting with other Nups in a NuPOD, we gradually removed these interactions from the system before restoring them over the course of 225 ns in each of four replicas per Nup species. The multi-Bennet acceptance ratio (MBAR) algorithm^64^ was used to extract the free energy cost of removing a chain and associated statistical errors (**Fig. 4d**). The costliest Nup chains to remove were Nup62 (∼40 *k*_B_*T*) and Nup98 (∼30 *k*_B_*T*) followed by Nup153 and Nup214 (∼15 *k*_B_*T*), whereas removing Nsp1 incurred no free energy penalty (**Fig. 4e**). The free energy cost of removing a copy of each Nup species generally correlates with (but is not always proportional to) the free energy cost to expand the NuPOD, which is reasonable considering the statistical errors associated with both calculations.

### Kap95 tightens nanopore-tethered Nsp1 into a plug-like structure, shrinking NuPODs

Having established the FG-Nup interactions determined by their intrinsic cohesiveness, grafting density and spatial confinement, we turned our attention to additional modulators, specifically NTRs and PTMs (**Fig. 5a,b**). Native NPCs are laden with NTRs (also known as karyopherins or Kaps) and, in animal cells, PTMs (e.g., *O*-linked β-N-acetylglucosamine or *O*-GlcNAc) on FG-Nups. Previous studies have found (i) NTRs such as Kap95 and importin β1 (Impβ1, human homolog of Kap95) modulate the physical state, dynamics, and permeability barrier of FG-Nups^19,62,65–67^ and (ii) *O*-GlcNAc modification accelerates both passive diffusion and facilitated translocation across the NPC and phase-separated FG-Nups^10,68^. As yeast NPCs lack *O*-GlcNAc, we first tested the effect of Kap95 on expandable Nsp1-NuPODs for simplicity. Kap95 is the principal yeast importin and binds multiple FG-Nups including Nsp1^57^; like Impβ1, it contains multiple FG-binding pockets that can simultaneously engage FG repeats^69,70^. Addition of 1 µM Kap95 to Free Nsp1-NuPODs significantly contracted the DNA ring, reducing *d*_in_ from 70.7 ± 2.4 to 62.4 ± 4.0 nm, and drastically changed the collective protein morphology – from the diffuse, thin haze in the absence of NTRs to a centrally localized dense cluster visible by negative-stain TEM (**Fig. 5d, S25**). Similar changes were observed in Hinged Nsp1-NuPODs upon Kap95 treatment, albeit with smaller ring size reduction (**Fig. S25**). The Kap95-dependent plug-like protein structure is consistent with our previous AFM study on purified NPCs and nonelastic Nsp1-NuPODs^19^. Notably, adding Kap95 broadened the size distribution of Free Nsp1-NuPODs, suggesting that while Kap95 tightens the Nsp1 mesh, the resulting protein network remains internally heterogenous and dynamic. These results demonstrate that expandable NuPODs can detect NTR-induced changes in FG-Nup interactions at the single-pore level and provide direct evidence for Kap95-driven coalescence of an otherwise non-cohesive FG-Nup in a confined space.

**Fig. 5.**
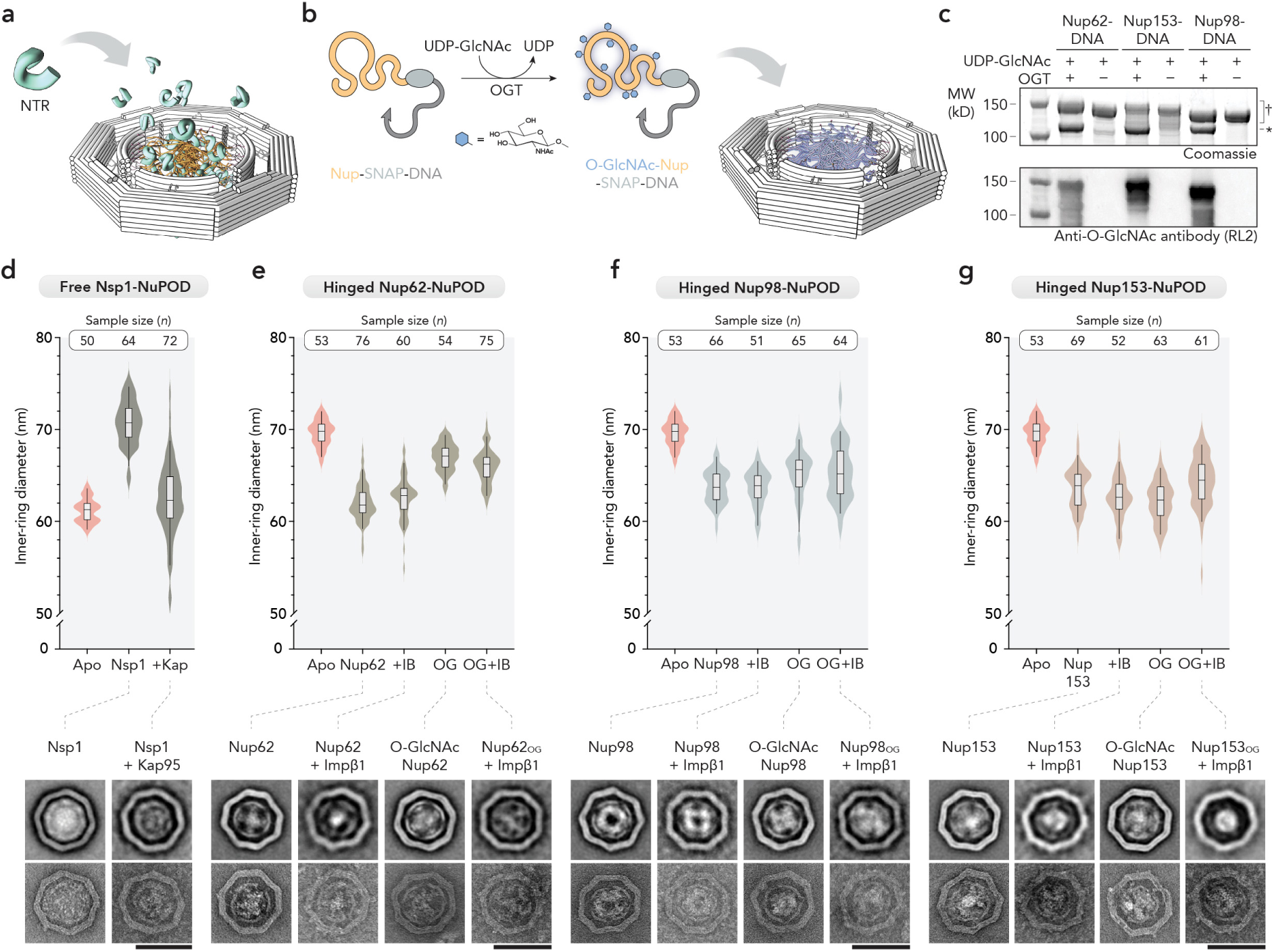
NTRs and *O*-GlcNAcylation modulate the NuPOD size and protein morphology. a, Schematic of NTR binding to FG-Nups within a NuPOD. **b,** Schematic of producing *O*-GlcNAc modified NuPODs. FG-Nups are enzymatically modified using *O*-GlcNAc transferase (OGT) and UDP-GlcNAc before loaded into the expandable ring. **c,** Validation of Nup *O*-GlcNAcylation by SDS-PAGE stained by Coomassie Brilliant Blue (top) and Western blot using an antibody against *O*-GlcNAc (bottom). Gels are loaded with FG-Nups treated with (+) or without (−) OGT. † denotes Nup-DNA conjugates with varying MW; * denotes OGT. **d,** Negative-stain TEM images and measured *d*_in_ of Free Nsp1-NuPODs in the absence or presence of Kap95. **e**–**g,** Negative-stain TEM images and measured *d*_in_ of Hinged-NuPODs containing Nup62 (**e**), Nup98 (**f**), or Nup153 (**g**) with or without Impβ1 binding (IB) and/or *O*-GlcNAcylation (OG). For **d**–**g**, Nup-free (Apo) ring sizes are taken from Fig. 1 and shown here for comparison. Class average TEM images are shown above representative single-particle images. Scale bars, 100 nm. Lines, boxes, and whiskers in violin plots show the median, 25th–75th, and 5th–95th percentiles, respectively.

### *O*-GlcNAcylation and Impβ1 remodel NuPODs containing cohesive FG-Nups

We next asked how IDP networks formed by cohesive FG-Nups are modulated by *O*-GlcNAcylation and Impβ1. Recent work measuring the dynamics of fluorescently labeled Nup98 in live cell NPCs and in reconstituted condensates found *O*-GlcNAc and Impβ1 work cooperatively to keep FG domains in a dynamic, liquid-like state in the NPC^21^. However, effects of *O*-GlcNAc and Impβ1, in isolation or combined, on nanopore-tethered FG-Nup species other than Nup98 have not been characterized experimentally. To bridge this gap, we examined three human FG-Nups (Nup62, Nup98, and Nup153) grafted on Hinged NuPODs in their unmodified and glycosylated forms before and after Impβ1 treatment. For unmodified Nup62, adding 1 µM Impβ1 did not significantly alter NuPOD size in 1 hour, despite increased contrast of the protein cluster indicative of Nup–NTR binding. Meanwhile, tethering *O*-GlcNAcylated Nup62 (Nup62_OG_, generated enzymatically, **Fig. 5b,c**) formed NuPODs with substantially increased diameter (*d*_in_ = 67.0 ± 1.5 nm, compared to 62.0 ± 2.0 nm for unmodified Nup62) (**Fig. 5e, Fig. S26,S27**), showing that *O*-GlcNAcylation reduced Nup62 cohesiveness and supporting a model in which hydrophilic *O*-GlcNAc moieties sterically hinder hydrophobic FG-FG contacts to enhance permeability of the FG-Nup barrier^68^. Additional Impβ1 had minimal effect on the size of Nup62_OG_-NuPODs. Nup98 displayed a distinctive morphological response to *O*-GlcNAcylation: while unmodified Nup98 primarily accumulated as a donut-like density along the inner periphery of the DNA ring, leaving a protein-scarce zone at the center, Nup98_OG_ became almost evenly distributed, filling a slightly dilated DNA channel (*d*_in_ = 65.1 ± 2.6 nm vs 63.8 ± 1.8 nm for unmodified Nup98-NuPOD) (**Fig. 5f, Fig. S28,S29**). We take this morphological change to mean that less cohesive Nup98_OG_ chains switched from a collapsed state to more extended conformations, further supporting the idea that *O*-GlcNAc impedes FG-Nup cohesion. Moreover, unlike unmodified Nup98 NuPODs, which showed no detectable morphological or size change after Impβ1 treatment as expected from the low Impβ1-binding capacity of densely grafted Nup98,^62^ the lumen of Nup98_OG_-NuPODs seemed to be readily perfused by Impβ1 (**Fig. S29**), consistent with enhanced permeability of reconstituted Nup98_OG_ hydrogels to NTRs^10^. Such a conformational transition may reflect the physiological role of *O*-GlcNAcylation in maintaining Nup98 in a more extended and functional state within the NPC. For Nup153 NuPODs, the effects of Impβ1 and *O*-GlcNAcylation on DNA ring width were less pronounced, though Impβ1 binding to both glycosylated and non-modified Nup153 led to more substantial channel-occluding protein mass in TEM class averages (**Fig. 5g, Fig. S30**). Kap95/Impβ1 therefore appeared to interact with FG-Nup collectives in a context-dependent manner: the NTRs have limited capability to enter the most compact FG mesh (unmodified Nup98), readily infiltrate FG-Nup networks consisting of intermediate Nup–Nup cohesions without generating major contractile or expansion force (e.g., Nup98_OG_, Nup62/Nup62_OG_, Nup153/Nup153_OG_), and act as a crosslinker to tighten poorly associated FG-Nup brushes (e.g., Nsp1).

### Impβ1 slows the dynamics of Nup62 in NuPODs

To directly visualize how Impβ1 remodels the DNA ring-tethered Nup62 in real time, we imaged Hinged Nup62-NuPODs by HS-AFM with varying concentrations (0, 100 or 1000 nM) of Impβ1, with the unloaded Hinged ring as a reference (**Fig. 6a–d**). The empty Hinged ring displayed a deep central cavity (*d*_in_ = 69.9 ± 5.9 nm), while Nup62 loading contracted the channel (*d*_in_ = 61.9 ± 6.0 nm) and partially filled the cavity with a diffuse, rapidly fluctuating protein density. Addition of Impβ1 to surface-adsorbed NuPODs revealed a dynamic plug-like cluster within the channel that grew progressively denser with the concentration, until at 1000 nM the central protein mass overfilled the channel and protruded above the DNA ring (**Fig. 6e**), confirming functional Nup62-Impβ1 binding^31^. Diameter of the expandable NuPOD did not significantly change upon Impβ1 treatment (*d*_in_ = 63.1 ± 5.7 nm and 63.3 ± 6.5 nm at 100 and 1000 nM Impβ1, respectively), consistent with our TEM measurements (**Fig. 5e**) and indicating that Impβ1 densifies the FG meshwork without exerting a detectable net radial force on the DNA ring.

**Fig. 6.**
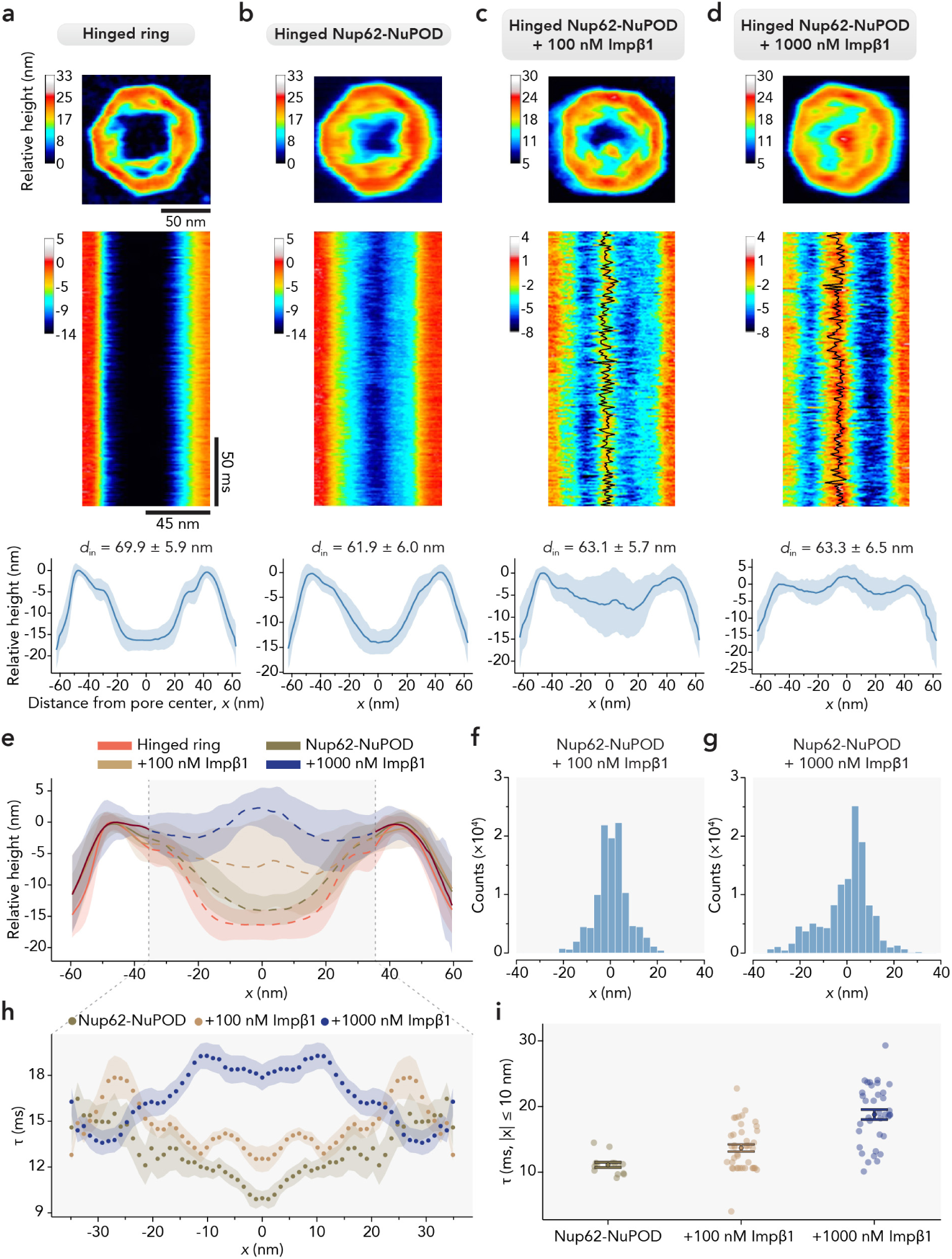
Protein dynamics in Nup62-NuPODs modulated by Impβ1. **a**–**d**, HS-AFM analysis of the Hinged ring (**a**; *n* = 13) and Hinged Nup62-NuPOD with 0 (**b**; *n* = 10), 100 nM (**c**; *n* = 8), and 1000 nM (**d**; *n* = 6) Impβ1. Top, representative HS-AFM images. Middle, representative HS-AFM kymographs of a horizontal line-scan (acquired at 1.875 ms/line) through the ring center. In **c** and **d**, the frame-by-frame trajectory of the central plug-like cluster is overlaid in black. Scale bars in **a** apply to **a**–**d**. Bottom, HS-AFM height profiles. Solid lines and shades indicate the grand mean across all pores and scan directions and s.d., respectively. **e**, Overlaid mean height profiles shown in **a**–**d**. **f**–**g**, Histograms of the protein cluster positions along the scan line for the 100 nM (**f**) and 1000 nM (**g**) Impβ1 conditions, expressed as the distance from the ring center (*x*); bin width, 2 nm; pooled counts from all kymographs. **h**, Autocorrelation decay time *τ* as a function of *x*, derived from per-line autocorrelation analysis of the kymographs of the Hinged Nup62-NuPOD with 0 (*n* = 9), 100 nM (*n* = 8), and 1000 nM (*n* = 7) Impβ1. Shaded regions represent ± 1 s.e.m. **i**, *τ* in the central area (|*x*| ≤ 10 nm) of NuPODs. Color points represent the mean *τ* of individual NuPODs; black dots and bars indicate the grand mean ± s.e.m.

Frame-by-frame tracking in the line-scanning kymographs (1.875 ms/line) showed that the central cluster formed by Nup62 and Impβ1 was highly mobile, fluctuating around a preferential position at the pore axis, with position histograms peaking sharply around the midline in 100 and 1000 nM Impβ1 conditions (**Fig. 6f,g**). Per-line autocorrelation analysis (**Fig. 6h,i**) yielded a characteristic decay time τ that was uniformly low for Nup62 alone (∼11 ms across the channel) – reflecting the rapid fluctuations of FG-domains in the absence of Impβ1 – but rose into a pronounced central peak at 1000 nM Impβ1 (∼19 ms at the midline), corresponding to the emergence of a ‘central plug’ with nearly two-fold slower fluctuations. Together, these HS-AFM data show that Impβ1 binds the ring-tethered Nup62 to slow its local dynamics and stabilize a centrally localized plug in a dose-dependent manner without altering the dilatory state of the ring.

## Discussion

The interactions of FG-Nups with one another and with NTRs underpin the barrier and transport functions of the NPC. While early work using reconstituted FG-Nup gels and droplets provided the general framework to understand the NPC’s selective barrier properties, recent studies have shown that FG-Nups move and interact differently in the NPC compared to in bulk solution or condensates. These findings point to the importance of investigating FG-Nup behaviors in their native environments. However, the immense complexity of the NPC and functional redundancy of its components present seemingly formidable challenges to finetune the NPC’s biochemical and geometrical attributes in cells. Motivated by the need for a model system to capture key features of the NPC, namely the densely grafted FG-Nups within a dilatable scaffold, here we developed DNA-origami–based nuclear pore mimics with dynamically variable widths and tunable mechanical responsiveness. Such expandable NuPODs enabled us to probe FG-Nup interactions in an elastic nanopore by single-particle TEM.

Using primarily two readouts, the NuPOD diameter and IDP morphology, we systematically analyzed the homotypic interactions of five FG-Nup species with various FG motifs, charge densities and intra-NPC localizations, as well as factors that modulate their self-binding propensity, providing the following insights. First, confined in a cylindrical space, Nsp1^FG^ chains repel each other via electrostatic and/or excluded volume interactions, whilst FG-domains of Nup98, Nup62, Nup214, and Nup153 coalesce, with GLFG-rich Nup98 showing the strongest cohesiveness. Second, nanopore-tethered FG-Nups exhibit interacting patterns absent from bulk condensates. For example, unmodified Nup98 chains collapse into an annular area near their anchoring points in NuPODs, supporting the experimental and computational evidence of peripheral enrichment of highly cohesive FG-Nups in the NPC^13,19,21,67^ while contrasting with the bulk condensates, gels and amyloid-like fibrils formed without geometrical constraints. Likewise, Nsp1 and Nup62 showed starkly contrasting cohesivity in NuPODs despite both forming macroscopic gels in solution. Third, cohesive FG-Nup interactions generate substantial contractile force that amounts to 25–40 pN (or 20–35 *k*_B_*T* elastic energy) from the ∼48 Nups in a Hinged NuPOD, as estimated by both experiment and simulation. An attempt to convert Free DNA rings housing cohesive FG-Nups to the Open conformation by rigidifying the inter-arc single-stranded DNA (ssDNA) strings yielded ∼61-nm wide NuPODs, which are ∼10 nm narrower than the protein-free Open ring (**Fig. S31,S32**), confirming that the cohesive FG-Nups can resist expansion force. These findings thus suggest the disordered FG domains may be force bearing and act as flexible adhesives to mechanically modulate NPC dilation. These hypotheses remain to be tested in live cells using force-sensitive fluorescence probes. Fourth, the NTRs’ modulatory effects on FG-Nups depend on relative strengths of Nup–NTR and Nup–Nup associations. This is further regulated by *O*-GlcNAc modification, which weakens FG-Nup cohesiveness and promotes the formation of pore-occluding Nup–NTR meshes characteristic of functional NPCs. Thus, in the NPC where multiple FG-Nup species with distinct cohesiveness and extensibility are anchored to different regions, spatial distributions of FG-domains and NTRs are likely uneven, which may result in different molecular trafficking ‘pathways’ or ‘active zones’. Recently observed concentric zones of Impβ1-mediated cargo transport by MINFLUX microscopy^71^ support this possibility, though the molecular identities of FG-domains in each zone remain to be determined.

Despite spanning a broad range of cohesivities, the FG-domains of Nup62, Nup153 and Nup214 extended well beyond their hydrodynamic sizes^59,62^ to form cohesive interactions at the pore center. Rather than mandating a static mesh, moderate cohesion therefore supports dynamic, pore-spanning FG domain assemblies that retain brush-like extensibility^72^. This observation reinforces that cohesive interactions and entropic fluctuations can coexist in an IDP assembly, consistent with a liquid-like environment^13,19,21^. Such extensibility is likely important for maintaining a continuous transport barrier across the pore, whereas overly strong cohesion results in barrier collapse as seen with unmodified Nup98^FG^. Indeed, the pronounced compaction of Nup98^FG^ further suggests that *O*-GlcNAcylation may counter excessive cohesion to maintain a more extended and transport-competent conformation within the native NPC. Accordingly, the dynamic fluctuations of Nup62 suggest that attractive Nup–Nup interactions are continuously balanced by steric repulsion and conformational entropy, while being further modulated by transient intermolecular contacts, including multivalent NTR–FG interactions.

The expandable NuPODs mark a first step of building NPC mimics with multiple dilatory states. Powered by the unparalleled programmability of DNA origami, these reconstituted systems feature well-defined geometry, mechanics, Nup positioning and stoichiometry, and importantly, the ability to be re-engineered to approach the complexity of native NPCs, thus providing an enabling platform for studying the mechanical and biochemical interactions that govern the NPC’s barrier-transporter functions. For example, in this work we loaded only one type of Nup into each NuPOD via the same inward-facing DNA handles to compare the Nups’ cohesiveness. In the future, it would be useful to attach Nups (especially those localized to the cytoplasmic filaments and nuclear basket) according to their native arrangement in the NPC to understand how altered spatial confinement affects FG-Nups’ molecular interactions. Stacking multiple expandable DNA rings together^30,31^ would form a deeper channel that more closely resembles the depth of the NPC transport channel and accommodates multiple Nup species. This would enable the study of the collective behaviors of different Nup species in proximity, including those that naturally form complexes (e.g., Nup62·Nup58·Nup54, Nsp1·Nup49·Nup57, Nup88·Nup62·Nup214)^73–75^. Furthermore, coupling expandable NuPODs to a closed compartment^53,76–78^ would allow the investigation of cargo transport through an elastic channel lined by FG-Nups, shedding light on how bulk molecular translocation affects and is affected by NPC dilation. Lastly, although we primarily used negative-stain TEM in this study, which limits our spatial resolution to ∼2 nm (i.e., we considered rings the same size if their diameters differ by <2 nm regardless of statistical significance), the NuPODs are compatible with a multitude of analytical methods that allow for greater resolution. Similarly, in addition to HS-AFM, one can apply other techniques such as FLIM-FRET and MINFLUX to monitor the dynamics of FG-Nups and NTRs in a near-native environment.

Several DNA nanostructures with reconfigurable components have been developed for mechanically manipulating proteins^79,80^ and measuring their interaction strengths^81^. So far, the applications of these DNA devices have been limited to studying the unfolding of structured proteins and the pairwise interactions between a protein of interest and its binding partner. The expandable DNA origami ring extends the capabilities of DNA-based nanomechanical tools to analyzing multivalent IDP interactions, which are fundamental in many cellular processes^82^. Beyond FG-Nups, the platform can be retooled to sense and apply forces acting in multiple directions on other biomolecules, including proteins, nucleic acids, lipids, and their assemblages, paving the way to studying mechanically modulated molecular systems and creating synthetic analogs with comparable dynamics and function.

## Acknowledgements

C.M and A.A. thank Henry de Vries and Patrick Onck for their help with validating the ARBD implementation of their coarse-grained model.

## Author contributions

E.C. and C.L. conceived and designed the project. E.C., L.B., L.E.K., R.J.M., D.S., K.Z., Z.W., Q.F., S.C., and Q.S. performed experiments. C.M. performed MD simulation. E.C., C.M., L.B., and L.E.K. analyzed data. E.C., C.P.L., Y.X., R.Y.H.L., A.A., and C.L. interpreted data. E.C., C.M., L.B., and C.L. prepared the manuscript. All authors participated in the discussions, and reviewed and approved the manuscript.

## Data and materials availability

All data needed to evaluate the conclusions in the paper are present in the paper and/or the Supplementary Materials.

### Supplementary Materials

Supplementary Note; Figs. S1 to S33; Tables S1 to S4; Movies S1 to S5; Energy and force model scripts; NuPOD simulation scripts.

## Materials and methods

### Cloning and expression

Nup62 (aa 1–522), Nup98^FG^ (aa 1–498), Nup214^C-term^ ^FG^ (aa 1912–2090) from *Homo sapiens*, together with Nsp1^FG^ (aa 2–603) from *Saccharomyces cerevisiae*, were cloned into a pET-28a-derived vector (Novagen) as 10×His-MBP-SUMO-Nup-SNAP constructs; Nup153^FG^ (aa 896–1475) was cloned into the same vector as a 10×His-SNAP-MBP-SUMO-Nup construct. All Nups were expressed in BL21-Gold (DE3) cells, with the exception of Nup62, which was expressed in LOBSTR cells. Cells were grown in Terrific Broth at 37°C in a shaking incubator until they reached an optical density at 600 nm (OD_600_) of 0.8–1.0. Protein expression was then induced with 1 mM IPTG for 4 h at 20°C. Importin β1 (*H. sapiens*) and Kap95 (*S. cerevisiae*) were cloned into pETM-11 and pPEP-TEV vectors, respectively, each with an N-terminal 6×His tag. Both nuclear transport receptors (NTRs) were expressed in BL21-Gold (DE3) cells to an OD_600_ of 0.8–1.0, followed by induction with 0.5 mM IPTG at 15°C overnight. *O*-GlcNAc transferase (OGT) was obtained from Addgene (plasmid #190821) and was expressed in BL21-Gold (DE3) cells to an OD_600_ of 0.8–1.0, followed by induction with 0.2 mM IPTG at 15°C for 24 h. All cell pellets were collected by centrifugation at 4,500 r.p.m. for 15 min using a JS-5.0 rotor (Beckman Coulter) and stored at −80°C until use.

### Protein purification

Cell pellets expressing Nsp1, Nup214, Impβ1, Kap95, and OGT were resuspended in lysis buffer (50 mM Tris-HCl, 300 mM NaCl, 0.1 mM PMSF, 1×Roche cOmplete protease inhibitors, pH 8.0) and lysed in a homogenizer (EmulsiFlex-C3, Avestin). Whole-cell lysates were then spun at 35,000 r.p.m. for 45 min with a Type 45 Ti rotor (Beckman Coulter). Subsequently, the supernatant was decanted and filtered through a 0.45-µm-pore-size Millex syringe filter. The resulting filtered lysate was applied to a 5 mL HisTrap column (Cytiva) on an ÄKTA system (Cytiva) at a flow rate of 1 mL min^−1^. The filtrate loaded to the column was rinsed with a wash buffer (50 mM Tris-HCl, 300 mM NaCl, 25 mM imidazole, pH 8.0) and eluted with gradient buffer (50 mM Tris-HCl, 300 mM NaCl, 25–500 mM imidazole, pH 8.0). For Nup62, Nup153 and Nup98-expressing cell pellets, the same procedure was followed except that all buffers (lysis buffer, wash buffer and gradient buffer) contained additional 2 M urea and 300 mM arginine. Impβ1, Kap95, and OGT were further purified by size exclusion chromatography using a Superdex 200 10/300 GL column (Cytiva) and an aqueous mobile phase (50 mM Tris-HCl, 300 mM NaCl, pH 8.0). Nup214 was further purified using a 5 mL MBPTrap column (Cytiva) and HiLoad 16/600 Superdex 75 pg column (Cytiva). Purified proteins were validated using SDS-polyacrylamide gel electrophoresis (SDS-PAGE) and were flash-frozen in liquid nitrogen and stored at −80°C until use. All the Nups were subjected to DNA conjugation prior to further purification (see “Protein-DNA conjugation” section).

### DNA origami design and preparation

DNA-origami structures were designed using caDNAno (staple strand sequences listed in **Table S3**). ssDNA handles were extended from the 3’ ends of selected staple strands at the positions indicated in **Fig. S1**. The handle sequences are 5’-AAATTATCTACCACAACTCAC-3’ (for tethering Nup153), 5’-CTACCATCTCTCCTAAACTCA-3’ (for tethering Nsp1, Nup62, Nup214, and Nup98), and 5’-CTATCACCTTCTAACAATTCAACCTACTAACA-3’ (for tethering Nup214). To create 53-bp or 64-bp tethers for Nup214, additional extender strands were used (**Fig. S8, Table S4)**.

The expandable ring was assembled as a homo-tetramer in a one-pot reaction. The M13mp18 bacteriophage-derived circular ssDNA scaffold strand (8064 base) was combined with all staple oligonucleotides (Integrated DNA Technologies) in a molar ratio of 1:6 in a 1×TE (5 mM Tris-HCl, 1 mM EDTA, pH 8.0), 10 mM MgCl_2_ buffer using an 85–25 °C annealing program over 36 h. The Free ring is the default assembly product; Open, Hinged, and Closed configurations were formed by including the corresponding reconfiguration oligonucleotides during annealing or by post-assembly addition at room temperature. Excess staples were removed by polyethylene glycol (PEG) fractionation^83^. The rings were incubated in a solution containing 4% (w/v) PEG 8,000 (VWR), 500 mM NaCl, 1×TE and 10 mM MgCl_2_, pH 8.0 for 30 min and centrifuged at 15,000 × g for 30 min at 4 °C. The pellets were then resuspended in 1×TE, 10 mM MgCl_2_, pH 8.0 buffer. For structural characterization, PEG-purified DNA rings were subjected to rate-zonal centrifugation^84^ on a quasi-linear 15–45% glycerol gradient in a 1×TE, 10 mM MgCl_2_, pH 8.0 buffer in an SW 55 Ti rotor (Beckman Coulter) at 48,000 r.p.m. at 4 °C for 35 min. Fractions containing the Nup-free rings were identified by agarose gel electrophoresis and stored at −20 °C. To form NuPODs, PEG-purified rings were first modified with FG-Nups prior to rate-zonal centrifugation (see “Anchoring Nups to the DNA-origami ring” section). Fractions containing the NuPODs were identified by SDS-agarose gel electrophoresis and stored at 4°C for no longer than one week.

### Protein-DNA conjugation

5’-amine labeled DNA anti-handle (Integrated DNA Technologies) were ethanol precipitated to remove free amine groups and resuspended in Milli-Q water to a final concentration of 2 mM. The crosslinker benzylguanine (BG)-GLA-NHS (New England Biolabs) was dissolved in DMSO at 20 mM. Amine-labeled anti-handles and BG-GLA-NHS were mixed in a 1:20 ratio in a 67 mM HEPES, pH 8.5 buffer, and incubated at room temperature for 1 h. BG-DNA was then purified from excess crosslinkers by ethanol precipitation. The BG-DNA pellets were dried and stored at −20 °C until use.

BG-DNA pellets were resuspended in Milli-Q water and mixed with purified Nups at a final concentration of 24 µM BG-DNA and 5–10 µM Nup. For Nup214, the conjugation buffer contained 50 mM Tris-HCl, 300 mM NaCl, 1 mM DTT, pH 8.0. Nsp1 conjugation reaction additionally contained ∼400 mM imidazole. Nup62, Nup153, and Nup98 reactions additionally contained ∼400 mM imidazole, 2 M urea, and 300 mM arginine. Reactions took place overnight at 4°C. Unreacted DNA was subsequently removed by size exclusion chromatography using a Superdex 200 10/300 GL column (Cytiva) with an aqueous mobile phase (50 mM Tris-HCl, 300 mM NaCl, pH 8.0). Conjugation efficiency was verified through SDS-PAGE followed by Coomassie blue staining (Thermo Fisher Scientific).

### Anchoring Nups to the DNA-origami ring

Anti-handle-conjugated Nups were added to handle-displaying DNA-origami rings at 2:1 anti-handle:handle ratio (for example, 5 nM ring × 48 handles/ring × 2 = 480 nM Nup-anti-handle) in the hybridization buffer (1×TE, 150 mM NaCl, 10 mM MgCl_2_, pH 8.0) and incubated at room temperature for 2 h. The resulting NuPODs were purified by rate-zonal centrifugation through a 15–45% glycerol gradient in the hybridization buffer, using the rotor and centrifugation conditions described above.

### TEV cleavage of NuPODs

To remove MBP-SUMO tag from Nup62 and Nsp1^FG^, assembled NuPODs were treated with 0.25 U µL^-1^ TEV protease (Promega, Cat. No. V6101) in a reaction mixture containing 1×ProTEV Buffer (Promega) at 30°C for 2 h. NuPODs were subsequently purified by rate-zonal centrifugation as described above. Removal of the MBP-SUMO tags was verified by SDS-PAGE and Western blot (**Fig. S12**).

### SDS-agarose gel electrophoresis

Samples were loaded in an SDS-agarose gel (1% agarose in 0.5×TBE, 10 mM MgCl_2_, 0.05% SDS, pH 8.0). Electrophoresis was performed at 65 V for 120 min in 0.5×TBE buffer containing 10 mM MgCl_2_ and 0.05% SDS, pH 8.0 using an Owl EasyCast B2 Mini Gel Electrophoresis Systems (Thermo Scientific). Following electrophoresis, gels were soaked in deionized H_2_O and shaken for 2 h to remove SDS, and subsequently stained overnight at room temperature in SYBR Gold solution (Invitrogen, 10,000× dilution in H_2_O). Gels were then imaged using a Typhoon FLA scanner (v1.0 software).

### SDS-PAGE

All SDS-PAGE experiments were performed using 4–12% acrylamide NuPAGE Bis-Tris gel (Thermo Fisher Scientific). Samples were boiled in 1×Laemmli sample buffer (Thermo Fisher Scientific) at 90 °C for 5 min before loading to the gels. Electrophoresis was carried out for 90 min at a constant voltage of 120 V in MES-SDS buffer (Thermo Fisher Scientific) using an XCell SureLock Mini-Cell system (Thermo Fisher Scientific). Gels were stained with Coomassie Blue. Gel images were acquired using the ChemiDoc MP imaging system (Bio-Rad) and analyzed in ImageJ v1.53k.

### *In vitro O*-GlcNAcylation of Nup-anti-handles

Purified Nup (Nup62, Nup153, Nup98)-anti-handles were *O*-GlcNAcylated using recombinant OGT and UDP-GlcNAc (MilliporeSigma, U4375). Reaction mixtures contained 1 µM Nup-anti-handle, 0.2 µM OGT, 5 mM UDP-GlcNAc in 50 mM Tris-HCl, 150 mM NaCl, 12.5 mM MgCl_2_, 1 mM DTT, pH 7.5 and were incubated at 28 °C for 2 h with gentle agitation. *O*-GlcNAcylated Nup-anti-handles were subsequently mixed with DNA-origami rings, followed by rate-zonal centrifugation as described above.

### NuPOD incubation with NTRs

Unless otherwise noted, purified NuPODs were incubated with 1 µM NTR (Impβ1 or Kap95) at room temperature for 1 h before imaged by negative-stain TEM without purification (e.g., **Fig. 5**). Alternatively, 1 µM NTR was introduced immediately after anchoring Nups to the DNA-origami rings, followed by incubation at room temperature for 1 h; the mixture was then subjected to rate-zonal centrifugation through a glycerol gradient supplemented with 1 µM NTR, using centrifugation conditions described above.

### Western blot

Protein samples were separated by SDS-PAGE and transferred to PVDF membranes. Membranes were blocked with EveryBlot Blocking Buffer (Bio-Rad) for 15 min at room temperature and incubated overnight at 4 °C with primary antibodies diluted in EveryBlot Blocking Buffer. Primary antibodies included an anti-*O*-GlcNAc monoclonal antibody (clone RL2; Abcam, Cat. No. ab2739) and an anti-Nup62 antibody (BD Transduction Laboratories, Cat. No. 610497). After washing with TBST, membranes were incubated with HRP-conjugated secondary antibodies for 1 h at room temperature. Immunoreactive bands were detected by enhanced chemiluminescence (ECL) and imaged with a ChemiDoc imaging system. Western blot images were analyzed in ImageJ.

### Negative-stain TEM

NuPOD samples were applied to glow-discharged 400-mesh Formvar/carbon-coated copper grids (Ted Pella) and stained with 2% uranyl formate (Electron Microscopy Sciences). Images were acquired on a JEOL JEM-1400 Plus transmission electron microscope operated at 80 kV and equipped with a bottom-mounted 4k × 3k CCD camera (Advanced Microscopy Technologies). Data were collected using AMT Image Capture Engine v602.

### Measurement of inner-ring diameter

Negative-stain electron micrographs were analyzed in ImageJ. Individual DNA ring or NuPOD particles exhibiting a top view were selected for analysis. The inner-ring boundary and the inner boundary of the outer octagonal frame were manually outlined, and their enclosed areas were measured. The inner-ring diameter was calculated as the diameter of a circle with an equivalent area. To correct for particle-to-particle variations in the projected size arising from image distortions and slight differences in particle orientation, the area enclosed by the inner octagonal frame was converted to the distance across flats of an equivalent regular octagon. A normalization factor was then calculated by comparing the measured across-flats distance with the designed value of 91 nm. The inner-ring diameter of each particle was multiplied by this normalization factor, and the resulting value was reported as the inner-ring diameter (*d*_in_). For particle averaging, individual particles were automatically picked using EMAN2 v2.9. Reference-free two-dimensional class averages were generated using standard image-processing workflow implemented in EMAN2.

## Statistical analysis

Statistical analyses were performed using GraphPad Prism v10.4.0. Unless otherwise, indicated, data are presented as mean ± s.d. Statistical significance between two groups was assessed using unpaired two-tailed Student’s *t*-tests assuming equal variances. Sample sizes are provided in the corresponding Supplementary Figures. Differences were considered statistically significant at *P* < 0.05.

### SPR experimental setup and analysis

The binding affinity and kinetics between different FG-Nups were measured in PBS buffer using Biacore T200 (Cytiva). FG-domain of Nup62^F→S^ (MW = 22 kDa), Nup62 (23 kDa) and Nup153 (62 kDa) were modified with cysteine at their C- or N-terminus to immobilize Nups on the gold surface of SIA kit Au SPR sensor chip (Cytiva, BR100405) in the flow channels 2, 3 and 4, respectively. The reference channel (flow channel 1) was blocked with 1 mM PUT3, i.e. (1-mercapto-11-undecyl)-(ethylene glycol)_3_, to prevent any nonspecific binding of analyte there. Then an increasing concentration (dilutions: 0.153, 0.312, 0.625, 1.25, 2.5, 5, 10, 20 µM) of MBP-SUMO-Nup62^F→S^-SNAP was injected in all 4 channels probing its binding to the immobilized FG-Nups followed by regeneration using 0.1 M NaOH. In the following cycles an increasing concentration of MBP-SUMO-Nup62-SNAP (dilutions: 15.3, 31.2, 62.5, 125, 250, 500, 1000, 2000 nM) was titrated. The obtained binding sensograms were analyzed using our custom scripts written for Igor Pro and MATLAB^85^. The affinity was obtained by fitting the equilibrium binding data to a one-component Langmuir isotherm. The kinetic constants were determined by fitting binding sensograms to kinetic equations as described previously^62,85^. The resulting kinetic maps were generated using a Python script. The height changes were analyzed using procedures described previously^86,87^. The grafting distance of the immobilized ligand was estimated using commonly accepted approximation that 1300 RU corresponds to the binding of 1 ng mm^-2^ of protein on a planar surface^62,86,88^:

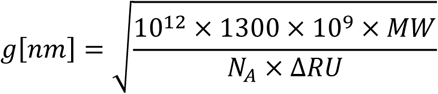

where *MW* is molecular weight of ligand molecule, *N*_A_ is an Avogadro constant, Δ*RU* is a change of the SPR signal due to ligand binding. The obtained grafting distances were similar to the spacing between adjacent FG-Nup anchoring sites in NuPODs.

### HS-AFM measurements

#### Sample preparation

Supported lipid bilayers (SLBs) were prepared as previously described^27^. Dipalmitoylphosphatidylcholine (Avanti Polar Lipids) and didodecyldimethylammonium bromide (Avanti Polar Lipids) were dissolved in chloroform and mixed at a 3:1 molar ratio. After evaporation of the solvent under a nitrogen stream followed by vacuum desiccation, the dried lipid film was rehydrated in Milli-Q water to a final lipid concentration of ∼1 mg mL^-1^. The suspension was sonicated at ∼65 °C for 15 min and extruded through a 100-nm track-etched polycarbonate membrane (Avanti Polar Lipids) maintained at ∼65 °C for at least 21 passes to generate small unilamellar vesicles (SUVs).

SLBs were formed by incubating SUVs on freshly cleaved mica mounted on an HS-AFM sample stage at ∼65 °C for 20 min, followed by gradual cooling to room temperature. Excess vesicles were removed by rinsing with Milli-Q water, and the sample was equilibrated by four sequential exchanges with imaging buffer (10 mM Tris-HCl, 0.1 mM EDTA, 10 mM MgCl₂, 150 mM NaCl, pH 8.0). NuPODs were subsequently deposited onto the SLB, followed by incubation with Impβ1 (100 nM or 1 µM, as indicated) for 30 min before HS-AFM imaging.

#### AFM data acquisition

HS-AFM measurements and data processing were performed as previously described^19^. Images were acquired on an HS-AFM 1.0 system (RIBM) operating in tapping mode with a standard scanner. QUANTUM-AC10-SuperSharp and QUANTUM-AC10-SuperSharp Enhanced cantilevers (NanoTools GmbH) with a nominal spring constant of 0.1 N m^-1^, resonant frequency of ∼0.5 MHz in water and a tip radius of ≤ 2 nm was used throughout. The free oscillation amplitude was maintained at 2–3 nm, and the set-point amplitude was adjusted to 80–90% of the free oscillation amplitude, as described previously^89,90^. For HS-AFM line-scanning (HS-AFM-LS), the slow-scan axis was disabled and only the fast scan axis was used, as described previously^91^. Line scans were acquired over 80 pixels with a line acquisition time of 1.875 ms.

#### Image processing and analysis

HS-AFM images were corrected for lateral drift and converted to TIFF format using in-house Python software, as previously described^19,20^. Subsequent image processing and analysis were performed in ImageJ and custom Python scripts. Images were corrected for sample tilt by first-order polynomial plane flattening, and heights were normalized to the NuPOD by subtracting the mean NuPOD height. Zoomed-in images were additionally processed using a two-dimensional Gaussian filter (standard deviation = 1 px).

Height profiles were extracted in ImageJ along two horizontal and two vertical line regions of interest passing through the pore center. Profiles were normalized by subtracting the maximum height of the outer octagonal frame (30–60 nm from the pore center). Mean height profiles and standard deviations were calculated by pooling all scan directions and particles within each condition. Inner-ring diameters were determined as twice the distance from the pore center to the detected inner-ring peak and averaged over the four scan directions for each particle.

#### HS-AFM-LS data analysis

HS-AFM-LS kymographs were processed using custom Python scripts. Drift and fast-scan-axis tilt were corrected, and heights were referenced to the NuPOD where the NuPOD surface was visible within the same kymograph. Periodic electronic noise was removed by fast Fourier transform filtering. The position of the central density was determined for each scan line and expressed as the signed distance from the pore center. Position distributions pooled from all kymographs were visualized as histograms. Autocorrelation functions were calculated for individual scan lines and fitted with a single-exponential decay to obtain the position-dependent decay time (τ). Fits that did not satisfy predefined quality criteria were excluded from further analysis. Mean τ profiles were calculated by averaging all retained values at each radial position. To quantify the dynamics of the central plug-like protein mass, the mean τ for each pore was calculated from all retained values within |x| ≤ 10 nm of the pore center.

### Coarse-grained equilibrium molecular dynamics simulations

All MD simulations were performed using ARBD^92^, an in-house developed GPU-accelerated code. Empty NuPODs were constructed using a Python script to read in the caDNAno design file and generate four quarter-ring mrDNA^63^ domains that were subsequently rotated and connected according to the sticky ends in the design file to form a complete NuPOD. Beads were then generated using the mrDNA package with a maximum of 4 bp (or 4-base ssDNA) represented by each bead. Each of the four designs, Open, Free, Hinged, and Closed were simulated for 20 µs using a Langevin thermostat with 295 K temperature with a 200 fs timestep and trajectory frames output every 100,000 steps. A 10 nm nonbonded interaction pair list was recalculated every 250 steps, with interactions smoothly truncated beyond 5 nm.

Simulations of NuPODs loaded with FG-Nups were performed by constructing a Hinged NuPOD as described above before adding, at each of 48 anchor points, a single dsDNA helix with a flexible crossover connection to the origami scaffold on one end and single bond to a one-amino-acid-per-bead model of an FG-Nup peptide at the other. The peptide was initially stretched linearly across the pore in such a way as to avoid clashes with other peptides. The peptide was represented by updating our previous ARBD implementation^93^ of the FG-Nup model developed and described by the Onck group^22^ to incorporate their recent improvements^12^. The folded domains of each peptide (MBP, SUMO and SNAP) were maintained in their initial PDB-derived configurations using an elastic network with harmonic bonds restraining all bead pairs within 1.2 nm (*k*_spring_ = 956 kcal mol^-1^ nm^-2^) and a reduction of nonbonded interactions as previously described by the Onck group^12^. For nonbonded interactions between DNA and peptide beads, we employed the screened Coulomb and 8–6 Lennard-Jones nonbonded interaction potential from the peptide model, but with *ε* = 0.1 kcal mol^-1^ and *σ* = 1.6 nm. The bond between the DNA linker and tethering bead of the peptide was represented by a harmonic potential with a 9.8 Å rest length, with *k*_spring_ = 1000 kcal mol^-1^ nm^-2^, a maximum restoring force of 100 kcal mol^-1^ nm^-1^, and with nonbonded exclusions applied. The mrDNA model employs a large damping coefficient (bead size dependent; typically ∼7000 ns^-1^ in this model) whereas the Onck model employs a small 20 ns^-1^ coefficient to promote rapid relaxation. To ensure the relaxation timescales of the models were not badly mismatched, we applied damping coefficients of 60 ns^-1^ and 400 ns^-1^ to peptide and DNA beads, respectively, while using a 20 fs timestep. Trajectory coordinates were recorded every 10,000 steps. All other simulation parameters remained unchanged compared to the Nup-free NuPOD simulations described above. Two replicas of each system were simulated for at least 7 µs and as much as 35 µs each.

Analysis of the inner ring diameter was performed by calculating the radius of gyration of the constituent beads. The mass density of the beads was extracted from a class average obtained using Relion^94^ from images of the mass density of configurations sampled from the first replica uniformly between 2.8 and 22 µs or the end of the trajectory, whichever is shorter. Each configuration was subject to a random rotation about its planar axis and random tilt by an angle sampled from a ±8 degree normal distribution. The structured domains of the peptides were neglected during the calculation.

### Coarse-grained free energy perturbation simulations

FEP^95,96^ simulations of each Nup-containing model were performed to estimate the total interaction energy between a single Nup and the rest of the mesh using four randomly selected configurations from the first replica of equilibrium trajectories described above. Within each configuration, the interactions of a single randomly chosen peptide with the other peptides were made to gradually disappear and reappear in accordance with the theory of FEP, and with the selection process detailed below. The simulations employed a weak harmonic restraint (*k*_spring_ = 0.1 kcal mol^-1^ nm^-2^) with rest length of zero applied between the geometric center of the selected Nup and the center of the DNA that was intended to prevent the chosen Nup from diffusing away from the central mesh. Hence, to account for effect of this restraint, we estimated the distribution of distances between each Nup and the DNA from the equilibration trajectories and weighted the probability of selecting a given Nup according to the Boltzmann distribution for the restraint potential.

A development feature of ARBD known as switching potentials was used to facilitate the FEP procedure. Briefly, the feature allows each particle to change its type (determining nonbonded interactions) smoothly according to a predefined schedule of control values. By introducing a set of peptide types identical to the original peptide types, but with zero nonbonded interaction between the two sets (while retaining intramolecular nonbonded interactions), it is possible to transform the system from a state where all Nups interact to one in which a single chain interacts only with itself. The FEP control parameter was made to vary gradually from zero (fully interacting) to one (non-interacting) in 28 steps according to the following schedule: 0, 0.02, 0.05, 0.1, 0.13, 0.16, 0.2, 0.3, 0.4, 0.5, 0.6, 0.64, 0.68, 0.72, 0.76, 0.8, 0.85, 0.9, 0.95, 0.96, 0.97, 0.975, 0.98, 0.985, 0.99, 0.995, 0.9975, 1. At the end of the procedure, the schedule was reversed. At each value of the control parameter, 200 samples were collected with 1000 steps between each sample. Switching between values was done smoothly over 1000 steps. Other than the initial condition, the added restraint, the particle type switching, and the higher 1000 step output period, the simulations were performed as described above for equilibration simulations. A fork of ARBD was used to post-process each trajectory, extracting the energy of each configuration for Hamiltonians using the current and next control parameter value for use in the FEP calculation. The multistate Bennet acceptance ratio (MBAR) algorithm^64^ was then used via the pymbar package to estimate the free energy cost of the transformation for each replica using both forward and reverse trajectories and a 160 step decorrelation time. The mean values and standard errors of the mean were then obtained by averaging over replicas. In addition to the FEP energy, the free energy results reported in this work incorporate smaller free energy changes associated with imposing the harmonic restraint in the initial state and removing it in the final state. The free energy associated with restraints was estimated from the distributions of peptide–DNA distances by direct evaluation of the configurational integral^97^. Bootstrapping was used to determine standard errors.

### Simulations estimating the mechanical work to expand the inner ring

The same configuration used to initialize the FEP simulations were used to probe the work required to move the anchors of the Nups from their equilibrium positions to their estimated positions in an unloaded (apo) Hinged DNA ring. Simulations were performed identically to the equilibrium simulations, except that the DNA was removed from each system and each peptide bead previously bound to the DNA was harmonically restrained about its initial position (*k*_spring_ = 10 kcal mol^-1^ nm^-2^). The displacement of each restrained bead during the trajectory along the radial direction and in the plane of the DNA scaffold reports the force the DNA ring would need to exert on the peptide to prevent further collapse of the mesh. For each Nup, we performed a relatively short 20 ns simulation to relax the mesh before moving the restraint positions radially outward by 2 Å and repeating the process until the restraint positions had shifted by an amount equal to the difference between the average radius of the Hinged Nup-free ring, as determined from our equilibrium simulations. Subsequently, all the restraint simulations were extended by an additional 200 ns. The work required to expand the ring was determined by trapezoidal integration of the average force.

