## Supplementary Materials for "Expandable DNA-origami rings as a nanomechanical platform for studying disordered nucleoporins"

Eason Cao *et al.*

**This PDF file includes:**

Supplementary Note
Figs. S1 to S33
Tables S1 to S4
Legend for movies S1 to S5
References

### SUPPLEMENTARY NOTE

#### Physical model of ring configurational energy (*E*) and effective radial force ($F_{radial}$)

**Geometric parameterization of ring expansion.**

The model is parameterized by $\Delta r$, the outward radial displacement of each inner-ring arc from the fully closed state ($\Delta r = 0$ corresponds to the fully closed state, with $d_{in}$ = 46.5 nm. The maximum value of $\Delta r$ depends on the ring configuration and is defined below for each model.). All energy and force terms are expressed as functions of $\Delta r$. Because the inner ring expands as a rounded square rather than a circle – comprising four fixed quarter-circle arc segments of radius $r_0$ = 23.25 nm connected by four straight inner-ring-segment gaps of length $\sqrt{2}\Delta r$ – the reported inner-ring diameter corresponds to that of a circle with equivalent enclosed area (*S*):

$$d_{\text{in}} = 2\sqrt{\frac{S(\Delta r)}{\pi}}, \quad S(\Delta r) = \pi r_0^2 + 4r_0\sqrt{2}\Delta r + 2\Delta r^2$$

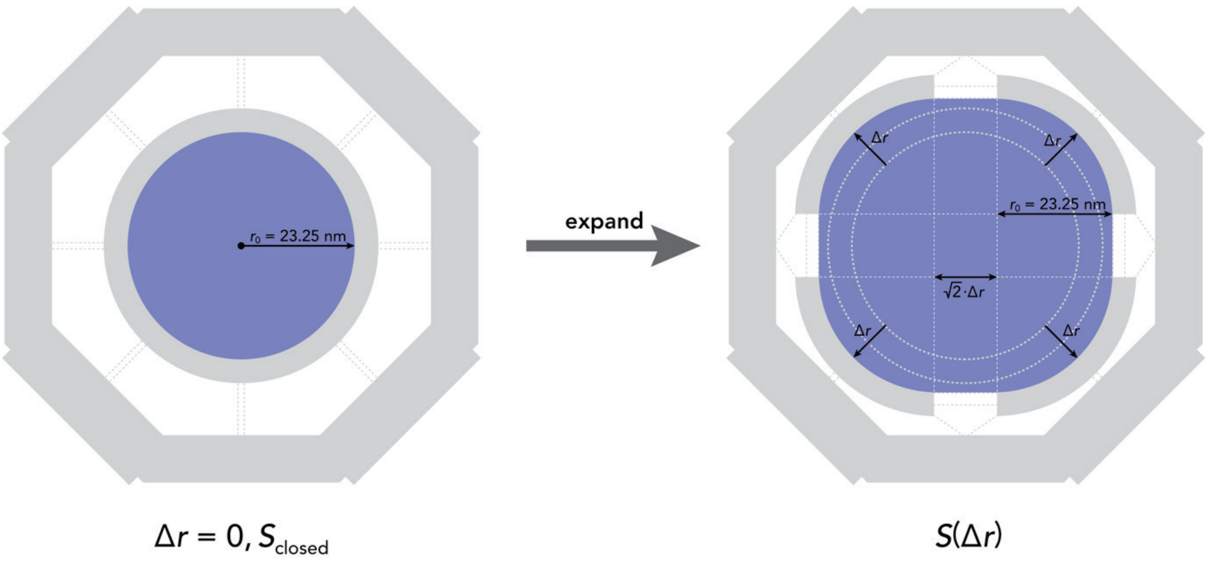


**Sign convention.**

Radial displacement $\Delta r$ is defined as positive in the outward direction from the fully closed state. Radial forces are defined as positive in the inward direction (i.e., tending to decrease $\Delta r$), such that

$$F_{\text{radial}} = \frac{dE}{d(\Delta r)}$$

**$E_{stretch}$: elastic stretching of ssDNA linkers.**

The inner ring is connected to the outer octagonal frame by two groups of geometrically distinct 51-nt ssDNA scaffold linkers ($N_1 = N_2$ = 16 linker copies per ring; contour length $L_{c1} = L_{c2}$ = 51 × 0.63 nm = 32.1 nm) whose extensions as functions of $\Delta r$ are determined by the device geometry. In the following geometry equations, all lengths are expressed in nm:

$$\mathrm{ext}_1(\Delta r) = \frac{90.6}{2} - \frac{46.5}{2} - 5 - \Delta r = 17.05 - \Delta r$$

$$\mathrm{ext}_2(\Delta r) = \sqrt{\Delta r^2 - 2 \times 17.05 \times \cos 45^\circ\ \Delta r + 17.05^2} = \sqrt{\Delta r^2 - 24.1\,\Delta r + 290.7}$$

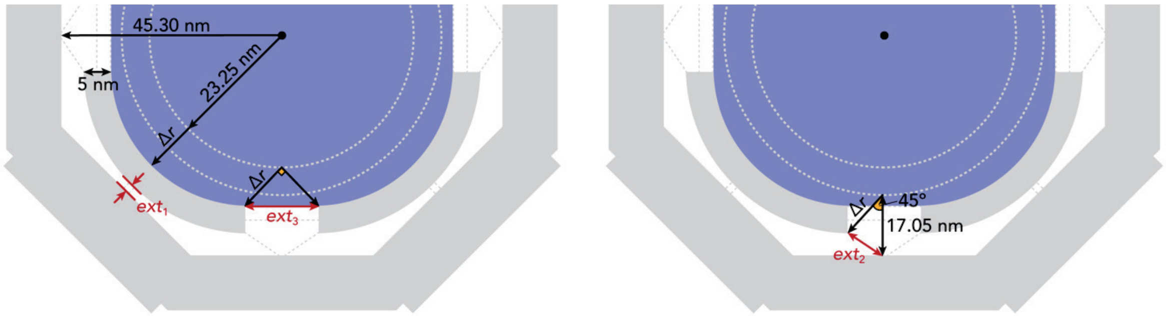


The contour length per nucleotide (0.63 nm) is a standard geometric parameter for ssDNA, whereas the persistence length ($L_{p,ss}$ = 1.05 nm) is estimated based on previously reported measurements under physiological ionic conditions[1]. The elastic energy is computed by integrating the Marko-Siggia worm-like chain (WLC) force-extension relation[2]:

$$E_{i,\mathrm{WLC}}(\Delta r) = \int_0^{\mathrm{ext}_i} F_{\mathrm{WLC}}(L)dL\,, \quad F_{\mathrm{WLC}}(L) = \frac{k_B T}{L_{p,\mathrm{ss}}}\left[\frac{1}{4\left(1 - \frac{L}{L_c}\right)^2} - \frac{1}{4} + \frac{L}{L_c}\right]$$

summed over all 16 linkers per ring for each group ($E_1$ and $E_2$). In the free ring, the 72-nt inter-segment ssDNA ($L_{c3}$ = 72 × 0.63 nm = 45.4 nm) is stretched to an end-to-end distance ($ext_3$) of $\sqrt{2}\Delta r$, contributing an additional inward elastic stretch term $E_3$ computed by the same WLC integral over $N_3$ = 16 copies. The effective radial force from each linker follows from the chain rule:

$$F_{i,\mathrm{radial}}(\Delta r) = F_{\mathrm{WLC}}(\mathrm{ext}_i)\,\frac{d(\mathrm{ext}_i)}{d(\Delta r)}$$

giving $\frac{d(\mathrm{ext}_1)}{d(\Delta r)} = -1$ (outward) and $\frac{d(\mathrm{ext}_2)}{d(\Delta r)} = \frac{\Delta r - 17.05/\sqrt{2}}{\mathrm{ext}_2} = \frac{\Delta r - 12.06}{\mathrm{ext}_2}$ (direction depends on $\Delta r$, changing sign at $\Delta r$ = 12.06 nm). For $F_3$, $\frac{d(\mathrm{ext}_3)}{d(\Delta r)} = \sqrt{2}$ (inward).

**$E_{\mathrm{bend}}$: elastic bending of inter-arc bridges.**

Hinged ring: The inter-arc bridge consists of two rigid 33-bp dsDNA arms ($L_{\mathrm{arm}}$ = 33 × 0.335 nm = 11.1 nm) flanking a soft 4-nt ssDNA pivot ($L_{\mathrm{hinge}}$ = 4 × 0.63 nm = 2.52 nm). Because $L_{p,\mathrm{ds}} \gg L_{\mathrm{arm}}$, the arms are treated as rigid rods and all angular deformation is concentrated at the pivot, modeled as an effective angular spring with stiffness $k_{\theta,\mathrm{hinged}}$:

$$k_{\theta,\text{hinged}} = \frac{k_B T L_{p,\text{ss}}}{L_{\text{hinge}}} \approx 0.42\ k_B T\ \text{rad}^{-2}$$

The bending angle $\theta_h$ is set by the inner-ring segment geometry:

$$\sqrt{2}\,\Delta r = 2L_{\text{arm}} \sin\left(\frac{\theta_h}{2}\right) \Rightarrow \theta_h = 2\arcsin\left(\frac{\sqrt{2}\,\Delta r}{2L_{\text{arm}}}\right)$$

The energy and radial force are:

$$E_{\text{bend}} = N_{\text{linkers}}\,\frac{1}{2}k_{\theta,\text{hinged}}(\theta_h - \pi)^2$$

$$F_{\text{bend,radial}} = N_{\text{linkers}}\,k_{\theta,\text{hinged}}\,(\theta_h - \pi)\,\frac{d\theta_h}{d(\Delta r)}, \quad \frac{d\theta_h}{d(\Delta r)} = \frac{\sqrt{2}}{L_{\text{arm}}\sqrt{1-\left(\frac{\sqrt{2}\,\Delta r}{2\,L_{\text{arm}}}\right)^2}}$$

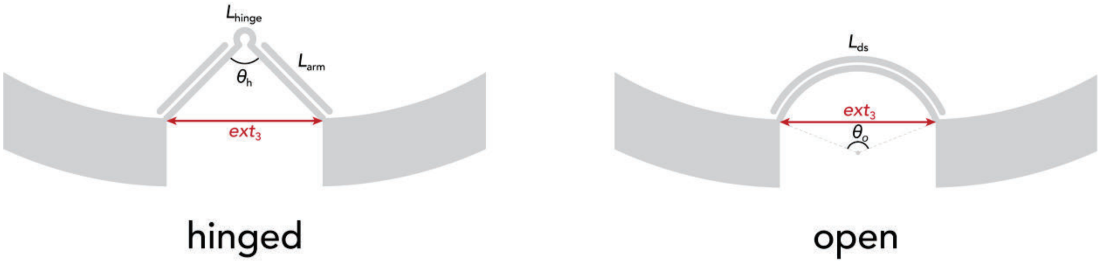


This bending description is only valid while the arcsine argument stays below 1, i.e., before the two rigid arms become fully collinear. The limit is reached at

$$\sqrt{2}\,\Delta r^* = 2L_{\text{arm}}, \quad \Delta r^* \approx 15.63\ \text{nm}$$

where $\theta_h = \pi$ and $E_{\text{bend}} = 0$. Since the Hinged ring model is evaluated beyond $\Delta r^*$, for $\Delta r > \Delta r^*$, we set the bending angle at $\theta_h = \pi$ and assign the additional gap to extension of the 4-nt ssDNA pivot itself, analogous to the ssDNA linker stretching in $E_{\text{stretch}}$:

$$\text{ext}_{\text{hinge}}(\Delta r) = \sqrt{2}\,\Delta r - 2L_{\text{arm}}, \quad \Delta r > \Delta r^*$$

$$E_{\text{hinge,stretch}}(\Delta r) = N_{\text{linkers}} \int_0^{\text{ext}_{\text{hinge}}(\Delta r)} F_{\text{WLC}}(L)\ d\text{L}$$

$$F_{\text{hinge,stretch,radial}}(\Delta r) = N_{\text{linkers}}\, F_{\text{WLC}}\big(\text{ext}_{\text{hinge}}\big)\, \frac{d\,\text{ext}_{\text{hinge}}}{d(\Delta r)}$$

The total inter-segment energy and force of the Hinged ring are then the piecewise sum of the bending and hinge-stretching terms.

Open ring: The inter-arc bridge is a single 71-bp dsDNA rod ($L_{\text{ds}}$ = 71 × 0.335 nm = 23.8 nm) modeled as a uniformly curved WLC circular arc. The total bending angle $\theta_{\text{o}}$ satisfies the chord-arc relation:

$$\sqrt{2}\,\Delta r = L_{\text{ds}}\,\frac{\sin(u)}{u}, \quad u = \frac{\theta_o}{2}$$

which is solved numerically at each $\Delta r$ using a lookup table and Newton-Raphson refinement. Because the dsDNA segment has a finite contour length, the open-ring bending model is evaluated only over the geometrically allowed range $\sqrt{2}\Delta r \leq L_{ds}$. Thus, for the open ring, the maximum radial displacement is set by $\Delta r_{max} = L_{ds}/\sqrt{2} \approx 16.82$ nm. The energy and radial force are:

$$E_{\text{bend}} = N_{\text{linkers}}\,\frac{1}{2}k_{\theta,\text{open}}\theta_o^2, \quad k_{\theta,\text{open}} = \frac{k_B T L_{p,\text{ds}}}{L_{\text{ds}}}$$

$$F_{\text{bend,radial}} = N_{\text{linkers}}\, k_{\theta,\text{open}}\,\theta_o\,\frac{d\theta_o}{d(\Delta r)}, \quad \frac{d\theta_o}{d(\Delta r)} = \frac{2\sqrt{2}u^2}{L_{\text{ds}}(u\cos u - \sin u)}$$

derived by implicit differentiation of the chord-arc relation. Since the rod straightens as $\Delta r$ increases, $d\theta/d(\Delta r) < 0$ and $F_{\text{bend}}$ is negative (outward) – the bent rod acts as a spring pushing the ring open. The effective angular stiffness of the dsDNA rod ($k_{\theta,\text{open}} \approx 2.1\ k_B T$ rad$^{-2}$) is approximately five-fold larger than the hinge pivot stiffness ($k_{\theta,\text{hinged}} \approx 0.42\ k_B T$ rad$^{-2}$), explaining why the open ring resists contraction far more than the hinged ring.

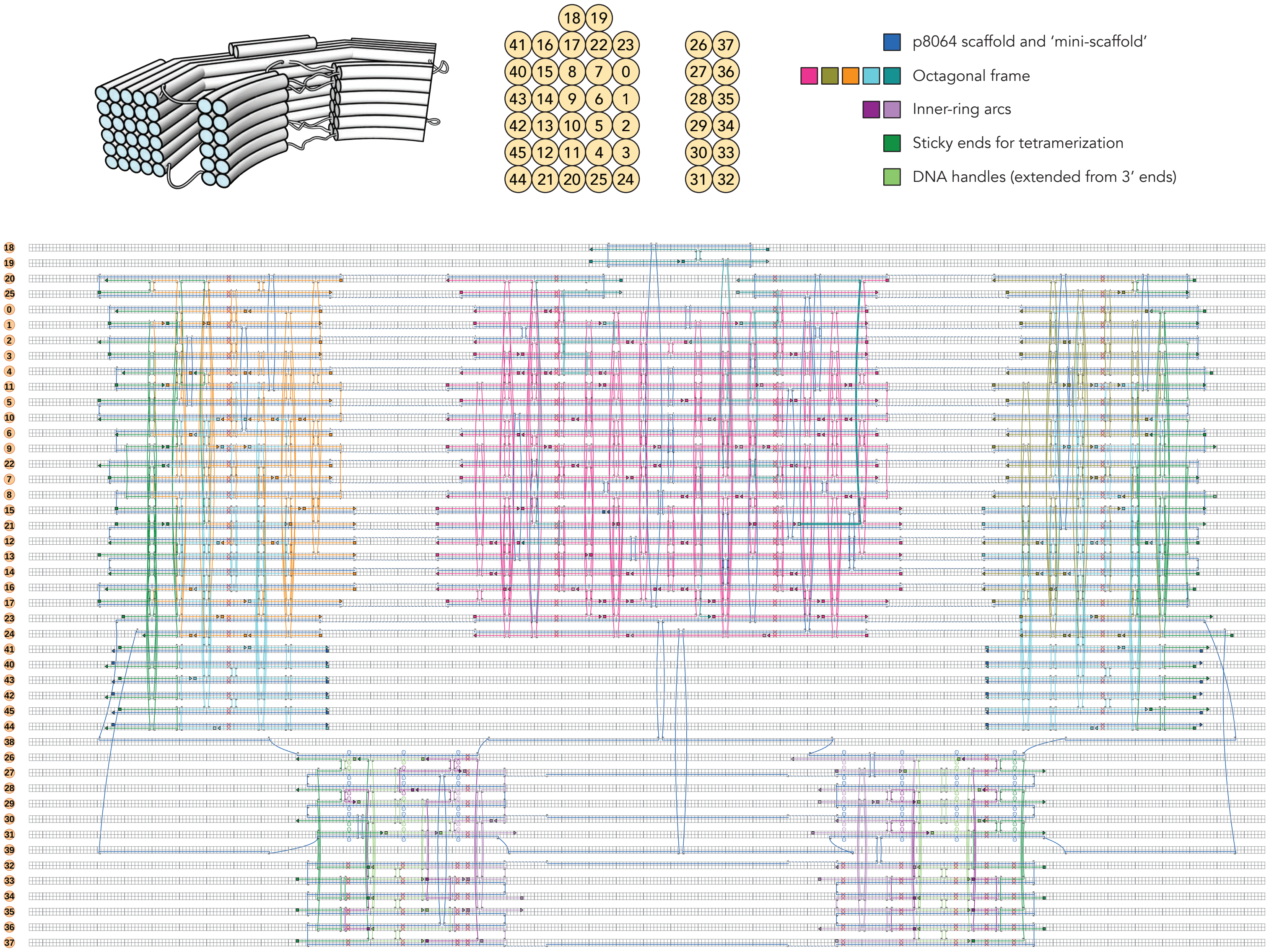


**Fig. S1** | CaDNAno design of the expandable ring. Only one of the self-tetramerizing monomers is shown. Top, schematic of the quarter-ring monomer, cross-sectional arrangement of DNA duplexes, and color scheme used in the strand diagram. Bottom, caDNAno design of the quarter ring shown as a strand diagram.

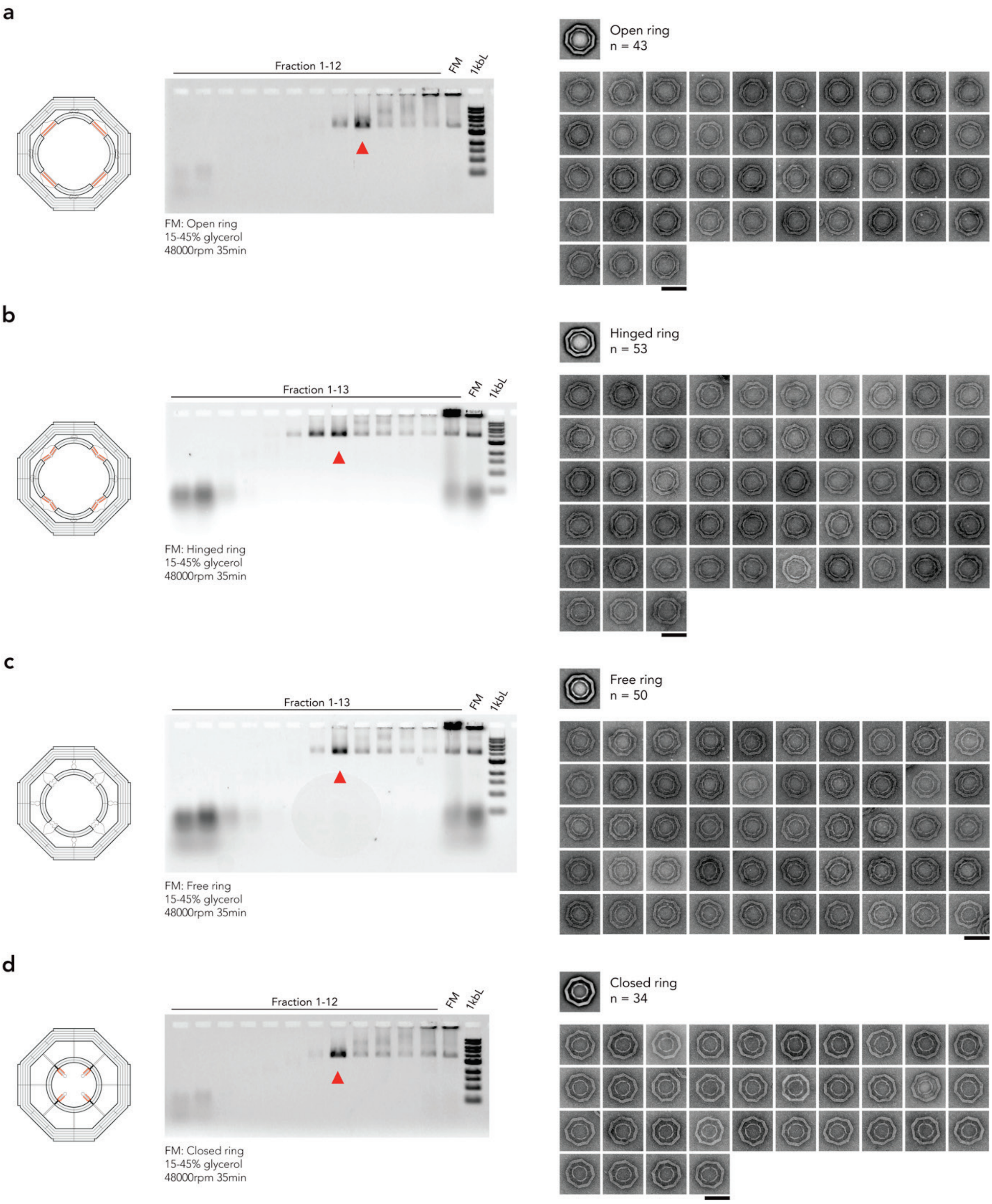


**Fig. S2 | Purification and structural characterization of Nup-free (Apo) expandable rings. a–d**, Agarose gel electrophoresis analysis of fractions collected after rate-zonal centrifugation (15–45% glycerol, 48,000 r.p.m., 35 min) of Open (**a**), Hinged (**b**), Free (**c**), and Closed (**d**) DNA rings. Red arrowheads indicate the fractions selected for negative-stain TEM analyses. Representative particle galleries of the purified DNA rings are shown on the right. FM, folding mixture; 1kbL, 1 kb DNA ladder. Scale bars, 100 nm.

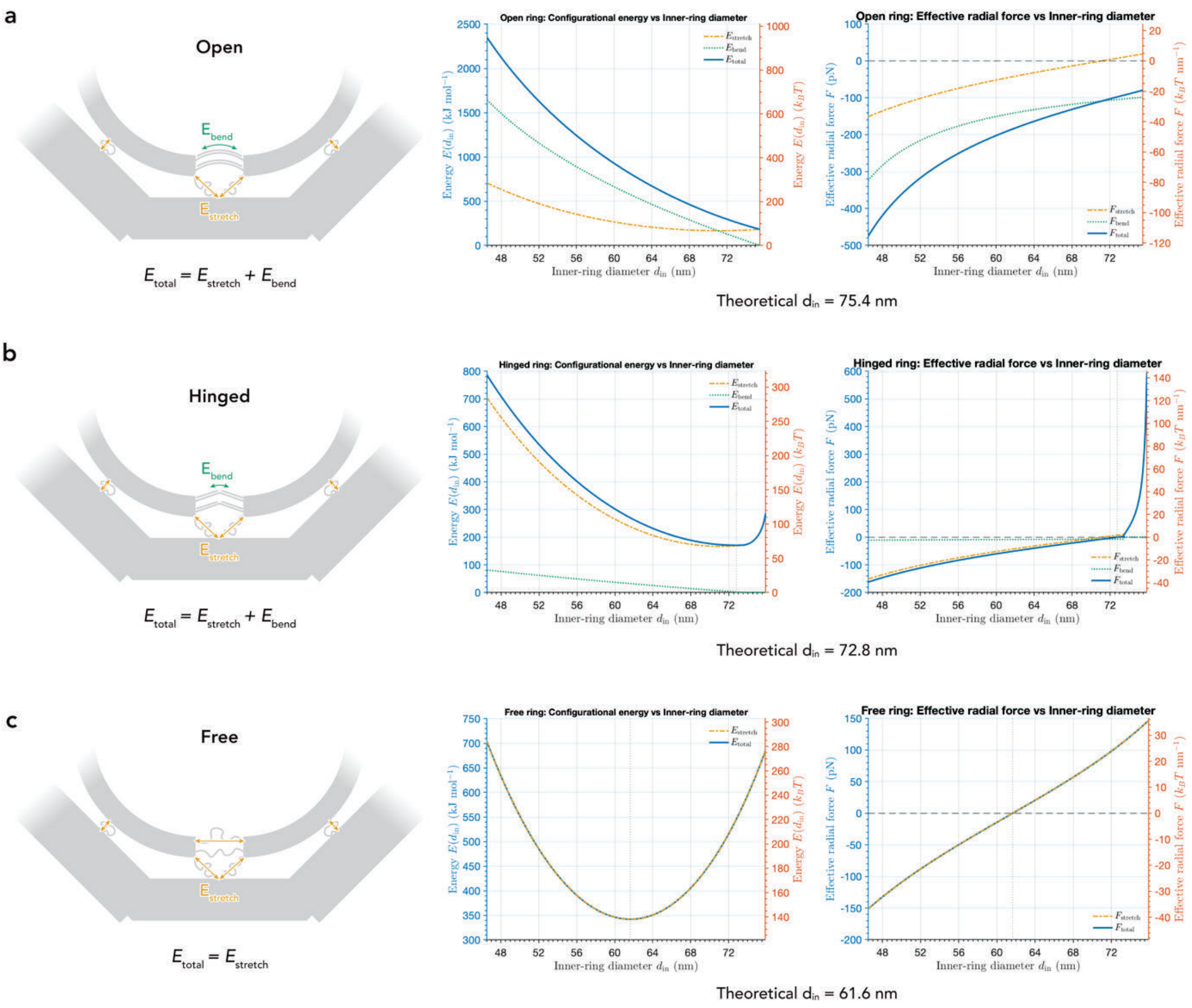


**Fig. S3 | Theoretical energy landscapes of expandable rings. a–c**, Schematic illustrations of the stretching and bending energy terms used in the theoretical model (left), calculated stretching, bending, and total configurational energies as functions of inner-ring diameter $d_{\mathrm{in}}$ (middle), and the corresponding effective radial forces (right) for the Open (**a**), Hinged (**b**), and Free (**c**) DNA rings. Theoretical equilibrium inner-ring diameters are indicated below each panel.

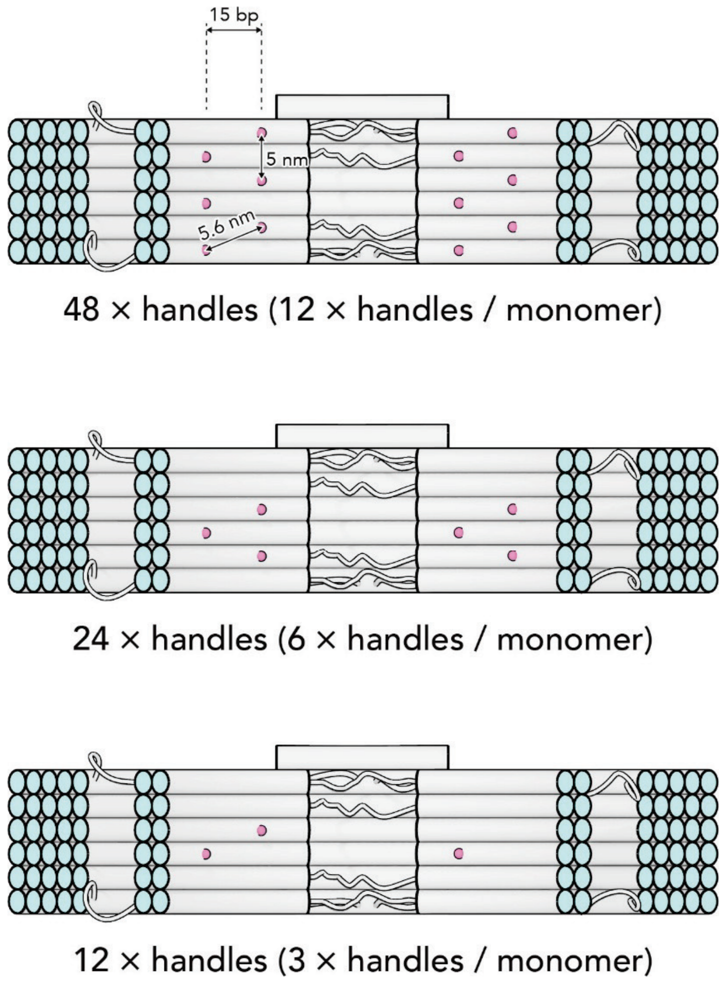


**Fig. S4 | Design of expandable rings with different handle densities.** Schematic illustration of the inner-ring arcs carrying 48, 24, or 12 DNA handles (12, 6, or 3 handles per monomer, respectively). DNA handles are spaced at ~5 or ~5.6 nm apart (15 bp along the DNA helix on alternating helices) within each inner-ring arc.

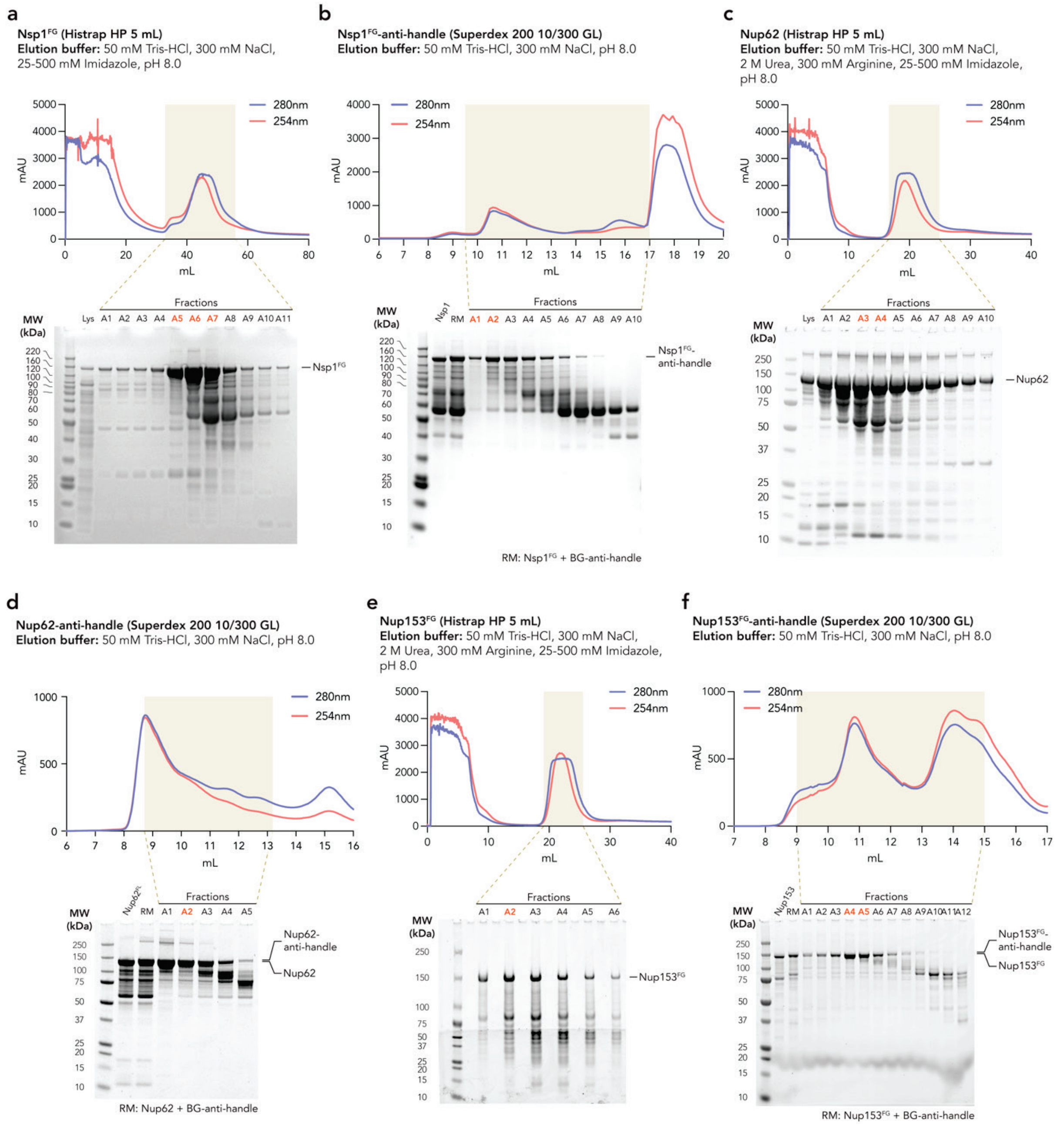


**Fig. S5 | Purification of Nsp1$^{FG}$, Nup62, and Nup153$^{FG}$ and their anti-handle conjugates. a–f**, Representative chromatograms and SDS-PAGE analyses of Nsp1$^{FG}$ purification by HisTrap affinity chromatography (**a**), purification of Nsp1$^{FG}$-anti-handle by size-exclusion chromatography (**b**), Nup62 purification by HisTrap affinity chromatography (**c**), purification of Nup62-anti-handle by size-exclusion chromatography (**d**), Nup153$^{FG}$ purification by HisTrap affinity chromatography (**e**), and purification of Nup153$^{FG}$-anti-handle by size-exclusion chromatography (**f**). Shaded regions indicate the fractions collected and analyzed by SDS-PAGE, and fractions highlighted in red were pooled for subsequent experiments. RM, reaction mixture; Lys, lysate.

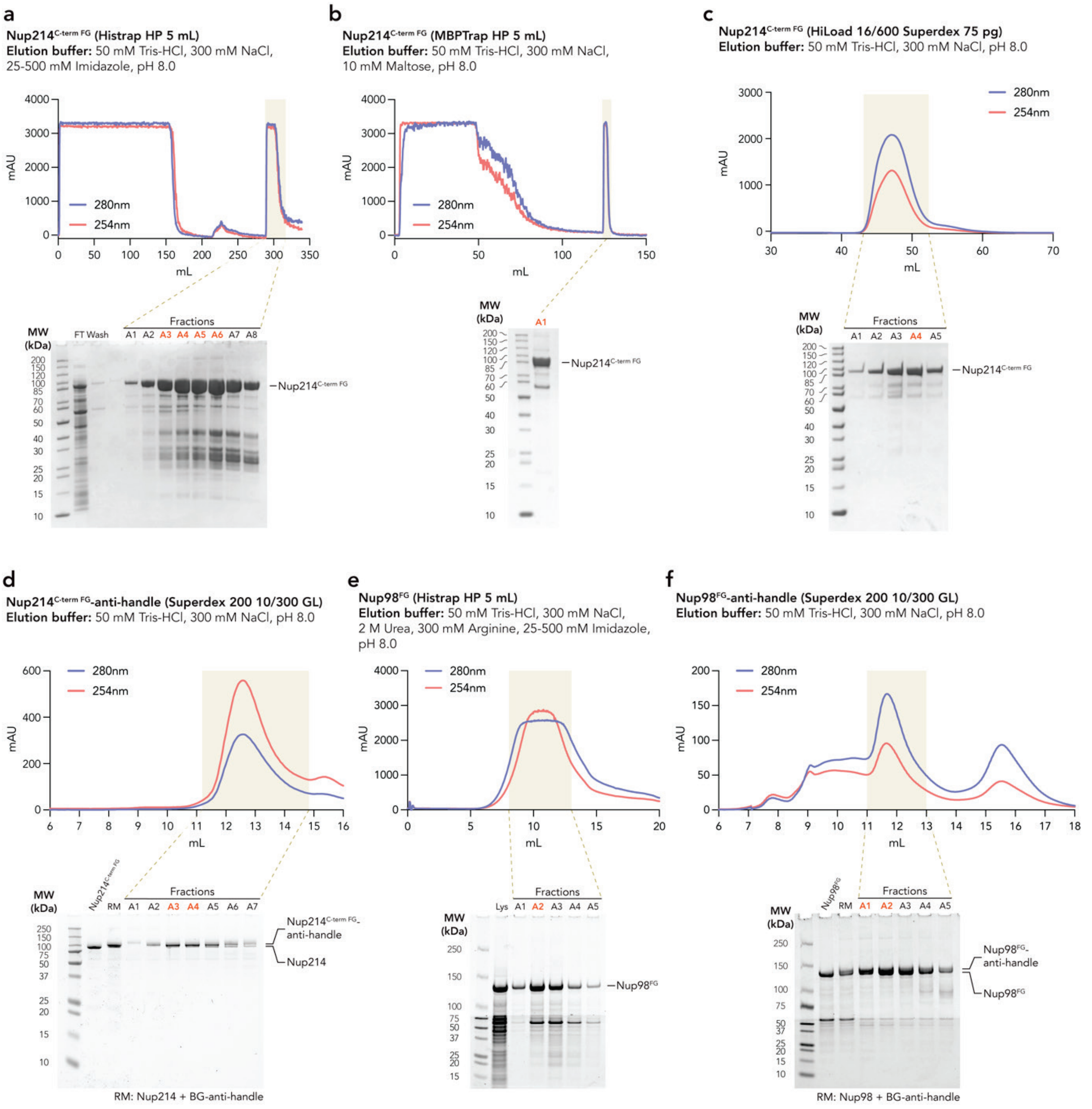


**Fig. S6 | Purification of Nup214$^{\text{C-term FG}}$, Nup98$^{\text{FG}}$, and their anti-handle conjugates. a–f**, Representative chromatograms and SDS-PAGE analyses of Nup214$^{\text{C-term FG}}$ purification by HisTrap affinity chromatography (**a**), MBPTrap affinity chromatography (**b**), and size-exclusion chromatography (**c**), purification of Nup214$^{\text{C-term FG}}$-anti-handle by size-exclusion chromatography (**d**), Nup98$^{\text{FG}}$ purification by HisTrap affinity chromatography (**e**), and purification of Nup98$^{\text{FG}}$-anti-handle by size-exclusion chromatography (**f**). Shaded regions indicate the fractions collected and analyzed by SDS-PAGE, and fractions highlighted in red were pooled for subsequent experiments. FT, flow-through; RM, reaction mixture; Lys, lysate.

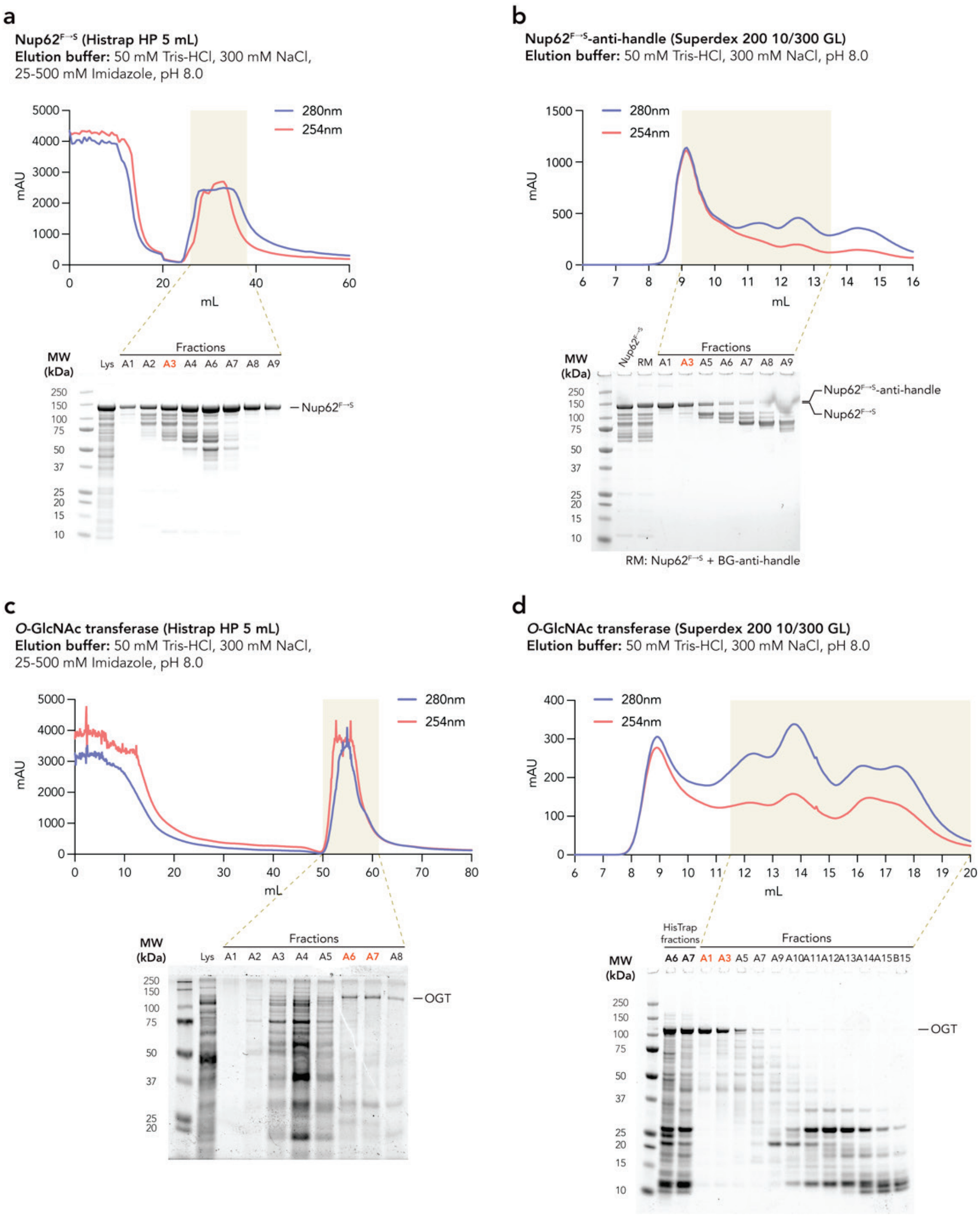


**Fig. S7 | Purification of Nup62$^{F\rightarrow S}$, its anti-handle conjugate, and *O*-GlcNAc transferase. a–d**, Representative chromatograms and SDS-PAGE analyses of Nup62$^{F\rightarrow S}$ purification by HisTrap affinity chromatography (**a**), purification of Nup62$^{F\rightarrow S}$-anti-handle by size-exclusion chromatography (**b**), *O*-GlcNAc transferase (OGT) purification by HisTrap affinity chromatography (**c**), and OGT purification by size-exclusion chromatography (**d**). Shaded regions indicate the fractions collected and analyzed by SDS-PAGE, and fractions highlighted in red were pooled for subsequent experiments. RM, reaction mixture; Lys, lysate.

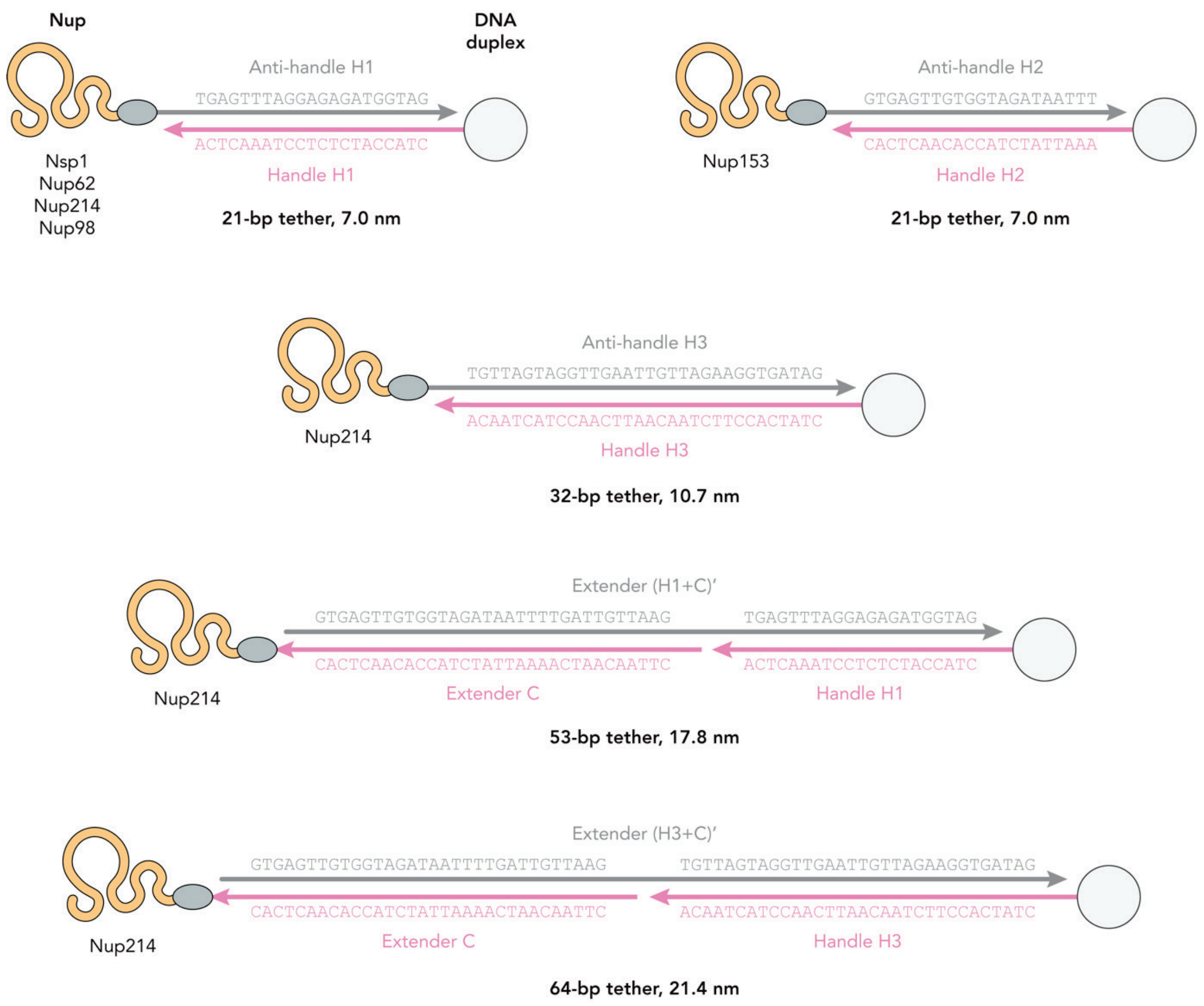


**Fig. S8 | Design of DNA tethers for anchoring Nups to the expandable ring.** Schematic illustration of the DNA tether designs used to anchor Nups to the expandable ring via hybridization. The light grey circle on the right denotes a DNA duplex on the inner-ring arc. Nsp1, Nup62, Nup98$^{FG}$, and Nup153$^{FG}$ were hybridized to the expandable ring using 21-bp DNA tethers (7.0 nm). Nup214 was hybridized using 21-, 32-, 53-, or 64-bp DNA tethers with expected lengths of 7.0, 10.7, 17.8, and 21.4 nm, respectively. Handle, anti-handle, and extender strand sequences are listed in **Table S3**. The prime symbol (') denotes oligonucleotides complementary to the corresponding handle and/or extender sequences.

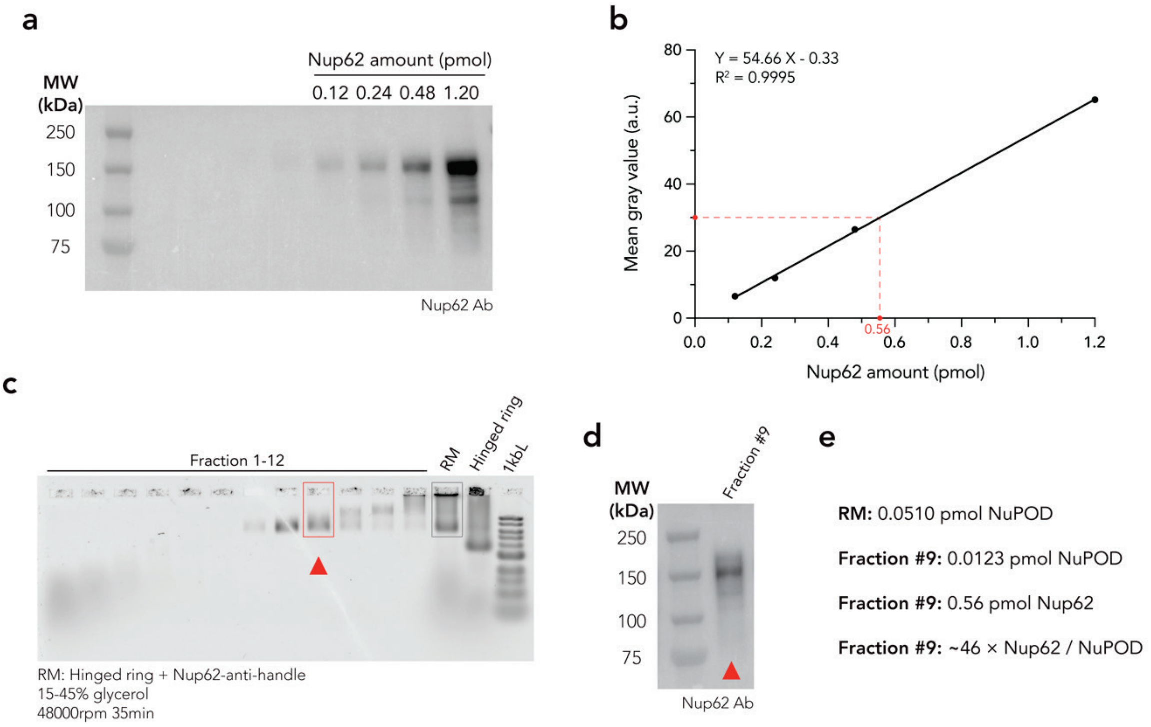


**Fig. S9 | Quantification of Nup62 copy number in Hinged Nup62-NuPOD. a**, Western blot of purified Nup62 standards used to generate a calibration curve. **b**, Calibration curve relating the mean gray value of the Nup62 band to the amount of Nup62. **c**, Agarose gel electrophoresis analysis of fractions collected after rate-zonal centrifugation (15–45% glycerol, 48,000 r.p.m., 35 min) of Hinged Nup62-NuPODs. The red arrowhead indicates fraction 9, which was selected for subsequent analysis. **d**, Western blot analysis of fraction 9 (on the same blot as **a**). **e**, Calculation of the average Nup62 copy number per NuPOD. RM, reaction mixture; 1 kbL, 1 kb DNA ladder.

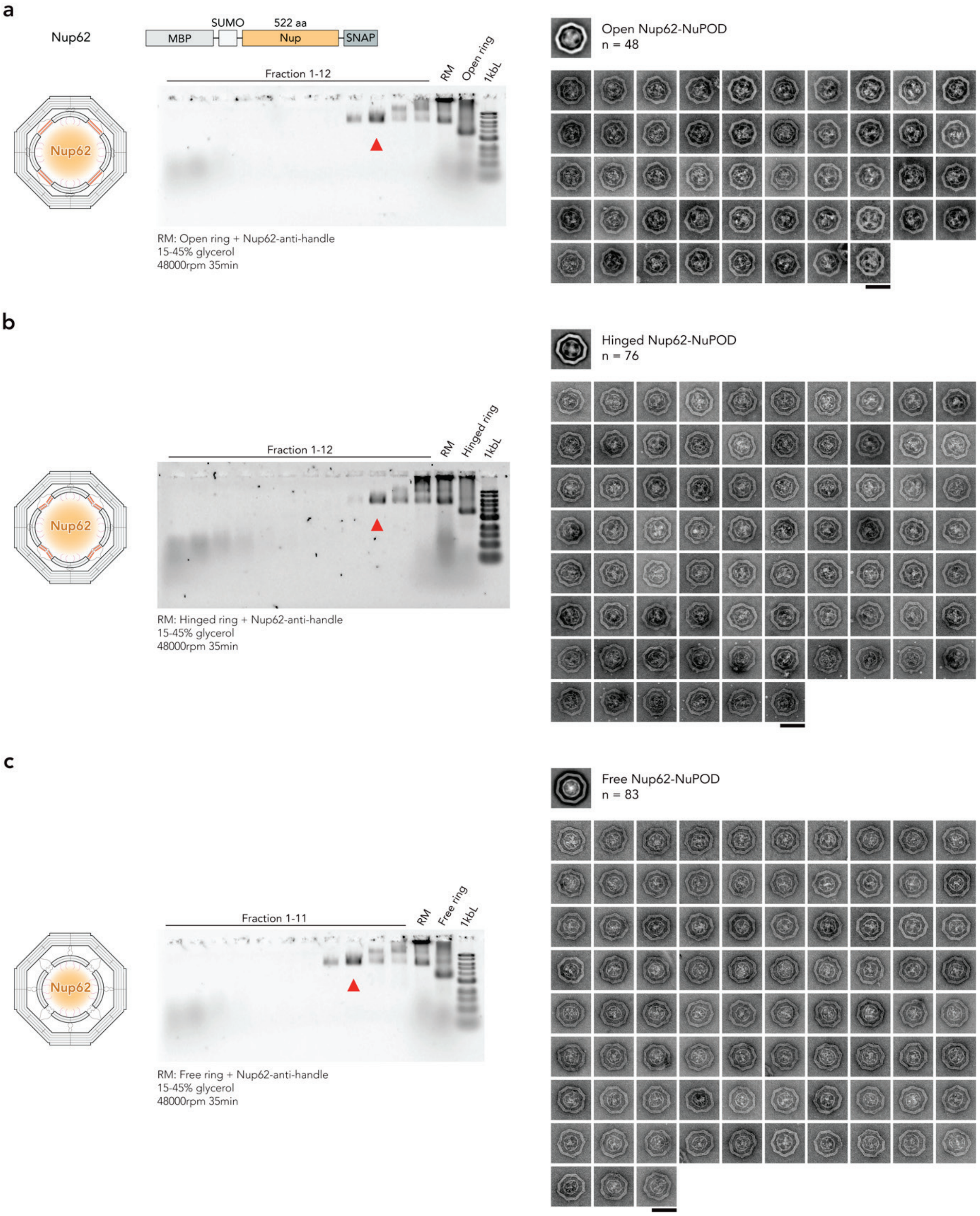


**Fig. S10 | Purification and structural characterization of Nup62-NuPODs. a–c**, Agarose gel electrophoresis analysis of fractions collected after rate-zonal centrifugation (15–45% glycerol, 48,000 r.p.m., 35 min) of Open (**a**), Hinged (**b**), and Free (**c**) Nup62-NuPODs. Red arrowheads indicate the fractions selected for negative-stain TEM analyses. Representative particle galleries of the purified Nup62-NuPODs are shown on the right. RM, reaction mixture; 1 kbL, 1 kb DNA ladder. Scale bars, 100 nm.

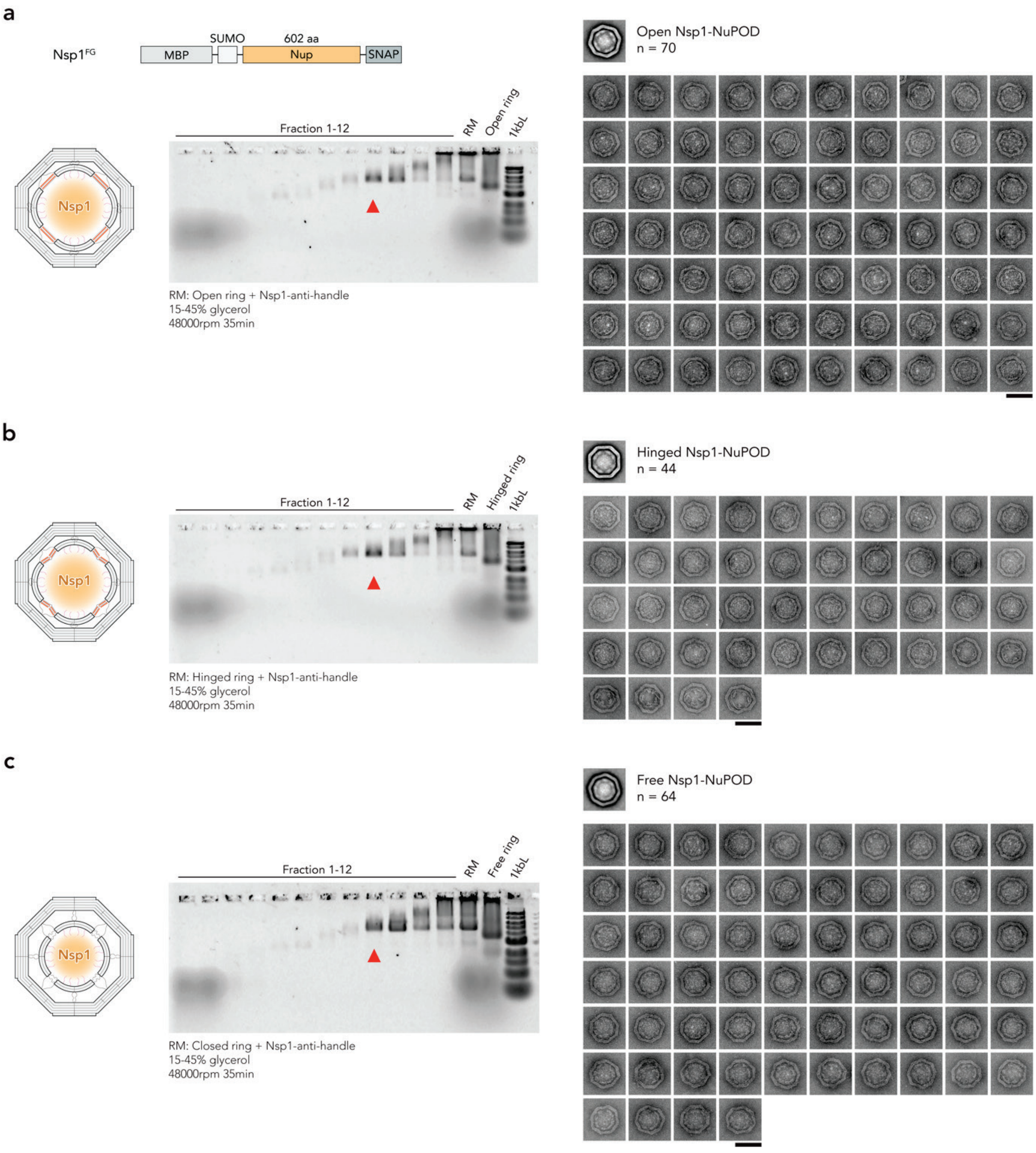


**Fig. S11 | Purification and structural characterization of Nsp1-NuPODs. a–c**, Agarose gel electrophoresis analysis of fractions collected after rate-zonal centrifugation (15–45% glycerol, 48,000 r.p.m., 35 min) of Open (**a**), Hinged (**b**), and Free (**c**) Nsp1-NuPODs. Red arrowheads indicate the fractions selected for negative-stain TEM analyses. Representative particle galleries of the purified Nsp1-NuPODs are shown on the right. RM, reaction mixture; 1 kbL, 1 kb DNA ladder. Scale bars, 100 nm.

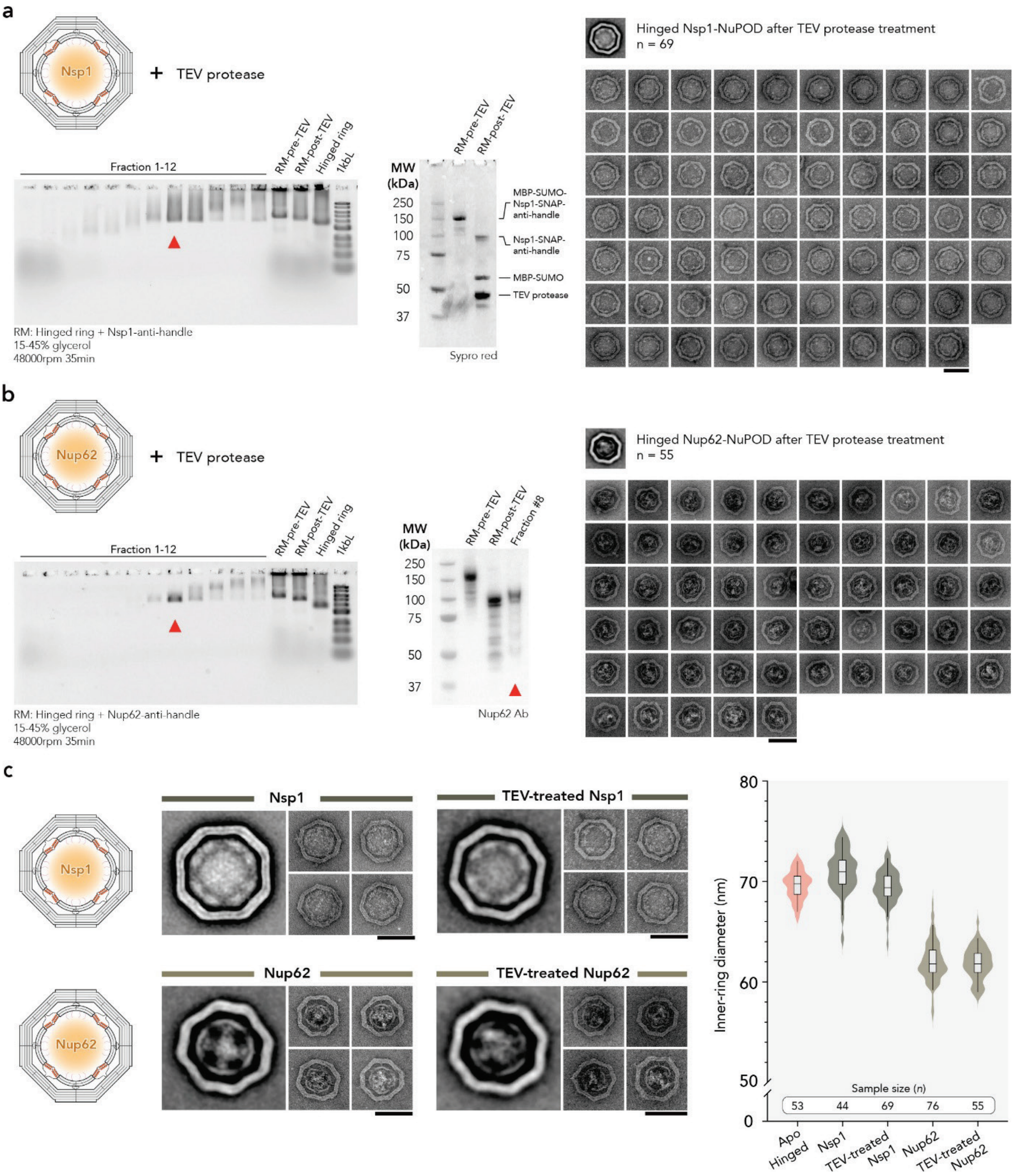


**Fig. S12 | TEV protease-treated Nsp1- and Nup62-NuPODs.** **a**, **b**, Purification and structural characterization of TEV-treated Hinged Nsp1- (**a**) and Nup62- (**b**) NuPODs. Agarose gel electrophoresis analysis of fractions collected after rate-zonal centrifugation (left), SDS-PAGE (**a**) or Western blot (**b**) confirming TEV cleavage (middle), and representative particle galleries of purified NuPODs (right, collected from fractions marked by red arrowheads). **c**, Schematics of Hinged NuPODs (left), negative-stain TEM analyses including a class average and four representative images for each NuPOD type (middle) and measured $d_{in}$ (right). The TEM images of untreated Nsp1- and Nup62-NuPODs are taken from **Fig. 2c**, whereas those of TEV-treated NuPODs are from panes **a** and **b** for comparison. Lines, boxes, and whiskers in violin plots show the median, 25th–75th, and 5th–95th percentiles, respectively. RM, reaction mixture; 1 kbL, 1 kb DNA ladder. Scale bars, 100 nm.

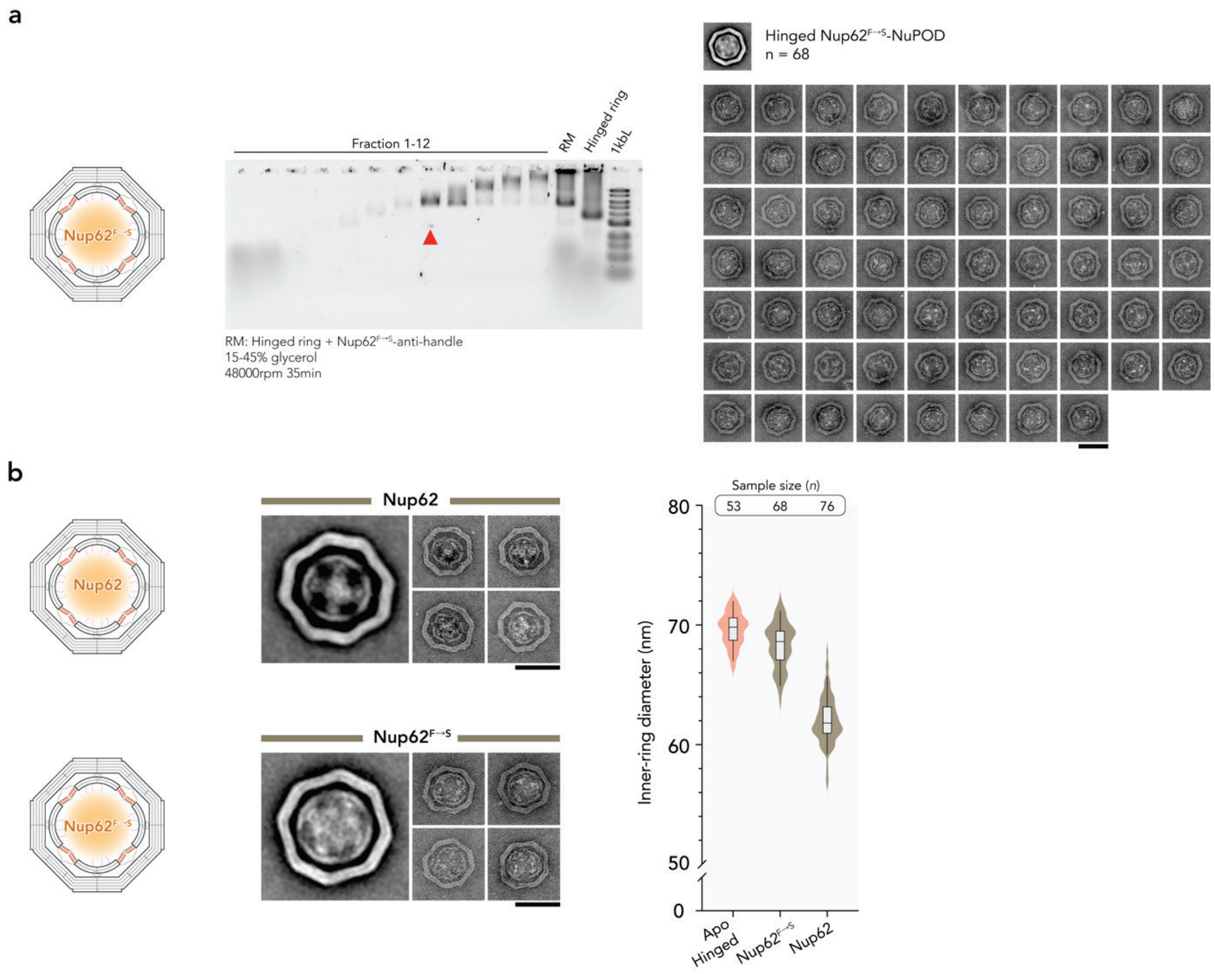


**Fig. S13 | Structural characterization of Hinged Nup62$^{F\rightarrow S}$-NuPOD. a**, Purification and structural characterization of Hinged Nup62$^{F\rightarrow S}$-NuPODs. Agarose gel electrophoresis analysis of fractions collected after rate-zonal centrifugation (15–45% glycerol, 48,000 r.p.m., 35 min). The red arrowhead indicates the fraction selected for negative-stain TEM analysis. Representative particle gallery of the purified Nup62$^{F\rightarrow S}$-NuPODs is shown on the right. **b**, Schematics of Hinged NuPODs (Left), negative-stain TEM analyses including a class average and four representative images for each NuPOD type (middle) and measured $d_{in}$ (right). The class average and particle galleries of Hinged Nup62-NuPODs are taken from **Fig. 2c**, whereas those of Hinged Nup62$^{F\rightarrow S}$-NuPODs are taken from panel **a** for comparison. Lines, boxes, and whiskers in violin plots show the median, 25th–75th, and 5th–95th percentiles, respectively. RM, reaction mixture; 1 kbL, 1 kb DNA ladder. Scale bars, 100 nm.

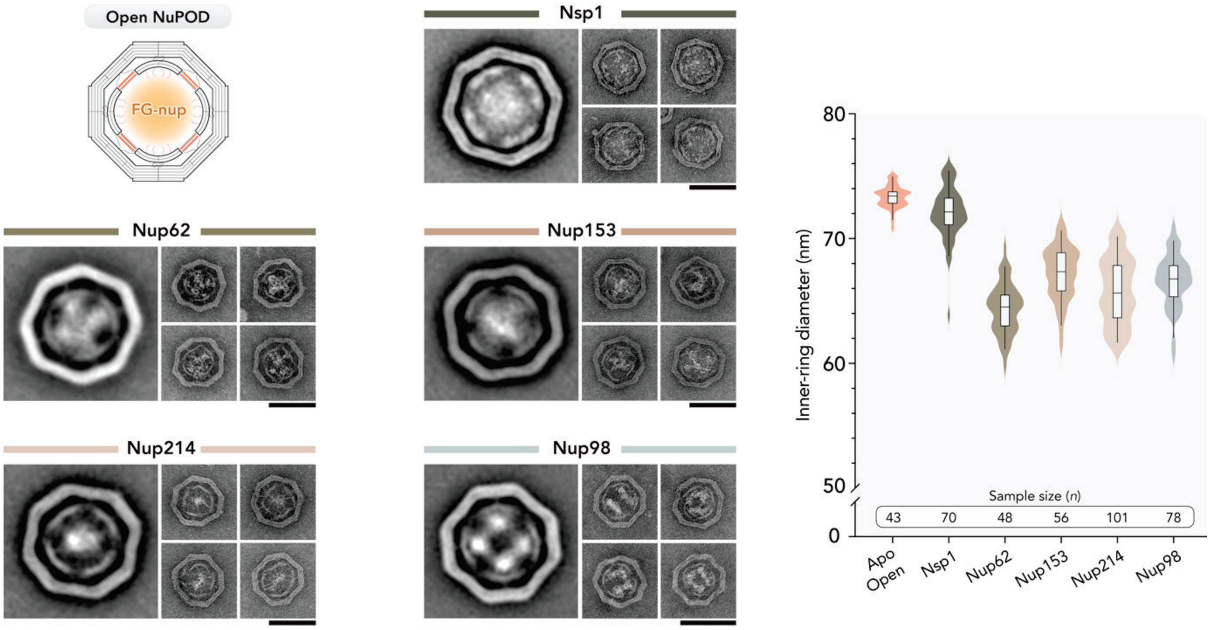


**Fig. S14 | Structural characterization of Open NuPODs.** Class averages, representative particle galleries, and $d_{in}$ measurement of Open Nsp1-, Nup62-, Nup153-, Nup214-, and Nup98-NuPODs. Nup-free (Apo) ring size is taken from **Fig. 1** for comparison. Lines, boxes, and whiskers in violin plots show the median, 25th–75th, and 5th–95th percentiles, respectively. Scale bars, 100 nm.

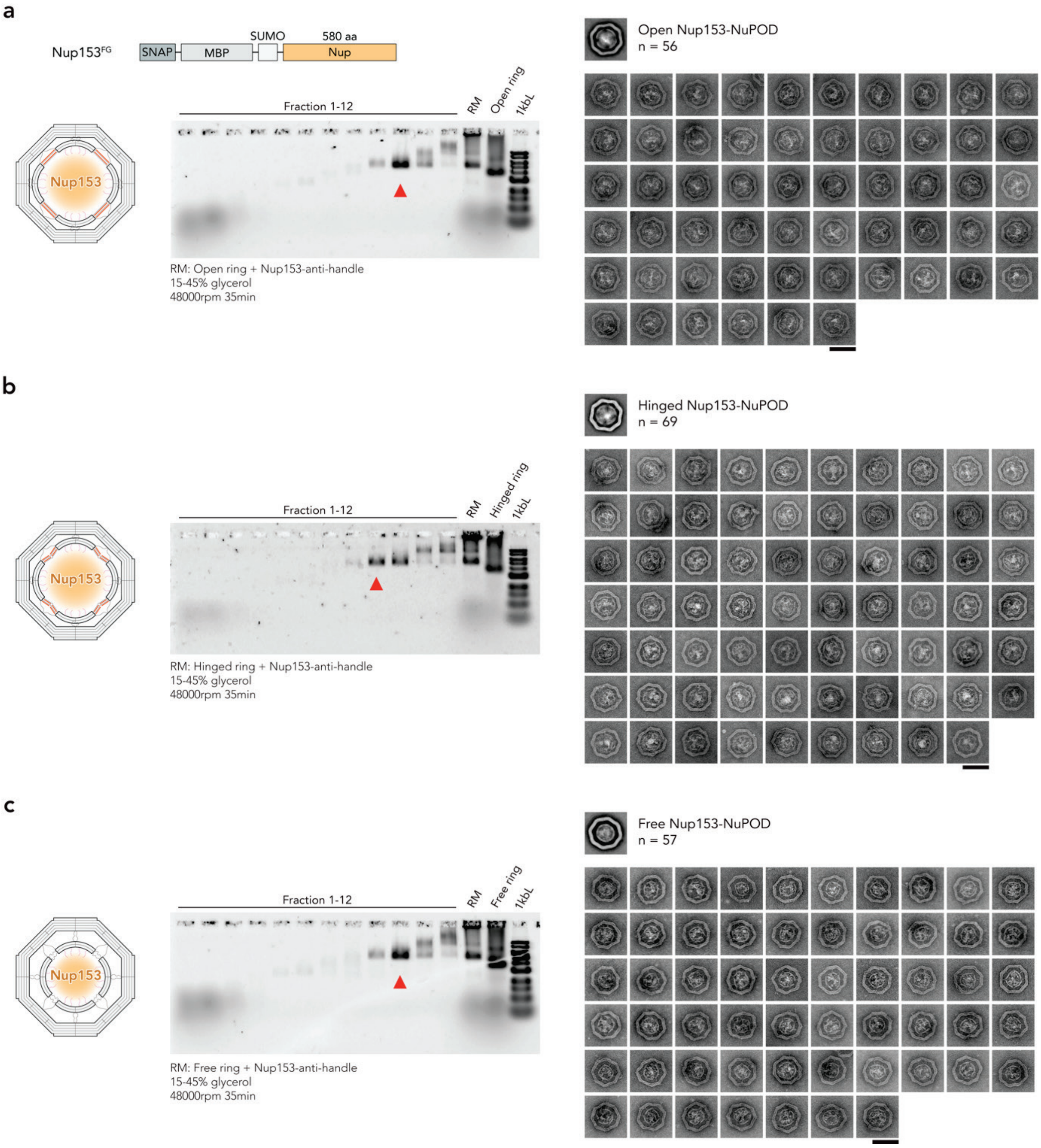


**Fig. S15 | Purification and structural characterization of Nup153-NuPODs. a–c**, Agarose gel electrophoresis analysis of fractions collected after rate-zonal centrifugation (15–45% glycerol, 48,000 r.p.m., 35 min) of Open (**a**), Hinged (**b**), and Free (**c**) Nup153-NuPODs. Red arrowheads indicate the fractions selected for negative-stain TEM analyses. Representative particle galleries of the purified Nup153-NuPODs are shown on the right. RM, reaction mixture; 1 kbL, 1 kb DNA ladder. Scale bars, 100 nm.

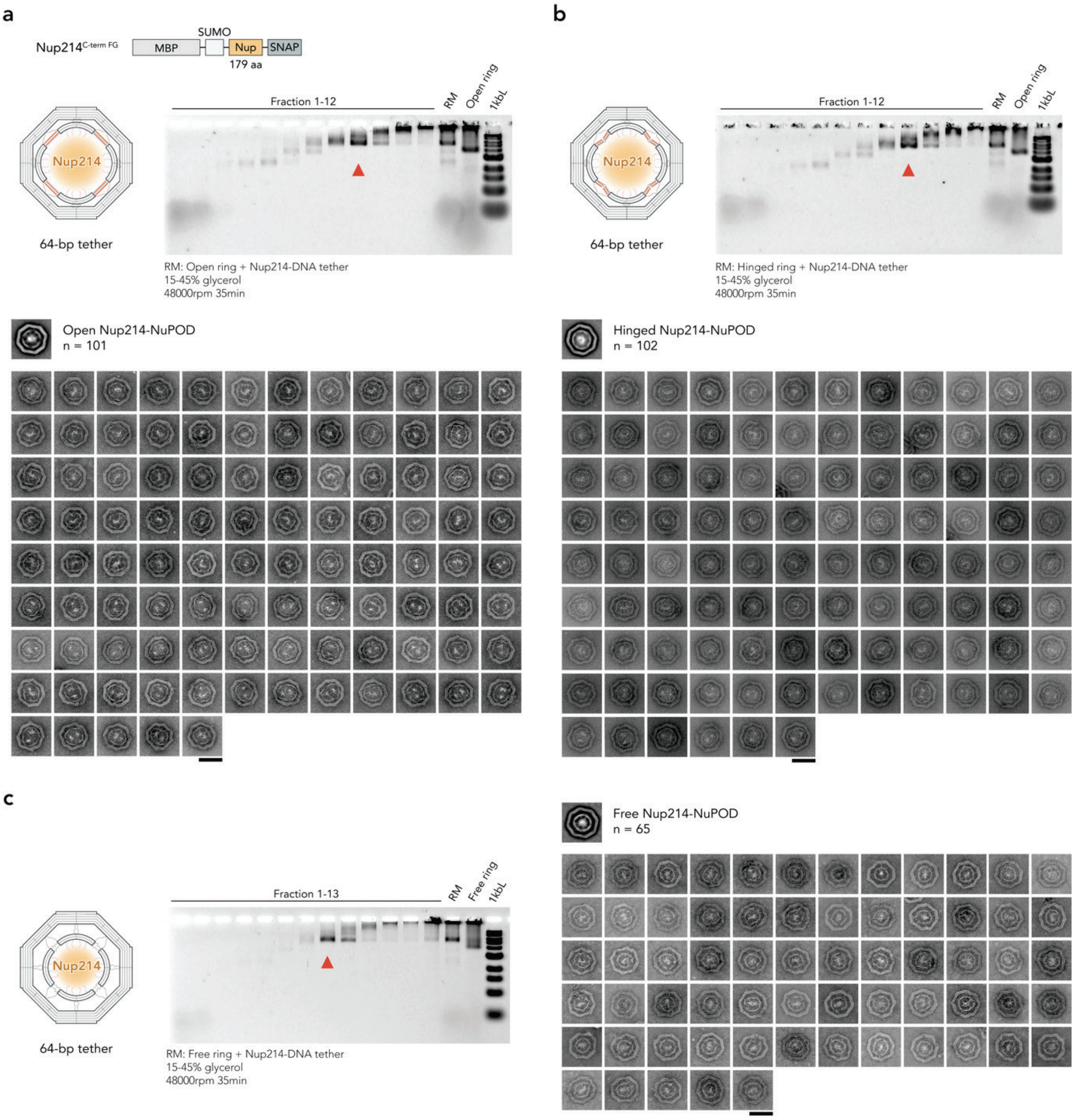


**Fig. S16 | Purification and structural characterization of Nup214-NuPODs assembled with 64-bp DNA tethers. a–c**, Agarose gel electrophoresis analysis of fractions collected after rate-zonal centrifugation (15–45% glycerol, 48,000 r.p.m., 35 min) of Open (**a**), Hinged (**b**), and Free (**c**) Nup214-NuPODs assembled with 64-bp DNA tethers. Red arrowheads indicate the fractions selected for negative-stain TEM analyses. Representative particle galleries of the purified Nup214-NuPODs are shown. RM, reaction mixture; 1 kbL, 1 kb DNA ladder. Scale bars, 100 nm.

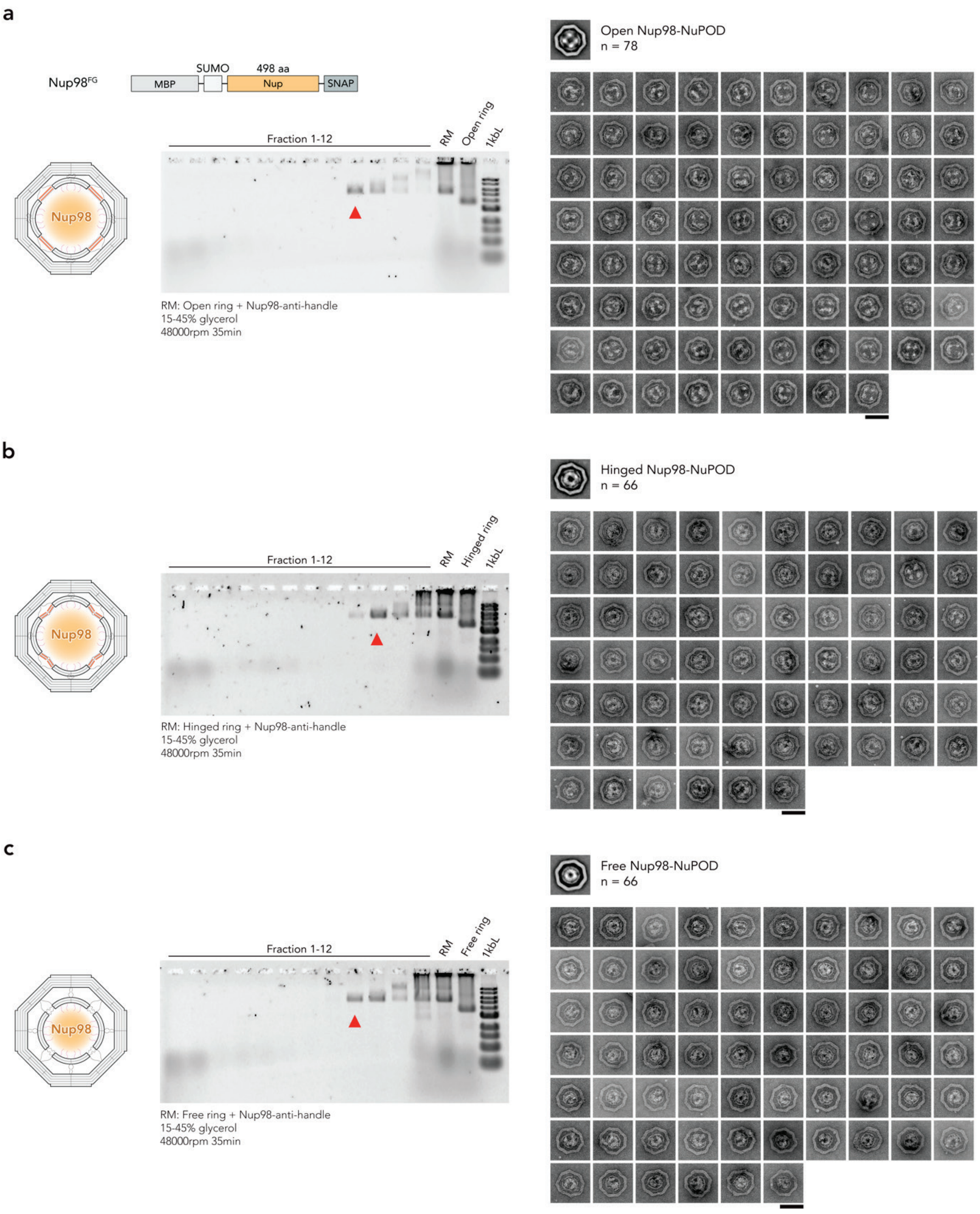


**Fig. S17 | Purification and structural characterization of Nup98-NuPODs. a–c**, Agarose gel electrophoresis analysis of fractions collected after rate-zonal centrifugation (15–45% glycerol, 48,000 r.p.m., 35 min) of Open (**a**), Hinged (**b**), and Free (**c**) Nup98-NuPODs. Red arrowheads indicate the fractions selected for negative-stain TEM analyses. Representative particle galleries of the purified Nup98-NuPODs are shown on the right. RM, reaction mixture; 1 kbL, 1 kb DNA ladder. Scale bars, 100 nm.

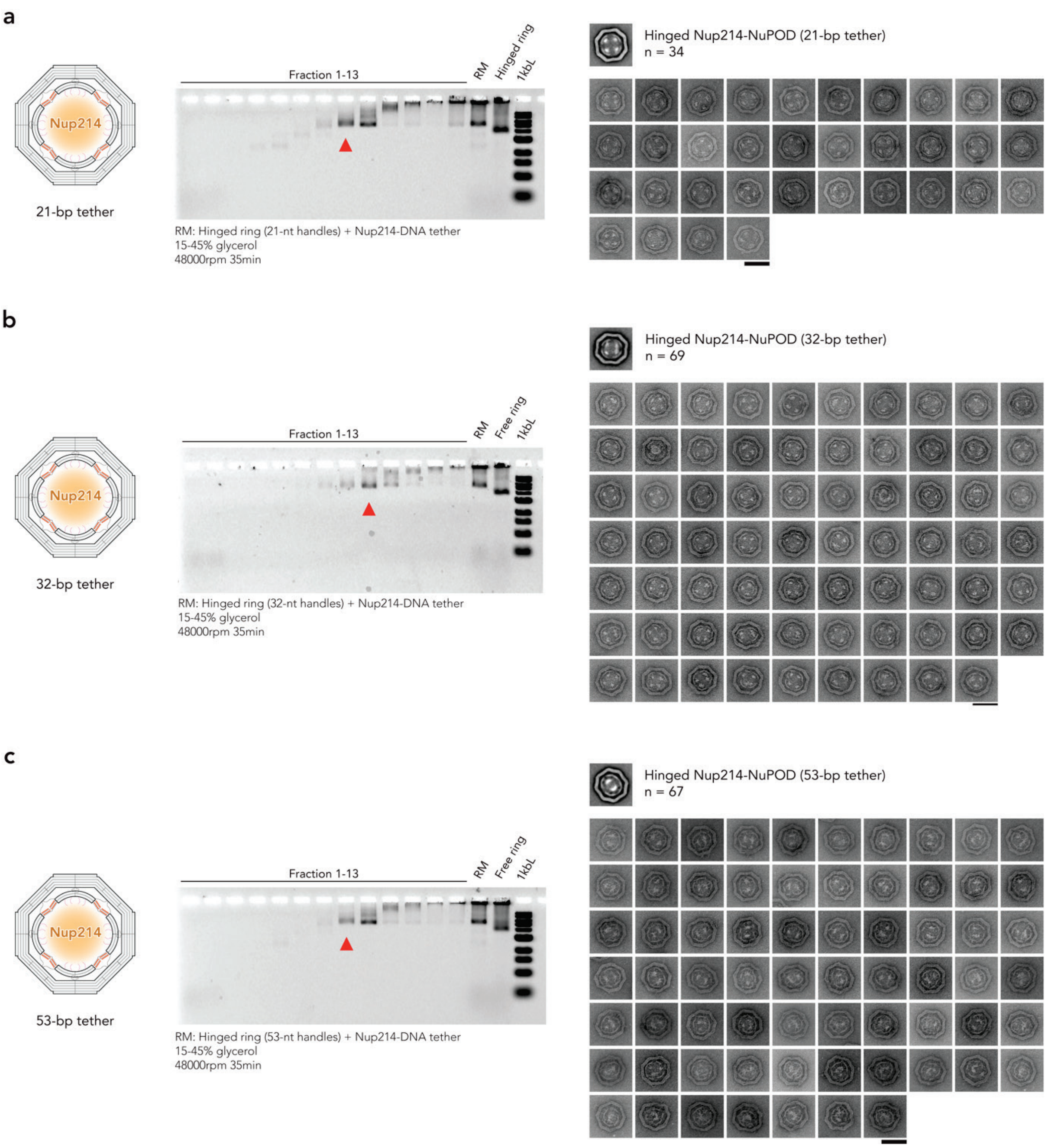


**Fig. S18 | Purification and structural characterization of Hinged Nup214-NuPODs assembled with 21-, 32-, and 53-bp DNA tethers. a–c**, Agarose gel electrophoresis analysis of fractions collected after rate-zonal centrifugation (15–45% glycerol, 48,000 r.p.m., 35 min) of Hinged Nup214-NuPODs assembled with 21-bp (**a**), 32-bp (**b**), and 53-bp (**c**) DNA tethers. Red arrowheads indicate the fractions selected for negative-stain TEM analyses. Representative particle galleries of the purified Nup214-NuPODs are shown on the right. RM, reaction mixture; 1 kbL, 1 kb DNA ladder. Scale bars, 100 nm.

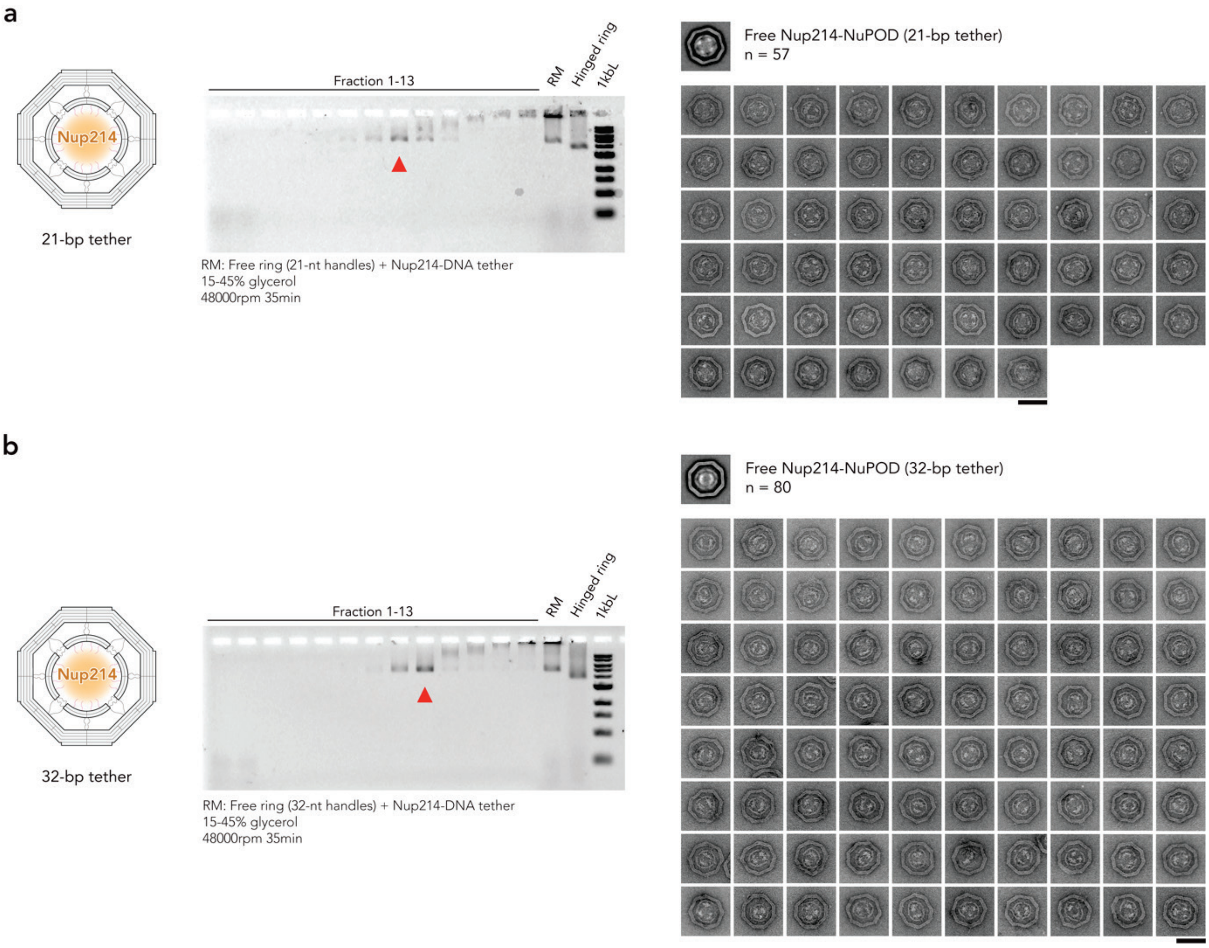


**Fig. S19 | Purification and structural characterization of Free Nup214-NuPODs assembled with 21- and 32-bp DNA tethers. a**, **b**, Agarose gel electrophoresis analysis of fractions collected after rate-zonal centrifugation (15–45% glycerol, 48,000 r.p.m., 35 min) of Free Nup214-NuPODs assembled with 21-bp (**a**) and 32-bp (**b**) DNA tethers. Red arrowheads indicate the fractions selected for negative-stain TEM analyses. Representative particle galleries of the purified Nup214-NuPODs are shown on the right. RM, reaction mixture; 1 kbL, 1 kb DNA ladder. Scale bars, 100 nm.

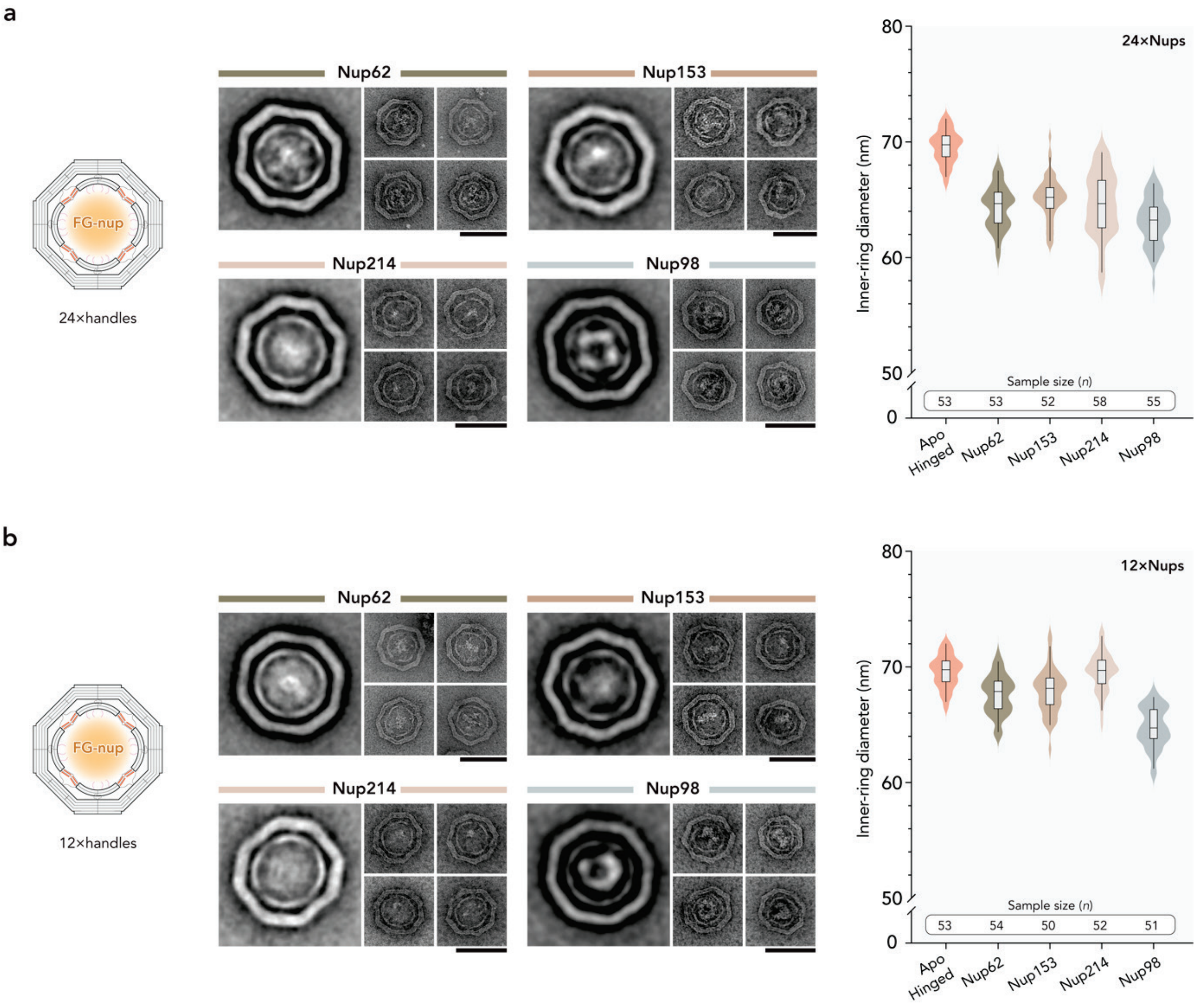


**Fig. S20 | Structural characterization of Hinged NuPODs assembled with reduced Nup copies. a**, **b**, Class averages, representative particle galleries, and $d_{in}$ measurement of Hinged NuPODs assembled with 24 (**a**) or 12 (**b**) Nup copies per ring. Nup-free (Apo) ring size is taken from **Fig. 1** for comparison. Lines, boxes, and whiskers in violin plots show the median, 25th–75th, and 5th–95th percentiles, respectively. Scale bars, 100 nm.

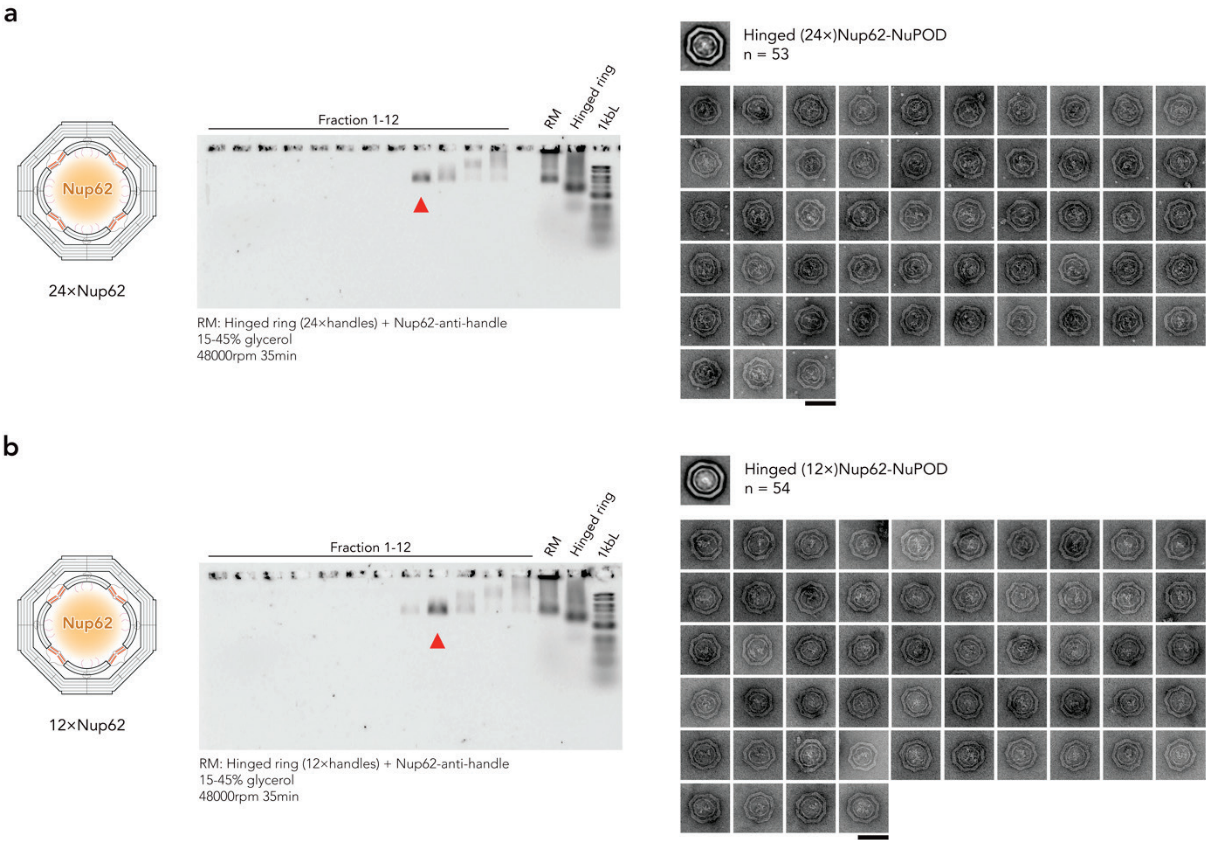


**Fig. S21 | Purification and structural characterization of Hinged NuPODs assembled with reduced Nup62 copies. a**, **b**, Agarose gel electrophoresis analysis of fractions collected after rate-zonal centrifugation (15–45% glycerol, 48,000 r.p.m., 35 min) of Hinged NuPODs assembled with 24 (**a**) or 12 (**b**) Nup62. Red arrowheads indicate the fractions selected for negative-stain TEM analyses. Representative particle galleries of the purified Nup62-NuPODs are shown on the right. RM, reaction mixture; 1 kbL, 1 kb DNA ladder. Scale bars, 100 nm.

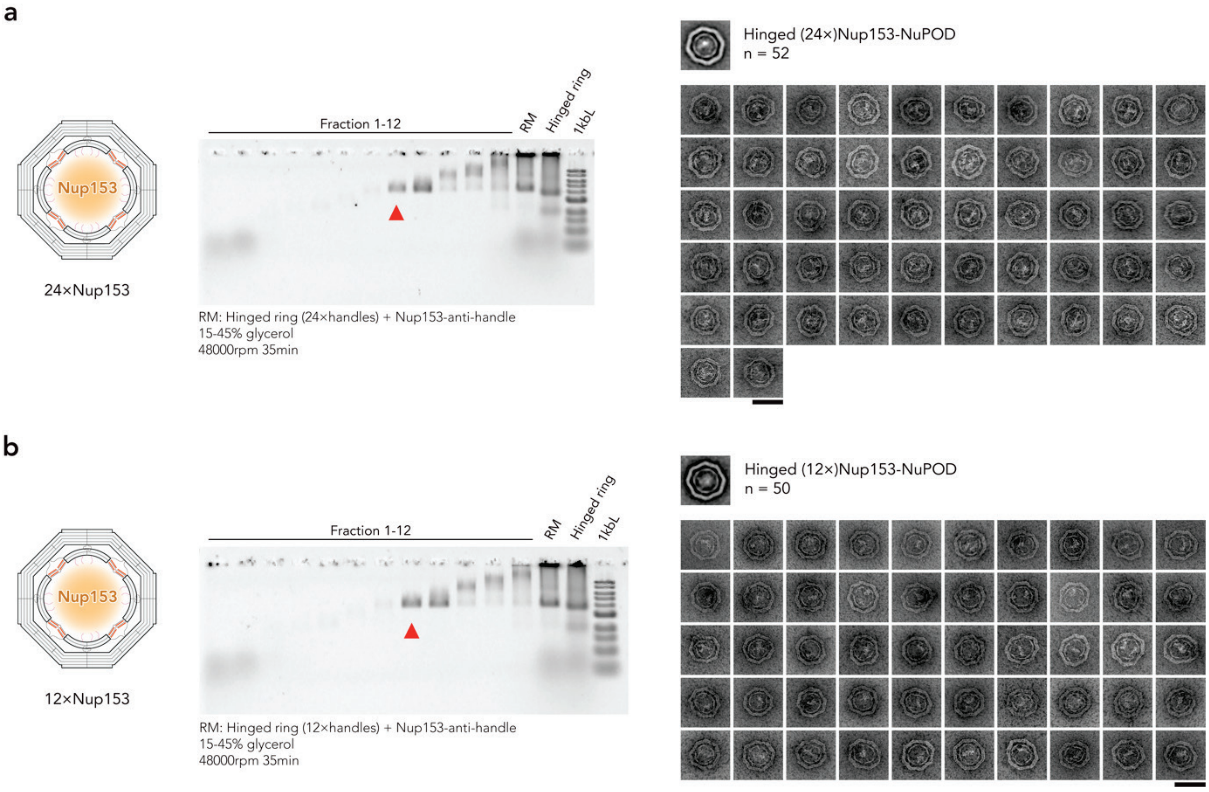


**Fig. S22 | Purification and structural characterization of Hinged NuPODs assembled with reduced Nup153 copies. a**, **b**, Agarose gel electrophoresis analysis of fractions collected after rate-zonal centrifugation (15–45% glycerol, 48,000 r.p.m., 35 min) of Hinged NuPODs assembled with 24 (**a**) or 12 (**b**) Nup153. Red arrowheads indicate the fractions selected for negative-stain TEM analyses. Representative particle galleries of the purified Nup153-NuPODs are shown on the right. RM, reaction mixture; 1 kbL, 1 kb DNA ladder. Scale bars, 100 nm.

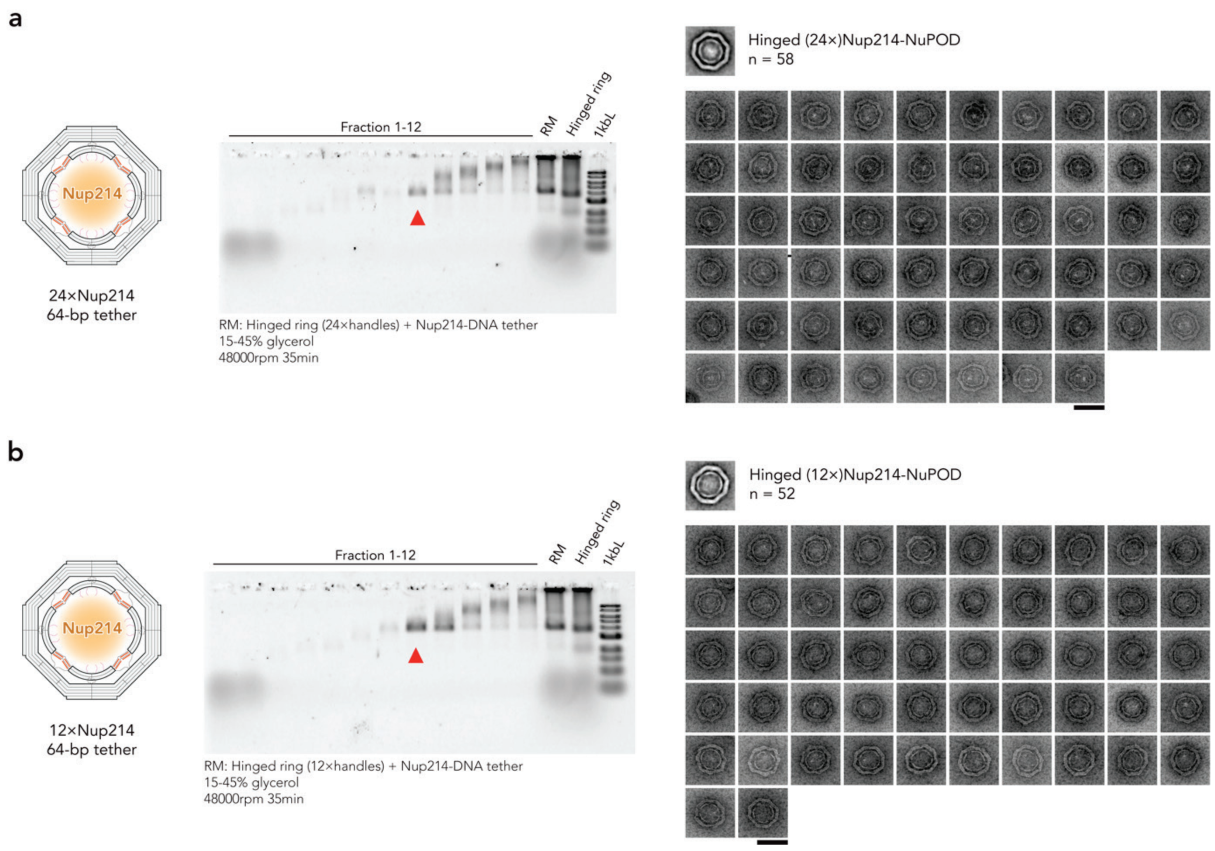


**Fig. S23 | Purification and structural characterization of Hinged NuPODs assembled with reduced Nup214 copies using 64-bp DNA tethers. a**, **b**, Agarose gel electrophoresis analysis of fractions collected after rate-zonal centrifugation (15–45% glycerol, 48,000 r.p.m., 35 min) of Hinged NuPODs assembled with 24 (**a**) or 12 (**b**) Nup214 using 64-bp DNA tethers. Red arrowheads indicate the fractions selected for negative-stain TEM analyses. Representative particle galleries of the purified Nup214-NuPODs are shown on the right. RM, reaction mixture; 1 kbL, 1 kb DNA ladder. Scale bars, 100 nm.

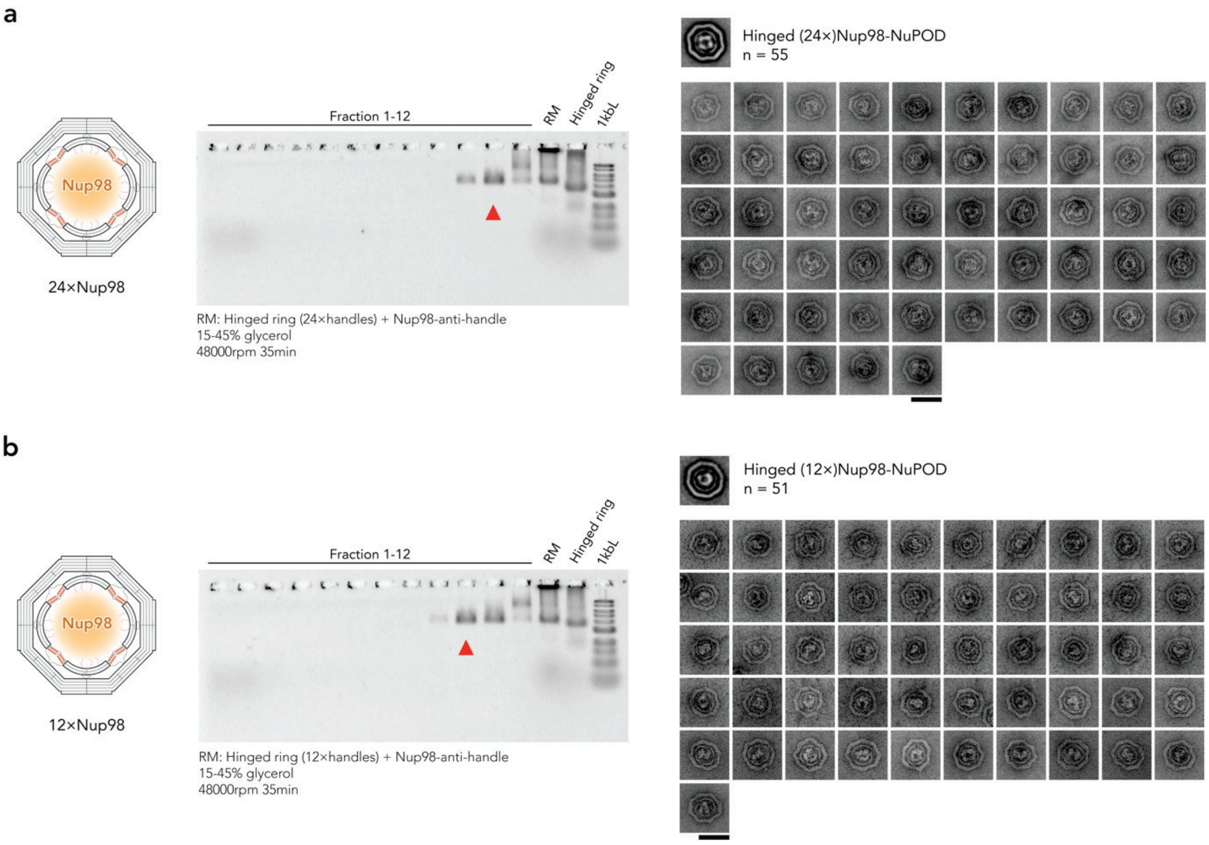


**Fig. S24 | Purification and structural characterization of Hinged NuPODs assembled with reduced Nup98 copies. a**, **b**, Agarose gel electrophoresis analysis of fractions collected after rate-zonal centrifugation (15–45% glycerol, 48,000 r.p.m., 35 min) of Hinged NuPODs assembled with 24 (**a**) or 12 (**b**) Nup98. Red arrowheads indicate the fractions selected for negative-stain TEM analyses. Representative particle galleries of the purified Nup98-NuPODs are shown on the right. RM, reaction mixture; 1 kbL, 1 kb DNA ladder. Scale bars, 100 nm.

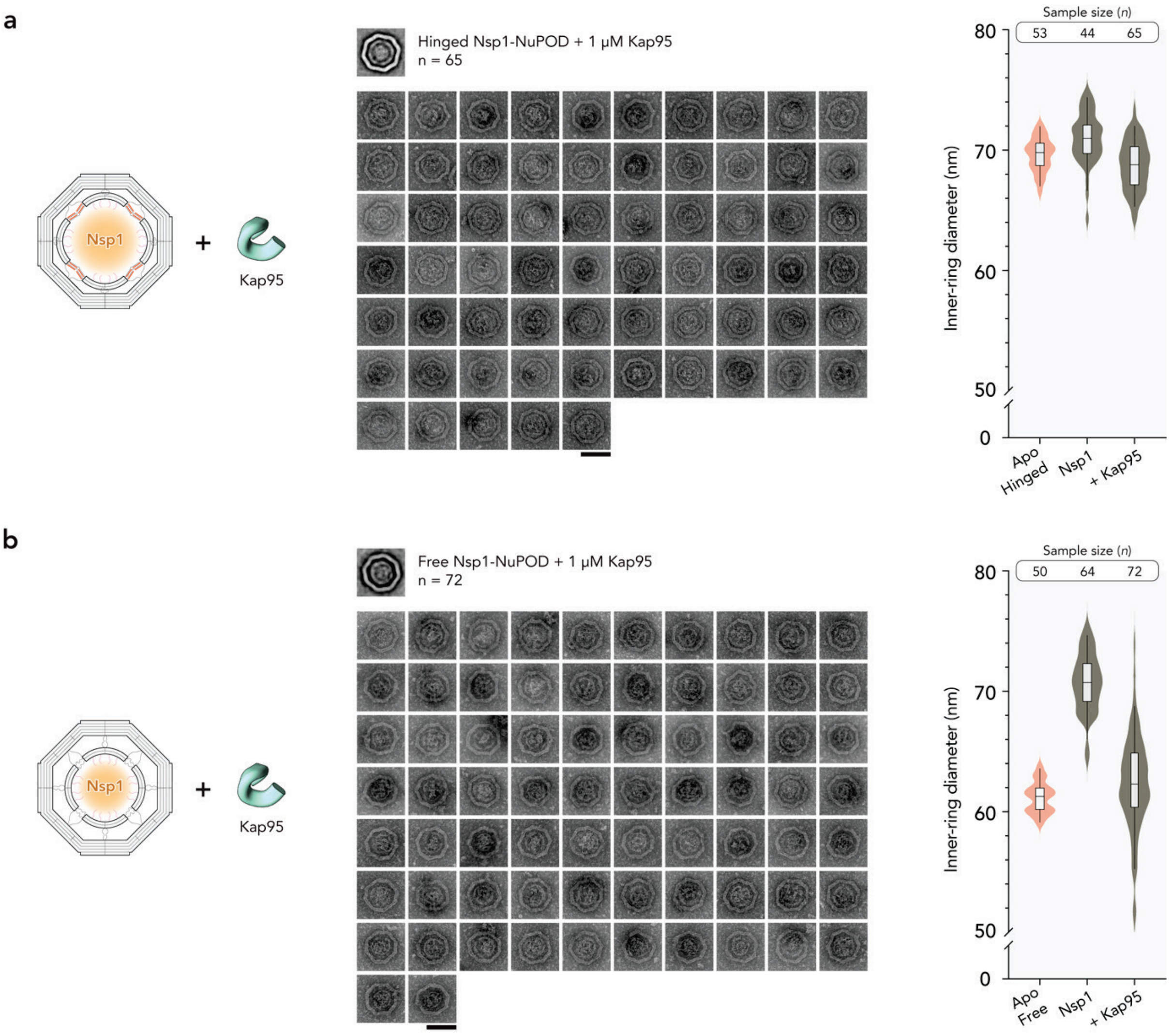


**Fig. S25 | Structural characterization of Nsp1-NuPODs incubated with Kap95. a**, **b**, Representative particle galleries, and $d_{\mathrm{in}}$ measurement of Hinged (**a**) and Free (**b**) Nsp1-NuPODs after incubation with 1 μM Kap95. Nup-free (Apo) ring sizes are taken from **Fig. 1** for comparison. Lines, boxes, and whiskers in violin plots show the median, 25th–75th, and 5th–95th percentiles, respectively. Scale bars, 100 nm.

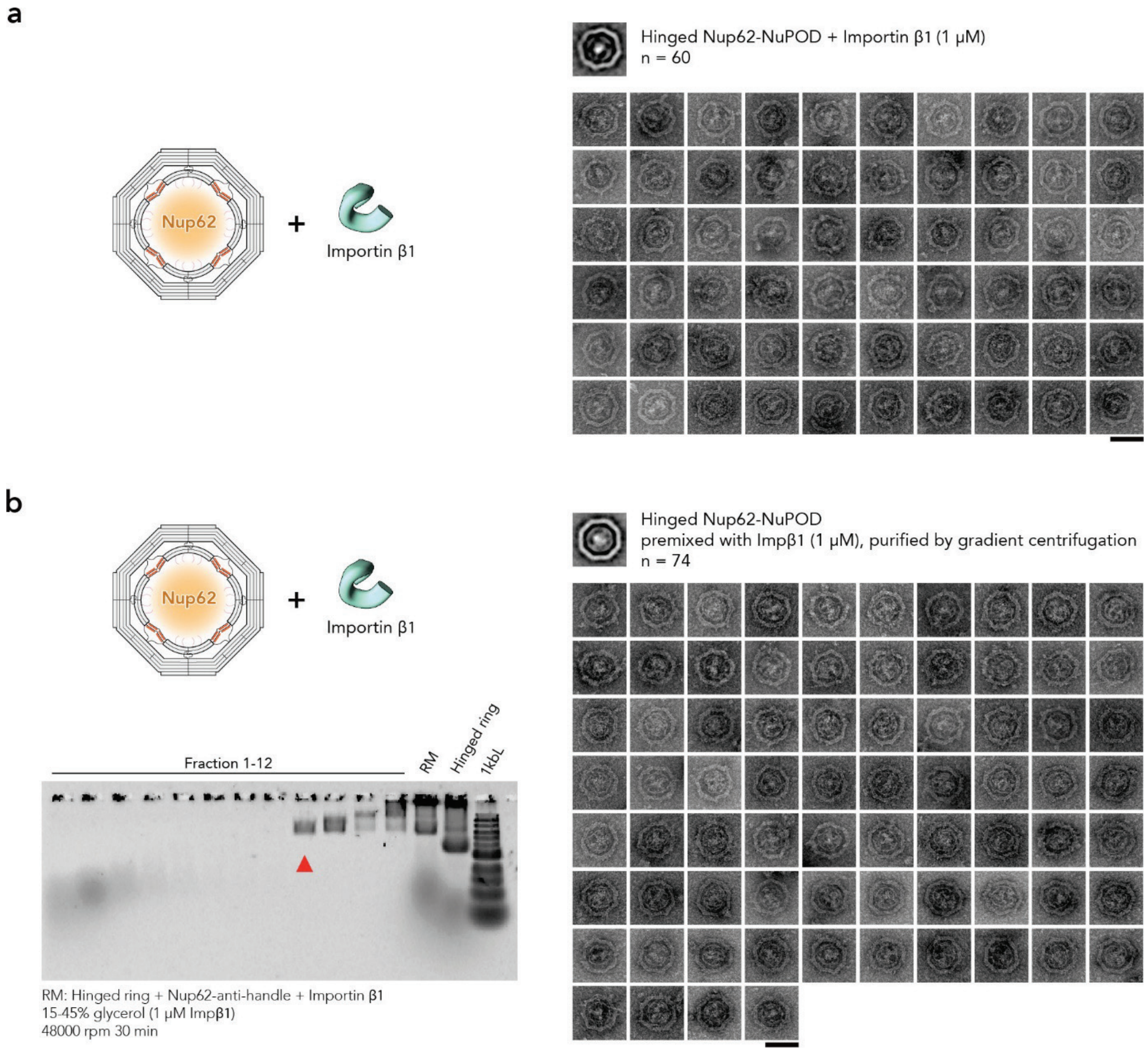


**Fig. S26 | Structural characterization of Nup62-NuPODs incubated with importin β1. a**, Representative particle gallery of Hinged Nup62-NuPODs after incubation with 1 μM importin β1 (Impβ1). **b**, Agarose gel electrophoresis analysis of fractions collected after rate-zonal centrifugation (15–45% glycerol, 1 μM Impβ1, 48,000 r.p.m., 30 min) of premixed Hinged Nup62-NuPODs and 1 μM Impβ1 (see **Methods**). Red arrowhead indicates the fraction selected for negative-stain TEM analysis. Representative particle gallery of the recovered Nup62-NuPODs is shown on the right. RM, reaction mixture; 1 kbL, 1 kb DNA ladder. Scale bars, 100 nm.

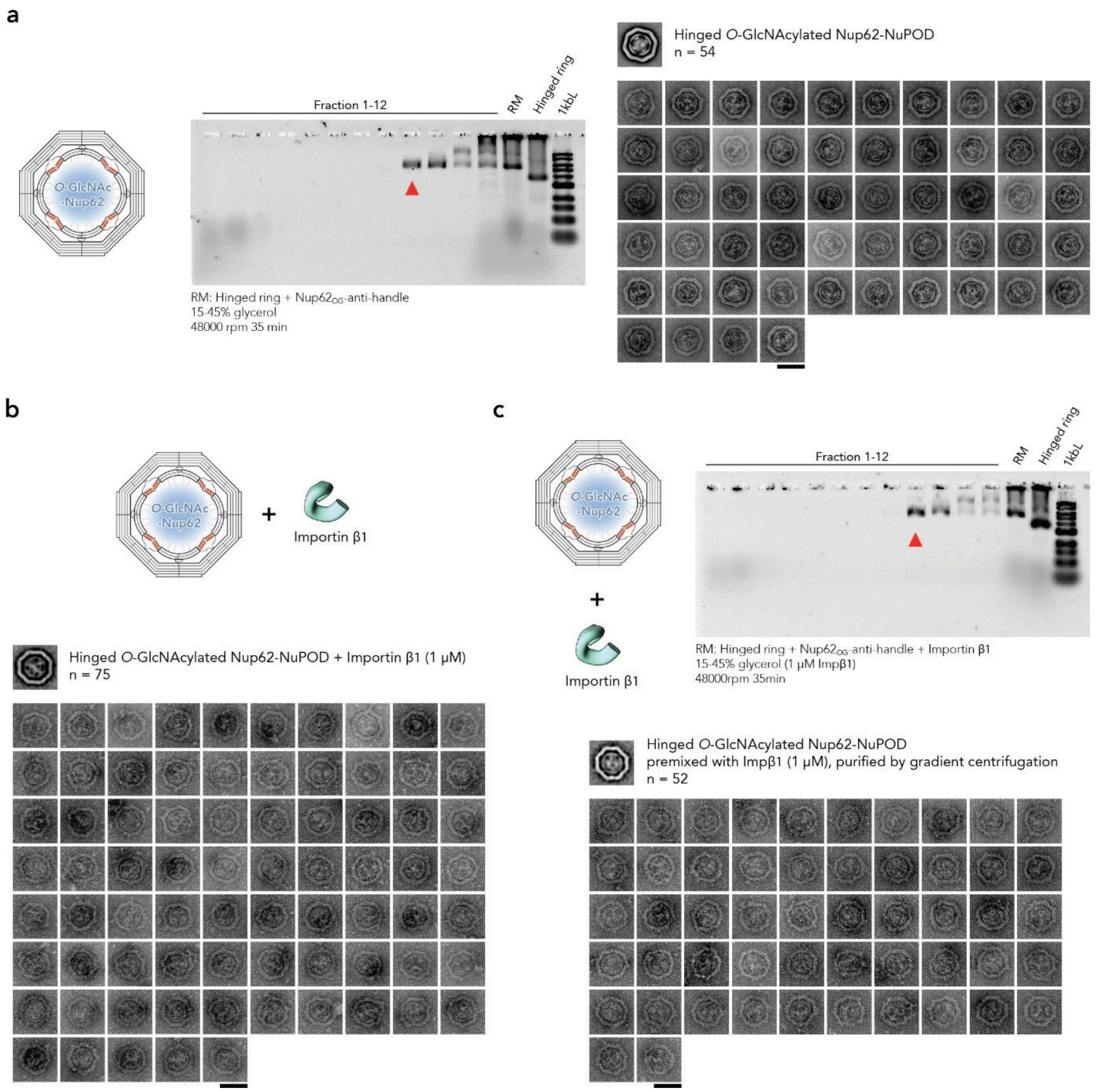


**Fig. S27 | Structural characterization of Hinged *O*-GlcNAcylated Nup62-NuPODs with and without importin β1. a**, Agarose gel electrophoresis analysis of fractions collected after rate-zonal centrifugation (15–45% glycerol, 48,000 r.p.m., 35 min) of Hinged *O*-GlcNAcylated Nup62 ($Nup62_{OG}$)-NuPODs. The red arrowhead indicates the fraction selected for negative-stain TEM analysis. Representative averaged particle and particle gallery of the purified $Nup62_{OG}$-NuPODs are shown below. **b**, Representative particle gallery of Hinged $Nup62_{OG}$-NuPODs after incubation with 1 µM importin β1 (Impβ1). **c**, Agarose gel electrophoresis analysis of fractions collected after rate-zonal centrifugation (15–45% glycerol, 1 µM Impβ1, 48,000 r.p.m., 30 min) of premixed Hinged $Nup62_{OG}$-NuPODs and 1 µM Impβ1 (see **Methods**). Red arrowhead indicates the fraction selected for negative-stain TEM analysis. Representative particle gallery of the recovered $Nup62_{OG}$-NuPODs is shown below. RM, reaction mixture; 1 kbL, 1 kb DNA ladder. Scale bars, 100 nm.

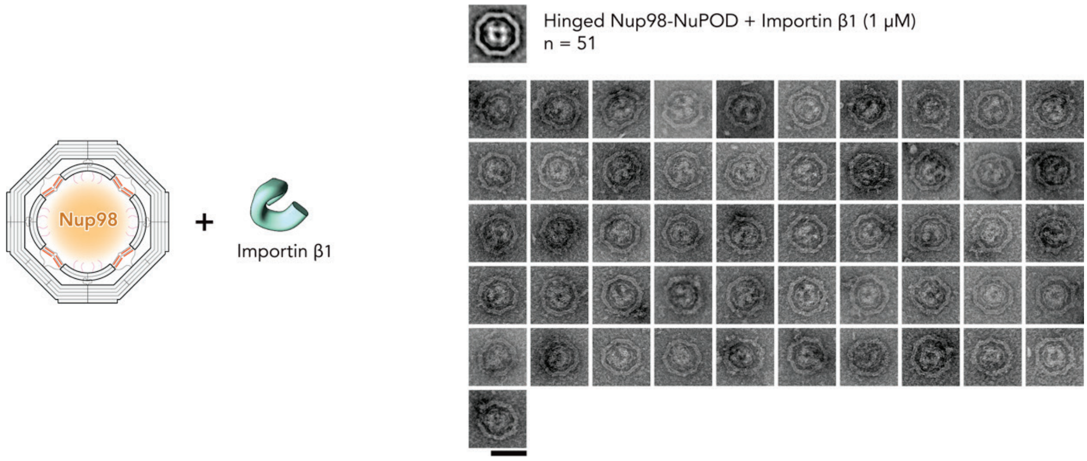


**Fig. S28 | Structural characterization of Hinged Nup98-NuPODs incubated with importin β1.** Representative particle gallery of Hinged Nup98-NuPODs after incubation with 1 μM importin β1. Scale bars, 100 nm.

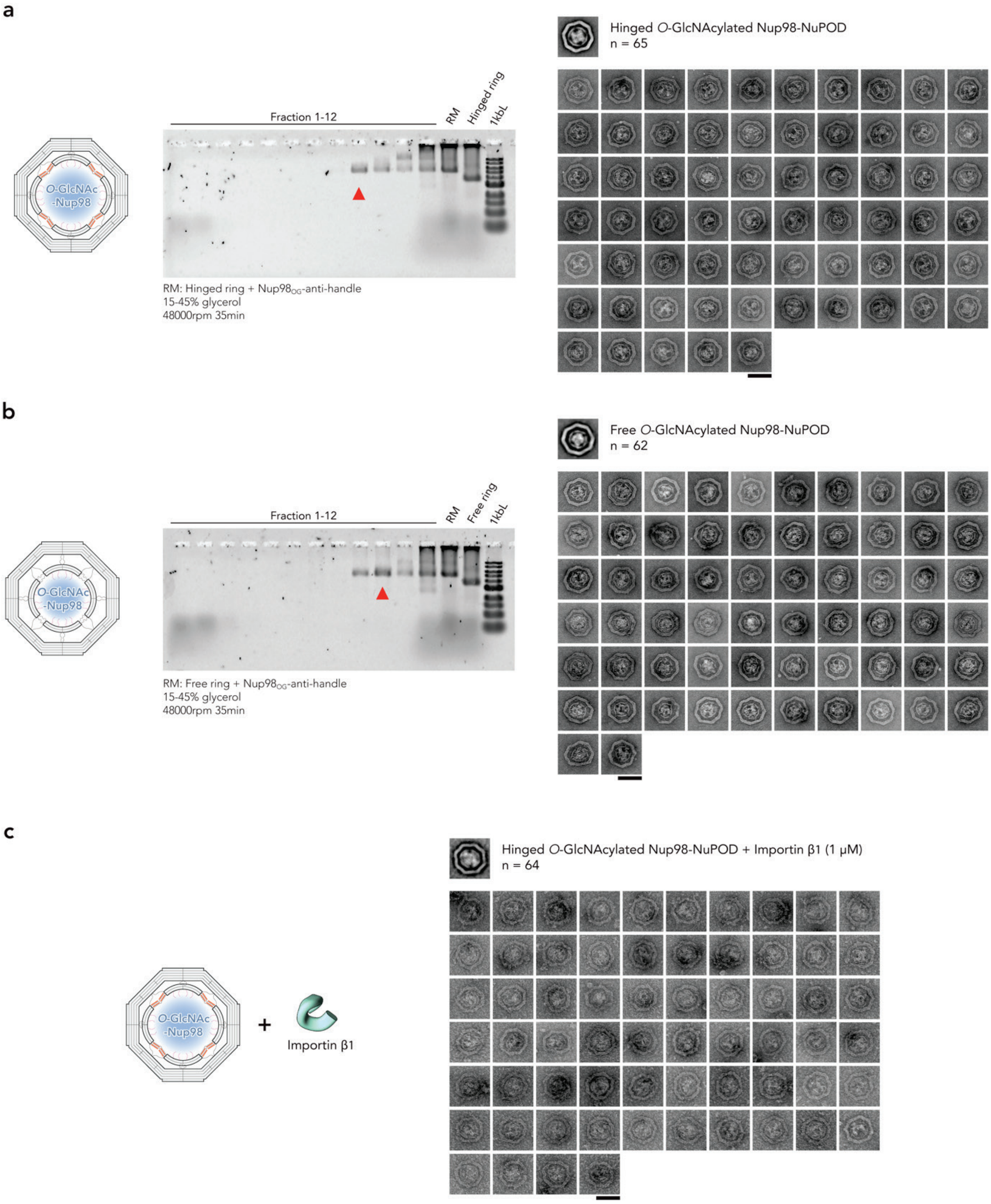


**Fig. S29 | Structural characterization of *O*-GlcNAcylated Nup98-NuPODs with and without importin β1. a**, **b**, Agarose gel electrophoresis analysis of fractions collected after rate-zonal centrifugation (15–45% glycerol, 48,000 r.p.m., 35 min) of Hinged (**a**) and Free (**b**) O-GlcNAcylated Nup98 ($Nup98_{OG}$)-NuPODs. Red arrowheads indicate the fractions selected for negative-stain TEM analyses. Representative particle galleries of the purified $Nup98_{OG}$-NuPODs are shown on the right. **c**, Representative particle gallery of Hinged $Nup98_{OG}$-NuPODs after incubation with 1 μM importin β1. RM, reaction mixture; 1 kbL, 1 kb DNA ladder. Scale bars, 100 nm.

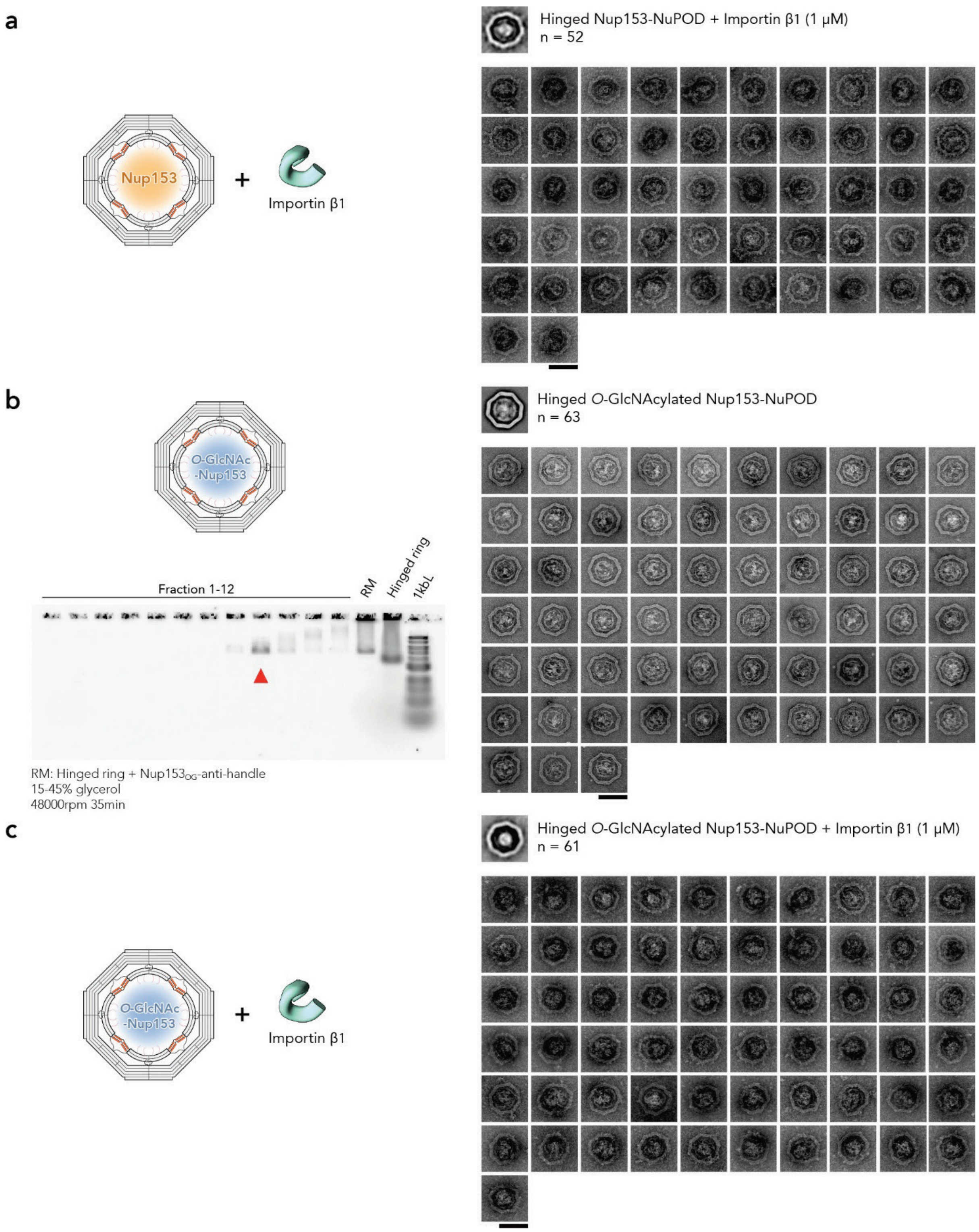


**Fig. S30 | Structural characterization of Nup153-NuPODs with and without *O*-GlcNAcylation and importin β1. a**, Representative particle gallery of Hinged Nup153-NuPODs after incubation with 1 μM importin β1 (Impβ1). **b**, Agarose gel electrophoresis analysis of fractions collected after rate-zonal centrifugation of Hinged *O*-GlcNAcylated Nup153 ($Nup153_{OG}$)-NuPODs. The red arrowhead indicates the fraction selected for negative-stain TEM analysis. A gallery of the purified $Nup153_{OG}$-NuPODs is shown on the right. **c**, Representative particle gallery of Hinged $Nup153_{OG}$-NuPODs after incubation with 1 μM Impβ1. RM, reaction mixture; 1 kbL, 1 kb DNA ladder. Scale bars, 100 nm.

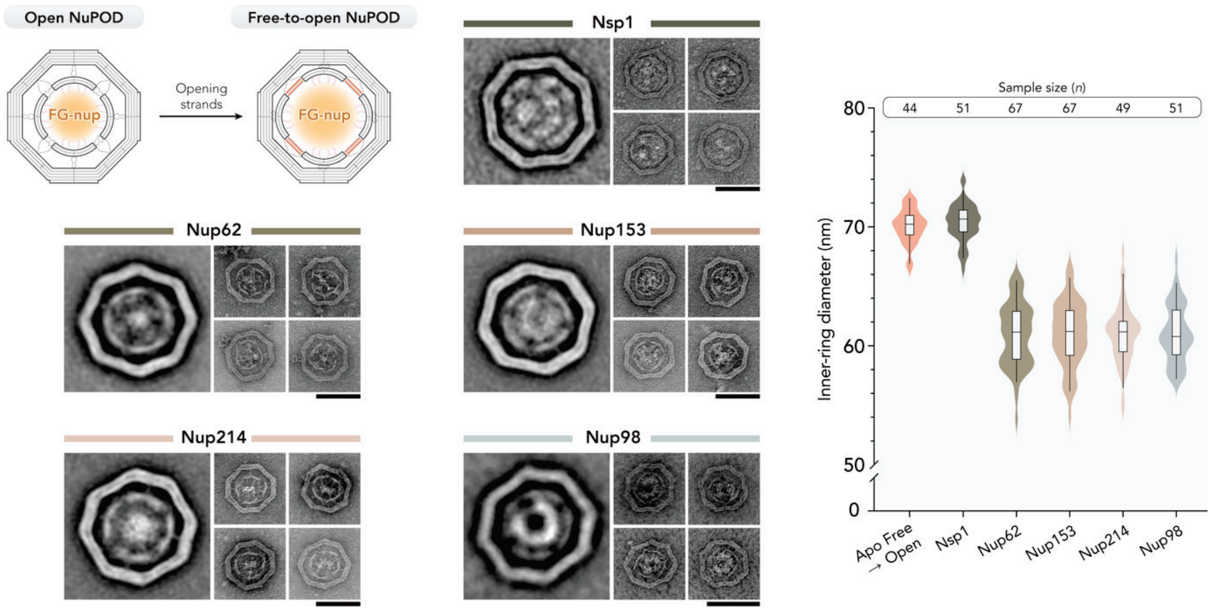


**Fig. S31 | Structural characterization of Free-to-open NuPODs.** Schematic illustration of the conversion of Free NuPODs to Open NuPODs by opening strands. Class averages, representative particle galleries, and $d_{in}$ measurement of Free-to-open Nsp1-, Nup62-, Nup153-, Nup214-, and Nup98-NuPODs are shown. Lines, boxes, and whiskers in violin plots show the median, 25th–75th, and 5th–95th percentiles, respectively. Scale bars, 100 nm.

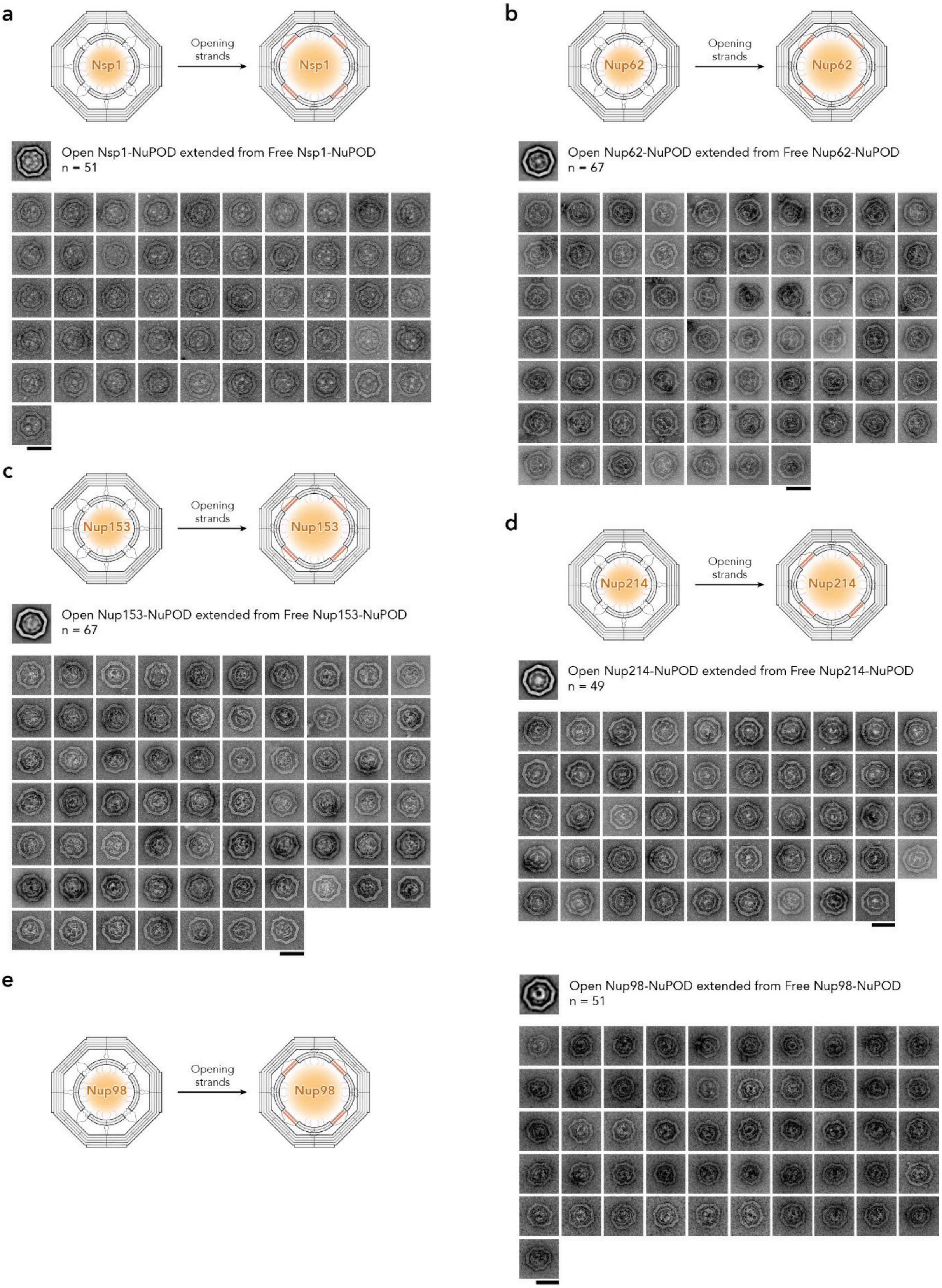


**Fig. S32 | Representative particle galleries of Free-to-open NuPODs. a**–**e**, Representative particle galleries of Open NuPODs converted from Free NuPODs by opening strands. NuPODs were assembled with Nsp1 (**a**), Nup62 (**b**), Nup153 (**c**), Nup214 (**d**), or Nup98 (**e**). Scale bars, 100 nm.

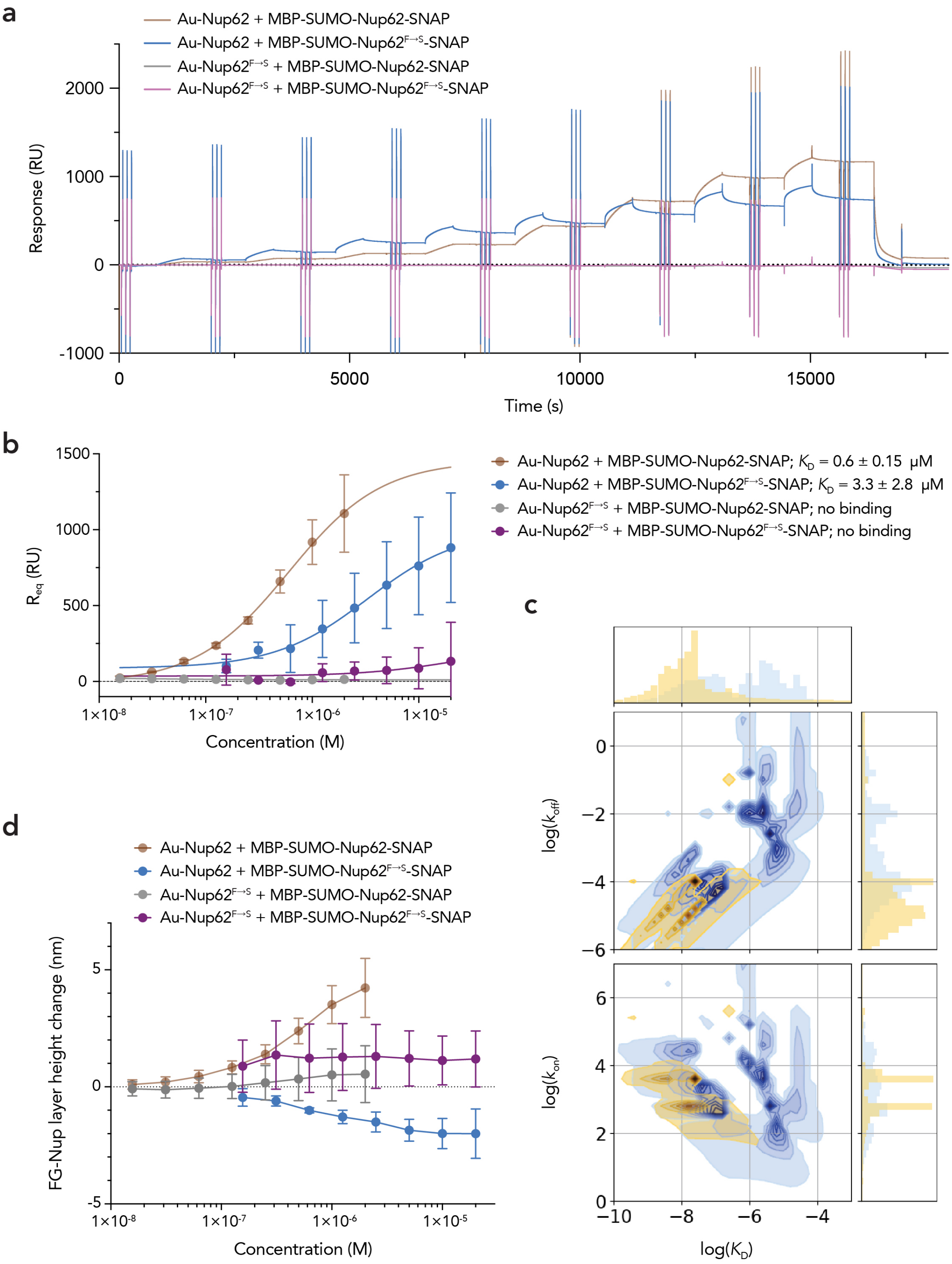


**Fig. S33 | SPR analysis of interaction between Nup62 and Nup62$^{F\to S}$. a**, Representative SPR sensograms showing interactions between immobilized Nup62 or Nup62$^{F\to S}$ and soluble MBP-SUMO-Nup62-SNAP or MBP-SUMO-Nup62$^{F\to S}$-SNAP at increasing analyte concentrations. Cysteine-modified FG domains were immobilized on the gold surface of the SPR. **b**, Langmuir binding isotherms derived from the equilibrium SPR responses. **c**, Distribution of fitted kinetic parameters for the indicated interactions (color code same as in **b**). **d**, Concentration-dependent change in the initial FG-Nup layer height ($d_0$) upon binding of MBP-SUMO-Nup62-SNAP or MBP-SUMO-Nup62$^{F\to S}$-SNAP. The FG-Nup grafting distance ($g$) and $d_0$ were 5.1 ± 0.3 nm and 6.03 ± 2.3 nm, respectively, for the Nup62 layer, and 6.1 ± 0.5 nm and 7.9 ± 1.1 nm, respectively, for the Nup62$^{F\to S}$ layer. Error bars represent mean ± s.e.m.; $n$ = 3 independent experiments.

### SUPPLEMENTARY TABLES

**Table S1 | Summary of inner-ring diameter ($d_{in}$) of DNA rings and NuPODs measured by negative-stain TEM.** Unless otherwise noted, the expandable ring in each configuration contains 48 handles; when included, [NTR] = 1 µM. The $d_{in}$ values are reported as mean±s.d.; unit: nm.

| FG-Nup / DNA ring | Nup-free (Apo) | Nsp1 | Nup62 | Nup153 | Nup214 | Nup98 |
|---|---|---|---|---|---|---|
| **Open** | 73.3±0.9 | 72.1±2.0 | 64.4±2.0 | 67.2±2.2 | 65.8±2.6 | 66.6±2.1 |
| **Hinged** | 69.6±1.4 | 71.0±2.1 | 62.0±2.0 | 63.6±2.3 | 63.3±2.4 | 63.8±1.8 |
| **Hinged (24 handles)** | | / | 64.3±1.9 | 65.1±1.9 | 64.6±2.8 | 63.1±2.0 |
| **Hinged (12 handles)** | | / | 67.6±1.7 | 67.9±1.9 | 69.6±1.7 | 64.8±1.8 |
| **Free** | 61.2±1.3 | 70.7±2.4 | 57.7±2.3 | 56.5±2.4 | 57.4±2.2 | 55.7±2.2 |
| **Open extend from Free** | 70.0±1.4 | 70.3±1.6 | 60.9±2.6 | 61.0±2.7 | 61.0±2.5 | 61.0±2.4 |
| **Closed** | 55.3±1.8 | / | / | / | / | / |

| FG-Nup / DNA ring | Nsp1 (+Kap95) | Nup62 (+Impβ1) | Nup153 (+Impβ1) | Nup98 (+Impβ1) |
|---|---|---|---|---|
| **Hinged** | 68.7±2.1 | 62.6±2.3 | 62.6±2.3 | 63.7±2.0 |
| **Free** | 62.4±4.0 | / | / | / |

| FG-Nup / DNA ring | Nup62$_{OG}$ | Nup153$_{OG}$ | Nup98$_{OG}$ | Nup62$^{F \to S}$ |
|---|---|---|---|---|
| **Hinged** | 67.0±1.5 | 62.2±2.2 | 65.1±2.6 | 68.3±1.9 |
| **Free** | / | / | 59.3±2.8 | / |

| FG-Nup / DNA ring | Nup62$_{OG}$ (+Impβ1) | Nup153$_{OG}$ (+Impβ1) | Nup98$_{OG}$ (+Impβ1) |
|---|---|---|---|
| **Hinged** | 66.0±1.9 | 64.3±2.8 | 65.2±3.0 |

**Table S2 | Amino acid sequences of recombinant Nup constructs.** The Nup sequences are shown in bold. Protein domain architectures are shown in **Fig. 2b**.

**Nsp1$^{FG}$**

HHHHHHHHHHKIEEGKLVIWINGDKGYNGLAEVGKKFEKDTGIKVTVEHPDKLEEKFPQVAATGDGPD
IIFWAHDRFGGYAQSGLLAEITPDKAFQDKLYPFTWDAVRYNGKLIAYPIAVEALSLIYNKDLLPNPP
KTWEEIPALDKELKAKGKSALMFNLQEPYFTWPLIAADGGYAFKYENGKYDIKDVGVDNAGAKAGLTF
LVDLIKNKHMNADTDYSIAEAAFNKGETAMTINGPWAWSNIDTSKVNYGVTVLPTFKGQPSKPFVGVL
SAGINAASPNKELAKEFLENYLLTDEGLEAVNKDKPLGAVALKSYEEELAKDPRIAATMENAQKGEIM
PNIPQMSAFWYAVRTAVINAASGRQTVDEALKDAQTASMSDSEVNQEAKPEVKPEVKPETHINLKVSD
GSSEIFFKIKKTTPLRRLMEAFAKRQGKEMDSLRFLYDGIRIQADQTPEDLDMEDNDIIEAHREQIGG
HMENLYFQG**NFNTPQQNKTPFSFGTANNNSNTTNQNSSTGAGAFGTGQSTFGFNNSAPNNTNNANSSI**
**TPAFGSNNTGNTAFGNSNPTSNVFGSNNSTTNTFGSNSAGTSLFGSSSAQQTKSNGTAGGNTFGSSSL**
**FNNSTNSNTTKPAFGGLNFGGGNNTTPSSTGNANTSNNLFGATANANKPAFSFGATTNDDKKTEPDKP**
**AFSFNSSVGNKTDAQAPTTGFSFGSQLGGNKTVNEAAKPSLSFGSGSAGANPAGASQPEPTTNEPAKP**
**ALSFGTATSDNKTTNTTPSFSFGAKSDENKAGATSKPAFSFGAKPEEKKDDNSSKPAFSFGAKSNEDK**
**QDGTAKPAFSFGAKPAEKNNNETSKPAFSFGAKSDEKKDGDASKPAFSFGAKPDENKASATSKPAFSF**
**GAKPEEKKDDNSSKPAFSFGAKSNEDKQDGTAKPAFSFGAKPAEKNNNETSKPAFSFGAKSDEKKDGD**
**ASKPAFSFGAKSDEKKDSDSSKPAFSFGTKSNEKKDSGSSKPAFSFGAKPDEKKNDEVSKPAFSFGAK**
**ANEKKESDESKSAFSFGSKPTGKEEGDGAKAAISFGAKPEEQKSSDTSKPAFTFGAQKDNEKKTEES**K
LMDKDCEMKRTTLDSPLGKLELSGCEQGLHEIKLLGKGTSAADAVEVPAPAAVLGGPEPLMQATAWLN
AYFHQPEAIEEFPVPALHHPVFQQESFTRQVLWKLLKVVKFGEVISYQQLAALAGNPAATAAVKTALS
GNPVPILIPCHRVVSSSGAVGGYEGGLAVKEWLLAHEGHRLGKPGLG

**Nup62**

HHHHHHHHHHKIEEGKLVIWINGDKGYNGLAEVGKKFEKDTGIKVTVEHPDKLEEKFPQVAATGDGPD
IIFWAHDRFGGYAQSGLLAEITPDKAFQDKLYPFTWDAVRYNGKLIAYPIAVEALSLIYNKDLLPNPP
KTWEEIPALDKELKAKGKSALMFNLQEPYFTWPLIAADGGYAFKYENGKYDIKDVGVDNAGAKAGLTF
LVDLIKNKHMNADTDYSIAEAAFNKGETAMTINGPWAWSNIDTSKVNYGVTVLPTFKGQPSKPFVGVL
SAGINAASPNKELAKEFLENYLLTDEGLEAVNKDKPLGAVALKSYEEELAKDPRIAATMENAQKGEIM
PNIPQMSAFWYAVRTAVINAASGRQTVDEALKDAQTASMSDSEVNQEAKPEVKPEVKPETHINLKVSD
GSSEIFFKIKKTTPLRRLMEAFAKRQGKEMDSLRFLYDGIRIQADQTPEDLDMEDNDIIEAHREQIGG
HMENLYFQG**MSGFNFGGTGAPTGGFTFGTAKTATTTPATGFSFSTSGTGGFNFGAPFQPATSTPSTGL**
**FSLATQTPATQTTGFTFGTATLASGGTGFSLGIGASKLNLSNTAATPAMANPSGFGLGSSNLTNAISS**
**TVTSSQGTAPTGFVFGPSTTSVAPATTSGGFSFTGGSTAQPSGFNIGSAGNSAQPTAPATLPFTPATP**
**AATTAGATQPAAPTPTATITSTGPSLFASIATAPTSSATTGLSLCTPVTTAGAPTAGTQGFSLKAPGA**
**ASGTSTTTSTAATATATTTSSSSTTGFALNLKPLAPAGIPSNTAAAVTAPPGPGAAAGAAASSAMTYA**
**QLESLINKWSLELEDQERHFLQQATQVNAWDRTLIENGEKITSLHREVEKVKLDQKRLDQELDFILSQ**
**QKELEDLLSPLEELVKEQSGTIYLQHADEEREKTYKLAENIDAQLKRMAQDLKDIIEHLNTSGAPADT**
**SDPLQQICKILNAHMDSLQWIDQNSALLQRKVEEVTKVCEGRRKEQERSFRITFD**KLMDKDCEMKRTT
LDSPLGKLELSGCEQGLHEIKLLGKGTSAADAVEVPAPAAVLGGPEPLMQATAWLNAYFHQPEAIEEF
PVPALHHPVFQQESFTRQVLWKLLKVVKFGEVISYQQLAALAGNPAATAAVKTALSGNPVPILIPCHR
VVSSSGAVGGYEGGLAVKEWLLAHEGHRLGKPGLG

**Nup153**$^{FG}$

HHHHHHHHHHMDKDCEMKRTTLDSPLGKLELSGCEQGLHEIKLLGKGTSAADAVEVPAPAAVLGGPEP
LMQATAWLNAYFHQPEAIEEFPVPALHHPVFQQESFTRQVLWKLLKVVKFGEVISYQQLAALAGNPAA
TAAVKTALSGNPVPILIPCHRVVSSSGAVGGYEGGLAVKEWLLAHEGHRLGKPGLGKIEEGKLVIWIN
GDKGYNGLAEVGKKFEKDTGIKVTVEHPDKLEEKFPQVAATGDGPDIIFWAHDRFGGYAQSGLLAEIT
PDKAFQDKLYPFTWDAVRYNGKLIAYPIAVEALSLIYNKDLLPNPPKTWEEIPALDKELKAKGKSALM
FNLQEPYFTWPLIAADGGYAFKYENGKYDIKDVGVDNAGAKAGLTFLVDLIKNKHMNADTDYSIAEAA
FNKGETAMTINGPWAWSNIDTSKVNYGVTVLPTFKGQPSKPFVGVLSAGINAASPNKELAKEFLENYL
LTDEGLEAVNKDKPLGAVALKSYEEELAKDPRIAATMENAQKGEIMPNIPQMSAFWYAVRTAVINAAS
GRQTVDEALKDAQTASMSDSEVNQEAKPEVKPEVKPETHINLKVSDGSSEIFFKIKKTTPLRRLMEAF
AKRQGKEMDSLRFLYDGIRIQADQTPEDLDMEDNDIIEAHREQIGGHMENLYFQG**NSAASSSFKFGVS
SSSSGPSQTLTSTGNFKFGDQGGFKIGVSSDSGSINPMSEGFKFSKPIGDFKFGVSSESKPEEVKKDS
KNDNFKFGLSSGLSNPVSLTPFQFGVSNLGQEEKKEELPKSSSAGFSFGTGVINSTPAPANTIVTSEN
KSSFNLGTIETKSASVAPFTCKTSEAKKEEMPATKGGFSFGNVEPASLPSASVFVLGRTEEKQQEPVT
STSLVFGKKADNEEPKCQPVFSFGNSEQTKDENSSKSTFSFSMTKPSEKESEQPAKATFAFGAQTSTT
ADQGAAKPVFSFLNNSSSSSSTPATSAGGGIFGSSTSSSNPPVATFVFGQSSNPVSSSAFGNTAESST
SQSLLFSQDSKLATTSSTGTAVTPFVFGPGASSNNTTTSGFGFGATTTSSSAGSSFVFGTGPSAPSAS
PAFGANQTPTFGQSQGASQPNPPGFGSISSSTALFPTGSQPAPPTFGTVSSSSQPPVFGQQPSQSAFG
SGTTPNSSSAFQFGSSTTNFNFTNNSPSGVFTFGANSSTPAASAQPSGSGGFPFNQSPAAFTVGSNGK
NVFSSSGTSFSGRKIKTAVRRRK**KL

**Nup214**$^{C\text{-}term\ FG}$

MGKIEEGKLVIWINGDKGYNGLAEVGKKFEKDTGIKVTVEHPDKLEEKFPQVAATGDGPDIIFWAHDR
LGGYAQSGLLAEITPDKAFQDKLYPFTWDAVRYNGKLIAYPIAVEALSLIYNKDLLPNPPKTWEEIPA
LDKELKAKGKSALMFNLQEPYFTWPLIAADGGYAFKYENGKYDIKDVGVDNAGAKAGLTFLVDLIKNK
HMNADTDYSIAEAAFNKGETAMTINGPWAWSNIDTSKVNYGVTVLPTFKGQPSKPFVGVLSAGINAAS
PNKELAKEFLENYLLTDEGLEAVNKDKPLGAVALKSYEEELAKDPRIAATMENAQKGEIMPNIPQMSA
FWYAVRTAVINAASGRQTVDEALKDAQTASMSDSEVNQEAKPEVKPEVKPETHINLKVSDGSSEIFFK
IKKTTPLRRLMEAFAKRQGKEMDSLRFLYDGIRIQADQTPEDLDMEDNDIIEAHREQIGGHMENLYFQ
G**STATSNTSNLFGNSGAKTFGGFASSSFGEQKPTGTFSSGGGSVASQGFGFSSPNKTGGFGAAPVFGS
PPTFGGSPGFGGVPAFGSAPAFTSPLGSTGGKVFGEGTAAASAGGFGFGSSSNTTSFGTLASQNAPTF
GSLSQQTSGFGTQSSGFSGFGSGTGGFSFGSNNSSVQGFGGWRS**MDKDCEMKRTTLDSPLGKLELSGC
EQGLHEIKLLGKGTSAADAVEVPAPAAVLGGPEPLMQATAWLNAYFHQPEAIEEFPVPALHHPVFQQE
SFTRQVLWKLLKVVKFGEVISYQQLAALAGNPAATAAVKTALSGNPVPILIPCHRVVSSSGAVGGYEG
GLAVKEWLLAHEGHRLGKPGLGHHHHHH

**Nup98**$^{FG}$

HHHHHHHHHHKIEEGKLVIWINGDKGYNGLAEVGKKFEKDTGIKVTVEHPDKLEEKFPQVAATGDGPD
IIFWAHDRLGGYAQSGLLAEITPDKAFQDKLYPFTWDAVRYNGKLIAYPIAVEALSLIYNKDLLPNPP
KTWEEIPALDKELKAKGKSALMFNLQEPYFTWPLIAADGGYAFKYENGKYDIKDVGVDNAGAKAGLTF
LVDLIKNKHMNADTDYSIAEAAFNKGETAMTINGPWAWSNIDTSKVNYGVTVLPTFKGQPSKPFVGVL
SAGINAASPNKELAKEFLENYLLTDEGLEAVNKDKPLGAVALKSYEEELAKDPRIAATMENAQKGEIM
PNIPQMSAFWYAVRTAVINAASGRQTVDEALKDAQTASMSDSEVNQEAKPEVKPEVKPETHINLKVSD
GSSEIFFKIKKTTPLRRLMEAFAKRQGKEMDSLRFLYDGIRIQADQTPEDLDMEDNDIIEAHREQIGG

HMENLYFQG**MFNKSFGTPFGGGTGGFGTTSTFGQNTGFGTTSGGAFGTSAFGSSNNTGGLFGNSQTKP GGLFGTSSFSQPATSTSTGFGFGTSTGTANTLFGTASTGTSLFSSQNNAFAQNKPTGFGNFGTSTSSG GLFGTTNTTSNPFGSTSGSLFGPSSFTAAPTGTTIKFNPPTGTDTMVKAGVSTNISTKHQCITAMKEY ESKSLEELRLEDYQANRKGPQNQVGAGTTTGLFGSSPATSSATGLFSSSTTNSGFAYGQNKTAFGTST TGFGTNPGGLFGQQNQQTTSLFSKPFGQATTTQNTGFSFGNTSTIGQPSTNTMGLFGVTQASQPGGLF GTATNTSTGTAFGTGTGLFGQTNTGFGAVGSTLFGNNKLTTFGSSTTSAPSFGTTSGGLFGFGTNTSG NSIFGSKPAPGTLGTGLGAGFGTALGAGQASLFGNNQPKIGGPLGTGAFGAPGFNTTTATLGFGAPQA PVALTDPNASAAQQAVLQQHINSLTYSPFGD**KLMDKDCEMKRTTLDSPLGKLELSGCEQGLHEIKLLG KGTSAADAVEVPAPAAVLGGPEPLMQATAWLNAYFHQPEAIEEFPVPALHHPVFQQESFTRQVLWKLL KVVKFGEVISYQQLAALAGNPAATAAVKTALSGNPVPILIPCHRVVSSSGAVGGYEGGLAVKEWLLAH EGHRLGKPGLG

**Nup62$^{F\rightarrow S}$**

HHHHHHHHHHKIEEGKLVIWINGDKGYNGLAEVGKKFEKDTGIKVTVEHPDKLEEKFPQVAATGDGPD IIFWAHDRLGGYAQSGLLAEITPDKAFQDKLYPFTWDAVRYNGKLIAYPIAVEALSLIYNKDLLPNPP KTWEEIPALDKELKAKGKSALMFNLQEPYFTWPLIAADGGYAFKYENGKYDIKDVGVDNAGAKAGLTF LVDLIKNKHMNADTDYSIAEAAFNKGETAMTINGPWAWSNIDTSKVNYGVTVLPTFKGQPSKPFVGVL SAGINAASPNKELAKEFLENYLLTDEGLEAVNKDKPLGAVALKSYEEELAKDPRIAATMENAQKGEIM PNIPQMSAFWYAVRTAVINAASGRQTVDEALKDAQTASMSDSEVNQEAKPEVKPEVKPETHINLKVSD GSSEIFFKIKKTTPLRRLMEAFAKRQGKEMDSLRFLYDGIRIQADQTPEDLDMEDNDIIEAHREQIGG HMENLYFQG**MSGSNSGGTGAPTGGSTSGTAKTATTTPATGSSSSTSGTGGSNSGAPFQPATSTPSTGL SSLATQTPATQTTGSTSGTATLASGGTGSSLGIGASKLNLSNTAATPAMANPSGSGLGSSNLTNAISS TVTSSQGTAPTGSVSGPSTTSVAPATTSGGSSSTGGSTAQPSGSNIGSAGNSAQPTAPATLPFTPATP AATTAGATQPAAPTPTATITSTGPSLFASIATAPTSSATTGLSLCTPVTTAGAPTAGTQGFSLKAPGA ASGTSTTTSTAATATATTTTSSSTTGFALNLKPLAPAGIPSNTAAAVTAPPGPGAAAGAAASSAMTYA QLESLINKWSLELEDQERHFLQQATQVNAWDRTLIENGEKITSLHREVEKVKLDQKRLDQELDFILSQ QKELEDLLSPLEELVKEQSGTIYLQHADEEREKTYKLAENIDAQLKRMAQDLKDIIEHLNTSGAPADT SDPLQQICKILNAHMDSLQWIDQNSALLQRKVEEVTKVCEGRRKEQERSFRITFD**KLMDKDCEMKRTT LDSPLGKLELSGCEQGLHEIKLLGKGTSAADAVEVPAPAAVLGGPEPLMQATAWLNAYFHQPEAIEEF PVPALHHPVFQQESFTRQVLWKLLKVVKFGEVISYQQLAALAGNPAATAAVKTALSGNPVPILIPCHR VVSSSGAVGGYEGGLAVKEWLLAHEGHRLGKPGLG

**Table S3 | Staple strands used for expandable ring assembly.**

| Oligonucleotide | Sequence (5’→3’) |
|---|---|
| Octagonal frame_1 | AAAAAAACCGTCTATCAGGGAAAGGAATGGTTTTGACCCAGCTAGCTAAACAGGTA |
| Octagonal frame_2 | ACATAAATCGTCTTTTAATGGAAAC |
| Octagonal frame_3 | GCAAATTTTTAGAACCCTCACTGGTGCCAGAA |
| Octagonal frame_4 | ACAGAAAAACATTATGACCATAAA |
| Octagonal frame_5 | GGAGGGCGTTTTATCATATTCCTGATTATC |
| Octagonal frame_6 | TGATTGCTTTGATAAGAGAGAATCAAAGGGAGTCCC |
| Octagonal frame_7 | CATTAATTCTGCGGCCGCCACCATTTTCGGTCA |
| Octagonal frame_8 | TGGGTACGCCAGCTGGCGAAAGGGGGAGGATG |
| Octagonal frame_9 | CGCCGTTACAAAACCGATAGTTGCGCCGACAATGAAGGGACAGAG |
| Octagonal frame_10 | ATAGCTATCTTACCGAGCATCACCTTGTTGCCAGTTACAACAGC |
| Octagonal frame_11 | AGAAGATAAAACACAGGAACGGAGGCCGGGCA |
| Octagonal frame_12 | TAAAAGCCCTTTTTAAGAAAAGTAAGCA |
| Octagonal frame_13 | AACACCGGAACAGTGCCTTG |
| Octagonal frame_14 | TATCATTCCAACGCTAACGGTTATCTAAAATATCACGT |
| Octagonal frame_15 | AATCATATGTCAAATCACCATCA |
| Octagonal frame_16 | TATTAAAGAACGTGGACTCCATTTA |
| Octagonal frame_17 | GAAACCACCAGAAGGAGCGGACCAA |
| Octagonal frame_18 | TTTTCAGCTAATCTGCATCAGACGATAATGGGTAAACTGGTGTGTTCA |
| Octagonal frame_19 | CGGGCGCTAGGGCGCATAAAGTACCTAT |
| Octagonal frame_20 | AAACTTTTAAATTAGTTGCTAACCTCAAGGCAAATCAACA |
| Octagonal frame_21 | TTGAATTGCTGAGCCTAAGTTGGGTAGAAAAT |
| Octagonal frame_22 | CATGAAAATTAATGATGAAATTCAATTATTCAGAGGTGGAGATTTCTTAAACAGCT |
| Octagonal frame_23 | TCATCAGGCAAGACAAAGAACG |
| Octagonal frame_24 | GACCTTCACCGGGCGAAGGGTGAGAAAGGCCG |
| Octagonal frame_25 | AGATATGCAACTGGTCAGTTATATCAAACCCTCAATGGGCCTCTGCTGTAGCGTGT |
| Octagonal frame_26 | TCTGTACATAAAGTTTTTGCGCCATTTGGGAAT |
| Octagonal frame_27 | AACACCACCCTGTAATACTTATTCCATACTGT |
| Octagonal frame_28 | CAGTGCCAAGCTCCTGAGCAAATCCTTTGAGAATAACAATTGGCAAAGAAGT |
| Octagonal frame_29 | TAGCCCTAAAAATAAGAAACGAT |
| Octagonal frame_30 | GTTGCGATGGCCCACTAAAATGCC |
| Octagonal frame_31 | AACATTTATTCATCAAACATCGATTTTAGAGAGGTGAGGCGGTCAAATC |
| Octagonal frame_32 | CAACTTTCACCGTTACCTGGAAGCATAAAGTGTAAAGCCT |
| Octagonal frame_33 | CACAAGAATTGAGTTAAGCCCAATAATAAGAAAGCGTATTAAAAAGTTT |
| Octagonal frame_34 | CGATGTTTTTCTAATAGATATACGAGCCGCAGCCAGCGGTGCCGATG |
| Octagonal frame_35 | GCCTCTGGCCTTCCTGTAGCCAGCTTTCAGC |
| Octagonal frame_36 | CCCCGCAGAACGCGCCTGTTGACAAAAGGAACCCTAGAGCGGGACAAT |
| Octagonal frame_37 | GAAGCCAACAGCTGATTGCCTTGCGGGATGAGGAAGAGCGTCTT |
| Octagonal frame_38 | CACCGTCACCGACTTGATGTGCTGCAAGGCGATACGGAACA |
| Octagonal frame_39 | GAGACAGTACCCCGGTCTGAGAGAGTTGCAGCAA |
| Octagonal frame_40 | CAGCAGGCAAAGCGCCATTCGCCATCGGCAGA |

| | |
|---|---|
| Octagonal frame_41 | ACAGTATATAATTCGCTATAAGGGCGATCGGTGCCAATATCTAAAGTACGTCAA |
| Octagonal frame_42 | AAGAACGCGACACTAACAACTAA |
| Octagonal frame_43 | GCTTGATTCCCAACATCCAAGAGCCGCCGCCAG |
| Octagonal frame_44 | GCGTTTATTTCACATAAAGCAAATTAAGAAATTCTTACCA |
| Octagonal frame_45 | TCCAGAGCCTAATCTGATTTTGCAAGCCTTAAGTAAAGCACAAG |
| Octagonal frame_46 | TGAGTAATGTGTAGGTAAAGACAG |
| Octagonal frame_47 | GATGCAAATCCAATCTAGGGCTTAATGAAA |
| Octagonal frame_48 | CAGTCACACGACCAGTAATAAACAACAACCCGCC |
| Octagonal frame_49 | GATAGCCCTGAGAGCCAGCAGCAAATGAAAGTATTAACACCGCCTGCA |
| Octagonal frame_50 | ATAACGGATTATCGCCCACG |
| Octagonal frame_51 | CCGGAGTGAGACGGGCAAATCTAAATCGGTAA |
| Octagonal frame_52 | CGTATAACGTGCTTAATCAGTGAGGCCA |
| Octagonal frame_53 | TTATTCCTCGTTAATTGGCAAGTGTAGCGG |
| Octagonal frame_54 | TCCAACATCGCCAAGAATACGTGGCACAGA |
| Octagonal frame_55 | TTCTTGCGGGGGAAACCTTTCCGGCACCGCTTTATATTTTATTC |
| Octagonal frame_56 | CCACACAACAAGTCCTGAAC |
| Octagonal frame_57 | AAGGATACGTGAACCATCACATTG |
| Octagonal frame_58 | GAGGATTCTGTCCAGACGACGACAATACCAATAAAGGAGGTGCCATCA |
| Octagonal frame_59 | TTTACGAACCACAATAAACAAAACAATGAAATAGCA |
| Octagonal frame_60 | AGCGTCATACATGGCTTTTGAAACG |
| Octagonal frame_61 | CTTAATTACCTTGCTATTAAAGCGATAGAATCCCCCAAAACGACGGC |
| Octagonal frame_62 | TGGGCGCCAGGGTGTTAGAGCTTGACGG |
| Octagonal frame_63 | CAAAGCCTGGCCTGATAATCAGAAAAGCCCCAAAAATTC |
| Octagonal frame_64 | TGATATCGCGCAGAGGCGAAGTAATTTAATTAAAGGAAGA |
| Octagonal frame_65 | AAGTTACCGAATCCTGAATCTTAATCAGCGAACCCCCC |
| Octagonal frame_66 | TTACGCAGGAGGAAACGCAA |
| Octagonal frame_67 | CTGGAATTAGCATAAATCGGTTTAGTATTGCTCGTCCTGGTAATAAGTTTTTGAT |
| Octagonal frame_68 | ATATTCATTGCTTAGATTAA |
| Octagonal frame_69 | ACCATTAAATGTGAGCGAGT |
| Octagonal frame_70 | GCAAAGCGCCATCAAAAATAAAAT |
| Octagonal frame_71 | CAGTATCGGCCTCAGGAAGATCGCACTCCATC |
| Octagonal frame_72 | ACATTTAATTTTGACGTTGTTCAAATGCTTTAAACAGTTCA |
| Octagonal frame_73 | GTAGCGCGTTTTCATTCAGGCTGCGCAATAACAGTTAGAG |
| Octagonal frame_74 | GGGTTCATAATTACTAGAAAACGCTCAAACAGGCAATCAT |
| Octagonal frame_75 | TATAATCGTTAACGGCATCAGGTG |
| Octagonal frame_76 | TCTTCATATGCGTTATAAACAACATGTTTTGGGGTCCCTCAGAGACGC |
| Octagonal frame_77 | TCACGCTGCGCGTAACCACCACACCCCAGTACCTTCAC |
| Octagonal frame_78 | TGGACGCTGGTTGTAAAACTAGCATGTC |
| Octagonal frame_79 | AGCTATTTTTGAAGAGTCTGTGGTTCCGGCCAGCTGGCAAGCCGCCAA |
| Octagonal frame_80 | TGAGGCTTGCATCGTCTGATTAACCCTCAAACTGT |
| Octagonal frame_81 | AAGAAAAATAGCTCACAATT |
| Octagonal frame_82 | AATATTGAGTCTTTAATGCGTAAAAGAGGCTTTGACGAGCA |
| Octagonal frame_83 | CCGAGCGAACTGAAGATGATGTTGCGGAACAAA |

| | |
|---|---|
| Octagonal frame_84 | TGAGGCCAGCCAACATTTTGACGCTCAAGGGA |
| Octagonal frame_85 | CAATATTTAGAGAGATAACC |
| Octagonal frame_86 | CTGTGTGAAATAACTCACACAACGCGCCAGGCGAAAAAT |
| Octagonal frame_87 | ACAACATTAATGATGTAGAACATAGCTGTTTC |
| Octagonal frame_88 | TAGCGTAGCGCCGCTACAGACTCATCAGCA |
| Octagonal frame_89 | GGAAAGGTATTCTGCGGTCCAACAAGAGTCCAC |
| Octagonal frame_90 | GTTAATGAATATACAGTAAGCCGCGCTTAATAAAGGGTATTA |
| Octagonal frame_91 | GCAAAAATGGATGTAATATCCAGAACACAAAGTCTAC |
| Octagonal frame_92 | TTTTTGTTTAACGTCAAAAATTATCATTGCAA |
| Octagonal frame_93 | ATATGATATTCAACCGTTCTAGCTGATACGGTAATCTGCCCCAGGGGG |
| Octagonal frame_94 | TATACTATGGTTTCTGTCCATTTACATCATATTCGGTCGC |
| Octagonal frame_95 | TTCAGCTTATCCGCCGGCGAGCGGTTTGCGTAT |
| Octagonal frame_96 | AGAGACGTGGCGAGAAACGCGGCGCGTAATCC |
| Octagonal frame_97 | AATTTACGGAAGGGAAGAAAGCGAAAGGAG |
| Octagonal frame_98 | AACCAAGTACGAGCAATCGGCTTAATTGAAGAGAATCGA |
| Octagonal frame_99 | CTAATATCTTGAATGGCTATTCTGTATTTACATTGGCAGATTCAC |
| Octagonal frame_100 | GGGGTGCCTAATGAGTGAGCTTGTTATCCATATCCCATCCT |
| Octagonal frame_101 | TAGATTAGAGCCGTCAATAGAGTT |
| Octagonal frame_102 | CATAACCGATGGGAGAAACA |
| Octagonal frame_103 | GCGATTCATCTTCCTCCGGCTTAGGTTGTTTATCAATTCAAATACAGG |
| Octagonal frame_104 | CAGGACCAGACCCGAAAGACAATCATAGGTTTAGACTGGA |
| Octagonal frame_105 | CATTGACAGGAGGTACTAATAGTAGTAG |
| Octagonal frame_106 | GACGCGCAGTACAAACGGCGGATTGGGGCGCATCAAA |
| Octagonal frame_107 | CATTATAAAAGACATGATTAAGACTCCTTA |
| Octagonal frame_108 | TAGCGTCCAATCAAAAATTCGCGTTGACA |
| Octagonal frame_109 | CAAATTAGTGAGTGAATAACCTTGCT |
| Octagonal frame_110 | AGTACATAAATCAATATATATTTAAC |
| Octagonal frame_111 | GCCAGTTTCGGATTCTAGAGCCACCACCCTCAGA |
| Octagonal frame_112 | GCCTTTTAGTTTCCTTTTGATAAGAGGTCA |
| Octagonal frame_113 | CGGAAGAGTCAATAGTGAAGGTTATACGAGCTTGGTGGCAGATA |
| Octagonal frame_114 | ACCGACCGTGTGTAGTTAATGCTGAAAACAAAGCGAATTAGAGACGGA |
| Octagonal frame_115 | TATATTTTCAAATATATTTATAAATAGTCA |
| Octagonal frame_116 | AACGAGTAGATTAGCGTCAGACT |
| Octagonal frame_117 | AACAACCCGTGAGGGGACGACGA |
| Octagonal frame_118 | GTTAATGCCCCAATGGAAAGATTGGCCCCAC |
| Octagonal frame_119 | TCTTTTTGTCACAATCAATAGAAATGGAACG |
| Octagonal frame_120 | GTGGCAACATAAAGGTGAATTAT |
| Octagonal frame_121 | CGAGAAAACTTGTAAATGCT |
| Octagonal frame_122 | TCAACCATCTTTCCACCCTCCCGTGGGACTCTGAATTTACCGTTCCAGTA |
| Octagonal frame_123 | GACGCTGAGAATCGTCATAA |
| Octagonal frame_124 | AGTAACAGTGAATAAGAATA |
| Octagonal frame_125 | GAAAACGAGAATGACCATAAATACTG |
| Octagonal frame_126 | ATCAAACCCGCGTGCTATACTCGAACGGGTCGGGGCCCGGAAACTGAG |
| Octagonal frame_127 | ATTGGGGGTTGATTCAAGTATATAGTAGAAGGCGAGAA |

| | |
|---|---|
| Octagonal frame_128 | CATAGAACCGTCATAATCGTAACCAAGACTTGTTT |
| Octagonal frame_129 | ACTCATTGGCGTATGAGGATTAAA |
| Octagonal frame_130 | GTAGAAGGATTGCACAATTCCCTGTCCGATCACCAGTATT |
| Octagonal frame_131 | TTGGTAATTTTTTTAGAAAATTGAAATAATCAACTCGCTCATACGCTCGGGT |
| Octagonal frame_132 | AGGTATGAATTCTTTTTTGAGAATAACAACGTTATATTATACTCGCCCAATGAGA |
| Octagonal frame_133 | CAGATTTGAGGAAAAAGTTTTTGTTTTCTTATATTTTATAATTGTCTATCACATTGAGCG |
| Octagonal frame_134 | CTTTACTTAAATAAAGGGACTGAG |
| Octagonal frame_135 | TGATTATCGGCCGCAGCCTTAGAGGGTAATTTGTGACATT |
| Octagonal frame_136 | TAGAGCCAGCAAAGAGCCTTAATTC |
| Octagonal frame_137 | ATAAGGAAGGTATACCAGCGCATAGCCCAAT |
| Octagonal frame_138 | CCCTATTATCTGTACAATACGTTGTTCCATAATACATATAGAAGTCAA |
| Octagonal frame_139 | CCAGTACCTTTTACCATTAGCGACTTTTTTCCAAAGTTTACATATTAACCAGCTGTGCATCT |
| Octagonal frame_140 | TGAAAATTAATGGGTTGAGTATCTGTTATTAATCTGGATAAACAAGGCCTTAGAGTAACATGAA |
| Octagonal frame_141 | AGCAATCCTGGAGATAGCCGGAGATTTACCCGAAT |
| Octagonal frame_142 | CCTCTTCGCAAAGTAATCAGTAGCAAGGGAGATCAACATATTGTCAAA |
| Octagonal frame_143 | AGGGGTATTAGAAATAGCCCTTTGATGGGAGCAAACCGTTGCGCTCAC |
| Octagonal frame_144 | TCGTTCGTAGAAATAATCAAACCGTACCCCCAAAAGAACATTCATATGGTTAAT |
| Octagonal frame_145 | AAAGATTAAAACTGAAATGTTTAA |
| Octagonal frame_146 | TAATAACGGAATAGATCAGTAGTTA |
| Octagonal frame_147 | ATTATCAATTCTTGAGGCAGAGGCGTTACCCGTATAAACA |
| Octagonal frame_148 | AGATTCTCCATTTCGTGATAACGGAAATTAGCACCATAATTGCTGACC |
| Octagonal frame_149 | AGAGGCTTACCCCTGACTCGTAAGTTAATG |
| Octagonal frame_150 | AGCATCACTTGGTCATAGTTCCGGAACCGCAGCACCTGGTCAATAGGT |
| Octagonal frame_151 | TAGCCCCCTTATTCGGCGTATCAAA |
| Octagonal frame_152 | TACCCTTAAACTAACACAGGAATTCTTAAGCGTTTGGTTT |
| Octagonal frame_153 | GTATAAAGCCAAAGCCTGTTGTACCGAGTGTAATAAACATTTTTGTTATTTTT |
| Octagonal frame_154 | TTTTTAGATAGAACCCTTCTGACCTGAGCAAGGCCATATTATT |
| Octagonal frame_155 | AAGTTACCAGAAGGAAACCTATGTTAGCAAACGTAACGCCAGGGTTTTCCTTTTT |
| Octagonal frame_156 | TTTTTAATCAGCTCATTTTTTCATTAAATCCCTTACAGGTT |
| Octagonal frame_157 | TTTTTCAGTCACCCCTTAGAAAGAAGATTACATTTCGCCATATTTAACAACGCC |
| Octagonal frame_158 | ATTCGAACCAATAGGAACAAATATTTAAATTGTTTTT |
| Octagonal frame_159 | TTTTTTAAACGTTAATATTTTGTTAAA |
| Octagonal frame_160 | AGTACATTTTCGAGCCAGTAAATACCAAAGAATCCTTTCTGGCCAACAGTTTTT |
| Inner-ring arc_1 | CAGCAACCGCAACTAAAACGGTAGAACGTTTTT |
| Inner-ring arc_2 | TTTTTGCCATGTTTACCAGTCGAAAGTTAGCGTAACGATCT |
| Inner-ring arc_3 | GACAAGAACCGGATATTCATTAATAAGTGCCGTCGACAGGCGCTTTCGCACTCAATTTTTT |
| Inner-ring arc_4 | TGACCAACTCAGCGTGGTGCTGGGA |
| Inner-ring arc_5 | TTTTTCGAACTAACGGAACATCTACGTTAGGG |
| Inner-ring arc_6 | ACGCCAAAAGCCGGAATTTGTG |

| | |
|---|---|
| Inner-ring arc_7 | GCGGGGTCAGGGCTTGTTTTT |
| Inner-ring arc_8 | TTTTTCTCATTTTCAATAAAAAGATGGTTTAATTTCAGTAAATTTTGGAGGGTTGAT |
| Inner-ring arc_9 | CAGCTAAAGTTATAAATTTCTGCTCATTTGCCGCTGAA |
| Inner-ring arc_10 | TTTTTAACGATGCTGATTGGAA |
| Inner-ring arc_11 | ATAGCCCTCAGAGCCACCACCTTTTT |
| Inner-ring arc_12 | GGGGGAGTGAGTCCAAAAAAAAGGCTTTTT |
| Inner-ring arc_13 | CTTAAAGAGGCCTACGAAGGCACCAACGTTT |
| Inner-ring arc_14 | CGCGACCCCAGAGCTTTGACCCCAGTTT |
| Inner-ring arc_15 | TCATCAGTATACCATAACACTCCAC |
| Inner-ring arc_16 | AGAAAAAACATTATAGCCCTCATTTACGAGGCATAG |
| Inner-ring arc_17 | TAAGGGAACCGAACATGCGGCGTCAGCAGC |
| Inner-ring arc_18 | TTAGTGATCCGTTCCGTAATGCCAAAAAGAATACA |
| Inner-ring arc_19 | CTTATGCTTGCCCTTGTATCAGGGGTTTTGC |
| Inner-ring arc_20 | AACGAGTAACTTTAGAACCGCCACAAGCCCAA |
| Inner-ring arc_21 | CTAAAATTTGTATCATCGCCTGGGGCCGTTT |

| | |
|---|---|
| Sticky end _1 | TGATCGTATTAAATCCGGCAATCTATCTAG |
| Sticky end _2 | AAAGGAATACCACATTCAACATCATAACAAC |
| Sticky end _3 | ATCAGGTCATTGCCTGGAGATCTACAAAGG |
| Sticky end _4 | GCTCATGGAAATACCTTTGCAACAGGAAAA |
| Sticky end _5 | GAACCAGAAATAAAGAAATTGCGTATAC |
| Sticky end _6 | TGCCATCATGGTACCAATCAATAATCGGCTG |
| Sticky end _7 | GGAGAGCGCCACTACGTGAACTCTGGAACT |
| Sticky end _8 | GAACACCCTGAAATATTACCTAGAAGAAGTTG |
| Sticky end _9 | ATTCAACCGATGTCACCAAACTCCAACAACCTGTTTAGCTATC |
| Sticky end _10 | TCTTAATGGTTTGTTCGGAACCTATTTC |
| Sticky end _11 | ATATTTTCATTTGCACAAACAAAT |
| Sticky end _12 | ACGTCAGAAGCAAAGTATTACCCTGACTATTATA |
| Sticky end _13 | CTGACTGGCTCATTTGAGATTACAAACTACCCTCGTTTACCAGA |
| Sticky end _14 | GAATTCGTACGCTTTCCAGTCGAAA |
| Sticky end _15 | TTTTTCGAAAAAAGTGCTTGAGGACTAAAGACTTCAG |
| Sticky end _16 | CAGACTTAGCCGGAACGAGGCGTTT |
| Sticky end _17 | AAAGAACCGCCTCCCTGGGATAGGTCCTCATTAAAGCCAGCTGCCTATAAAT |
| Sticky end _18 | ACGGGAGAATTAGAATTTCTACGTG |
| Sticky end _19 | TTAAAAATCCCGTATCTCGTCGCATGAGGACAGCGATTATAGGT |
| Sticky end _20 | GGAAACCTGTCGTAAATCGGCGTAGGAATACAATTCGACAAC |
| Sticky end _21 | CTACCATATCAAAATTATGGAAGGGTTAGAAC |
| Sticky end _22 | GGAGGTTCGGATTGCATCAAAAAGA |
| Sticky end _23 | GATTTTCAGGTTTTTTGATTAG |
| Sticky end _24 | AGAGCAGCCTCCGGTGCTCCATAGCGCGAAAGCACGCGTGCCTG |
| Sticky end _25 | ACTAAACAAAGCACGTTTCATAGC |
| Sticky end _26 | CTTCACGTTGGTGTAGATACCGTAATCAGAGCCGTTGATATTGGGC |
| Sticky end _27 | TGTGTACATCGACATCGGCTACAGAGGCTGAATAAT |
| Sticky end _28 | ATAACATCACTAGGGAAGCGCATT |

| | |
|---|---|
| Sticky end _29 | TGGAGGGGGTAATAGTAAAATGTCTGAGAGAC |
| Sticky end _30 | GTTTGCAGTTAATGCTTTATTAAT |
| Sticky end _31 | TGACTTGGTAAGGTTCTACTATGCTTTTTGCCCGATAAAAACTGCC |
| Sticky end _32 | GTTTGAATAAGGCGATTTTATTAGTACCGGAGTTTCGTCACTTT |
| Sticky end _33 | ATCCCTTATAAGTCAAGAATGTAAA |
| Sticky end _34 | TTTGGAAAGTAGAGCCACACTCTC |
| Sticky end _35 | ACCGTCTGAATAATTTGCACAATACTTCAACGTCAGAAGGCCGCTTT |
| Sticky end _36 | CAAAGCTGCTCATTAGGATTAGCCCGTACTCAGGACCA |
| Sticky end _37 | TATTAAGAGGAAGCCGGAAGCAACTTTTTAACTGACCTAAAT |
| Sticky end _38 | AACCATCGATAAGAGCCACCACCG |
| Sticky end _39 | CTATATAAGTCCCCAATAGATAGATTATCAAAAGCTTTACAACATT |
| Sticky end _40 | TTTCCTTATCATTTTTTTATTTTC |
| Sticky end _41 | TGAAATTAGTGACCAAAGACAAAAGGGCGACAG |
| Mini scaffold_1 | TTAAACATTTCACAGATAATAGGGATTCGGGTAAAGAGCGTATTATTGGGGACTTATATAGGC |
| Mini scaffold_2 | TACATTCTTGACAATCTATCGAGCGAGTTGATTATTAACAGATGTATTGTAGTTTTAATCTTT |
| Mini scaffold_3 | CTCAGTCCCTTTTGTTTATCCAGATTAATTCAATTCCCTCATTTAGAACCTTACCAAGTCAAT |
| Mini scaffold_4 | AGATAGATTGCCAAGCATAGAACTTACGAGTCAGGGAAAACAATAAGGCCTATTTAAGTAAAG |
| Mini scaffold_5 | TTTAATCCTCATCAATTATAAAATATAAGGTAAGCCAAAAAAGCACGTAGTGGCGCTCTCCGA |
| Mini scaffold_6 | CGTAGAAATTCTCCAGAGTTAATTCATACCTAATGTCACAAATGTGATAGAACGCCAATGAGT |
| Mini scaffold_7 | TATGAAACGTGCCCAAGTGATGCTAAACAAGTCTTAGCTGGTTCCTGTGTTAGTTTAAGGGTA |
| Mini scaffold_8 | TTTGATACGCCGTAAGAATTAATATGTAAACTTTGCGCGGGTTAACTATGATTTGTTTAGTTT |
| Mini scaffold_9 | TAATAAAGCATTCCGTTCGAGTATAGCAGAAAAAACGCCTTCTGAATTGTGCAATCCTTCTAC |
| Mini scaffold_10 | GAATTAAGGCTCCGGACAGGACTATATACTTGAATTTGATCTCGCCCCGACAACTGCAAACCT |
| Mini scaffold_11 | GAGTGTGGCTCTTTCATTTGACAATATGCAACCCCTATCACGATTGATTATTTCTACGAACGA |
| Mini scaffold_12 | TAACTACTGATCGGTACGGTAATGGAGAATCTATTGGGCTATGTCACTAATACTTTCCAAACA |
| Staple for handle placement_1 | CCATTAAACGGGAAAGGAATCCGCACAGTGGGCGGT |
| Staple for handle placement_2 | TCAGTACCAGGCGGCCCAAATCAACACCAG |
| Staple for handle placement_3 | AATCAATAGAAAGGAACAACTTAAAATACGGCAAACGATAACGG |
| Staple for handle placement_4 | CTGCCACAAAGTACAACGGAGACACTCATCACATCCTCCGGTCCGT |
| Staple for handle placement_5 | TAGGAACCCCACAGACTACAGGTAGATACATA |

| | |
|---|---|
| Staple for handle placement_6 | TAAGAGCAACACTTAATGCAGAAAGAT |
| Staple for handle placement_7 | GCCTGTAGCATTCATGTACCGGTCAGGAGAATTAC |
| Staple for handle placement_8 | CCTCAGAACCGCCGGAATAGGGACGAGAAACGTAA |
| Staple for handle placement_9 | AATTTTTTCACGTCAGCAGTGCGGCCT |
| Staple for handle placement_10 | AACGTGCCGTCTGATAAATTGTGTCAATCA |
| Staple for handle placement_11 | TCACGGTCATACCGGGGGCAGACGGTCGAAATC |
| Staple for handle placement_12 | ATAAGTATAGCCCACCCTCAATCATTGTCGTTGGGA |
| Hinged staple_1 | TTGAAAGAGGACAGATGAACGGTGTACAGACCA |
| Hinged staple_2 | GAGATAGACTTTCTCCGTGGTGAAGGGATAGCT |
| Hinged staple_3 | CGGAAAAAGAGACGCAGAAACAGCGGATCAAAC |
| Hinged staple_4 | AAGTTTTGTCGTCTTTCCAGACGTTAGTAAATG |
| Hinged staple_5 | GCAACCAGCTTACGGCTGGAGGTGTCCAGCATC |
| Hinged staple_6 | CATAGGCTGGCTGACCTTCATCAAGAGTAATCT |
| Hinged staple_7 | TTCTGTATGGGATTTTGCTAAACAACTTTCAAC |
| Hinged staple_8 | AATGCCAACGGCAGCACCGTCGGTGGTGCCATC |
| Open staple_1 | TTGAAAGAGGACAGATGAACGGTGTACAGACCAGGCGCATAGGCTGGCTGACCTTCA TCAAGAGTAATCTT |
| Open staple_2 | GAGATAGACTTTCTCCGTGGTGAAGGGATAGCTCTCACGGAAAAAGAGACGCAGAAA CAGCGGATCAAACT |
| Open staple_3 | AAAGTTTTGTCGTCTTTCCAGACGTTAGTAAATGAATTTTCTGTATGGGATTTTGCT AAACAACTTTCAAC |
| Open staple_4 | GAATGCCAACGGCAGCACCGTCGGTGGTGCCATCCCACGCAACCAGCTTACGGCTGG AGGTGTCCAGCATC |
| Closed staple_1 | TTCTGTATGGGATTTTGCTAAACAACTTTCAACAAGTTTTGTCGTCTTTCCAGACGT TAGTAAATG |
| Closed staple_2 | CATAGGCTGGCTGACCTTCATCAAGAGTAATCTTTGAAAGAGGACAGATGAACGGTG TACAGACCA |
| Closed staple_3 | CGGAAAAAGAGACGCAGAAACAGCGGATCAAACGAGATAGACTTTCTCCGTGGTGAA GGGATAGCT |
| Closed staple_4 | GCAACCAGCTTACGGCTGGAGGTGTCCAGCATCAATGCCAACGGCAGCACCGTCGGT GGTGCCATC |

**Table S4 | DNA sequences used for Nup tethering.** The organization of these oligonucleotides into the different DNA tether architectures is illustrated in **Fig. S8.**

| **Oligonucleotide** | **Sequence (5'→3')** |
|---|---|
| Handle H1 | CTACCATCTCTCCTAAACTCA |
| Anti-handle H1 | TGAGTTTAGGAGAGATGGTAG |
| Handle H2 | AAATTATCTACCACAACTCAC |
| Anti-handle H2 | GTGAGTTGTGGTAGATAATTT |
| Handle H3 | CTATCACCTTCTAACAATTCAACCTACTAACA |
| Anti-handle H3 | TGTTAGTAGGTTGAATTGTTAGAAGGTGATAG |
| Extender C | CTTAACAATCAAAATTATCTACCACAACTCAC |
| Extender (H1+C)' | GTGAGTTGTGGTAGATAATTTTGATTGTTAAGTGAGTTTAGGAGAGATGGTAG |
| Extender (H3+C)' | GTGAGTTGTGGTAGATAATTTTGATTGTTAAGTGTTAGTAGGTTGAATTGTTAGAAGGTGATAG |

### SUPPLEMENTARY MOVIES

**Movies S1–S5 | Simulation movies of Hinged NuPODs.** Simulations movies of Hinged Nsp1- (**S1**), Nup62- (**S2**), Nup153- (**S3**), Nup214- (**S4**), and Nup98- (**S5**) NuPODs, showing the inner ring in top and side views, with a cutaway in the side view.
